# Protocol for calcium imaging of acute brain slices from *Octopus vulgaris* hatchlings during application of neurotransmitters

**DOI:** 10.64898/2026.03.16.711860

**Authors:** Amy Courtney, Marie Van Dijck, Ruth Styfhals, Eduardo Almansa, Horst A. Obenhaus, William R. Schafer, Eve Seuntjens

## Abstract

*Octopus vulgaris* and other cephalopods are of increasing interest as neurobiological model organisms. This protocol describes a method to record calcium activity from individual cells in acute brain slices from *Octopus vulgaris* hatchlings during exogenous application of neurotransmitters. Using this protocol, we characterized single-cell responses to specific neurotransmitters in the optic lobes, which process visual information. The approach is readily adaptable to other cephalopods and small invertebrate species.

**Graphical abstract:** 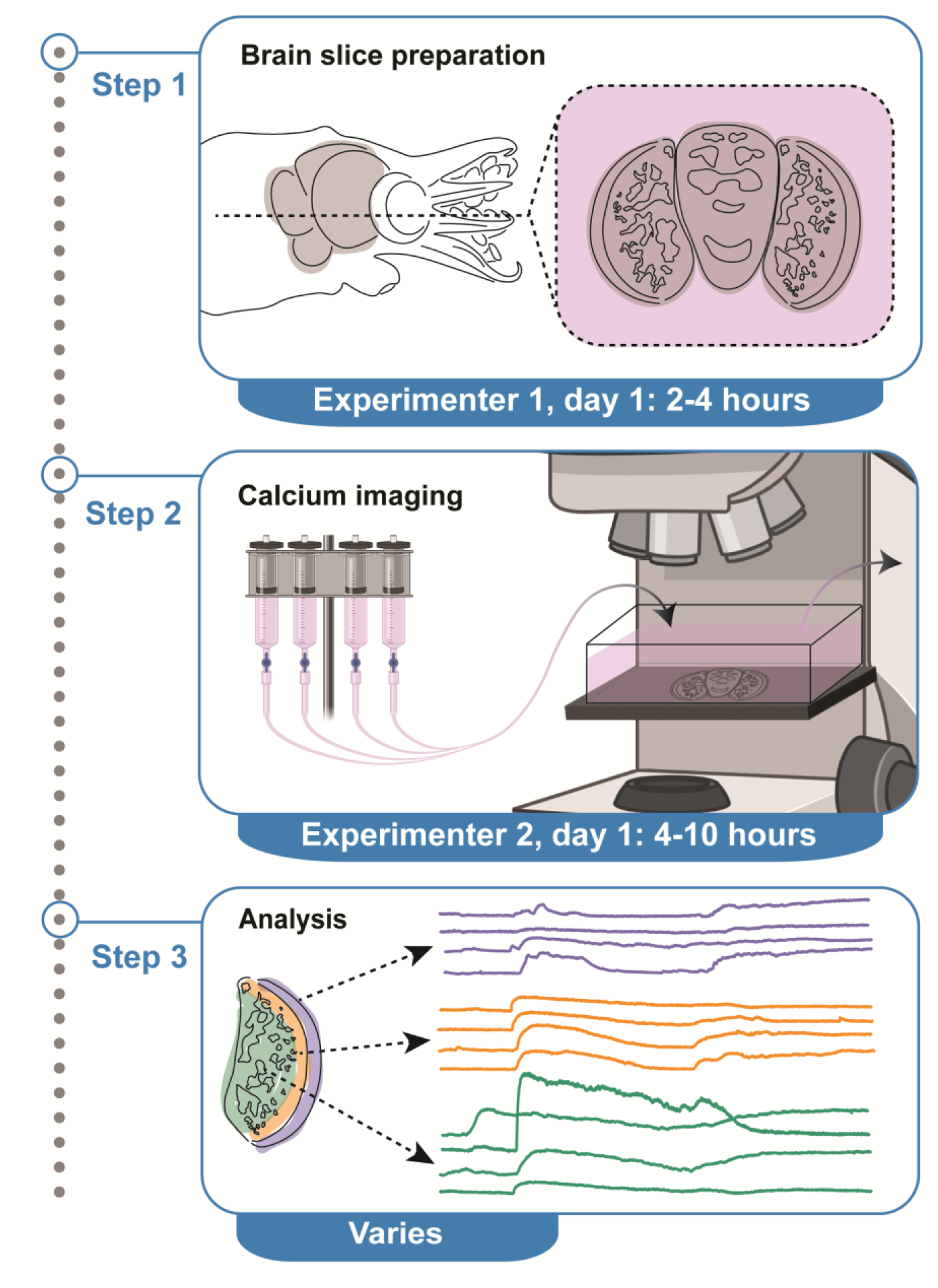

## Before you begin

Cephalopods (octopus, cuttlefish and squid) diverged ∼600 million years ago from mammals, yet they possess a complex brain and display sophisticated visually-driven behaviors (1,2). This independent evolution of brain complexity offers a rare comparative window into molecular and circuit mechanisms of complex brain function. Despite their biological interest, the molecular and neurochemical foundations of cephalopod behavior are largely unknown. With the recent advent of high quality genomic and transcriptomic datasets from multiple cephalopod species (3–12), researchers are increasingly able to probe the intricacies of synaptic transmission and the functional logic of microcircuits. To answer these questions, the development of new tools, such as neuronal calcium imaging, is necessary.

In a previous study, we identified and functionally characterized an excitatory dopamine-gated channel and an inhibitory acetylcholine-gated channel from *Octopus vulgaris*, and mapped their expression within different layers of the optic lobe (13). This led us to hypothesize that dopamine and acetylcholine play excitatory and inhibitory roles, respectively, in the optic lobe. To test this hypothesis, we established a calcium imaging protocol which would allow us to monitor the activity of individual cells in the optic lobe while applying neurotransmitters. While this is not the first time calcium imaging has been used to functionally characterize cells in the optic lobes of cephalopods (14,15), or the first time acute brain slices have been prepared from cephalopod brains (16–18), our approach is the first time both have been combined to enable neurotransmitter mediated single cell activity to be recorded across an entire optic lobe simultaneously. This approach was optimized to image the optic lobe of *Octopus vulgaris* hatchlings, however, this method could easily be adapted to other brain regions, for example the learning and memory center (vertical lobe) (19), to other cephalopod species (squids, cuttlefish) or to other small invertebrates. This approach has also been recently adapted to another cephalopod organ, the posterior salivary gland, which is neuronally controlled to release cephalopod-specific toxins for prey capture (20).

Our approach involves dissecting *Octopus vulgaris* hatchling brains, generating acute brain slices, imaging calcium activity using a calcium-sensitive dye and an epifluorescent microscope, applying neurotransmitters exogenously using an automated perfusion system and finally analysing the dynamics of the responses using Suite2p (21) and a custom image analysis pipeline. The brains of *Octopus vulgaris* hatchlings are relatively small, approximately 540 µm anterior/posterior, 560 µm ventral/dorsal and 750 µm left/right. On the one hand, their small brains are tractable, as we can examine the whole network at high spatial resolution simultaneously. On the other hand, small brains present challenges, and this protocol describes ways to overcome them.

Before starting these experiments, we need access to *Octopus vulgaris* hatchlings. All experiments were performed on animals 1 day post hatching, however, this protocol has also been successfully applied to animals up to 1 week post hatching. We mated adult male and female wild caught *Octopus vulgaris* in the Instituto Español de Oceanografía (IEO, CSIC, Tenerife, Spain) (22). *Octopus vulgaris* females typically spawn ∼300,000 embryos over a two-week period (23). The embryos (stage III to stage XVII) were shipped to KU Leuven in Belgium within ∼ 12 hours in 50 ml falcon tubes filled with fresh oxygenated seawater at room temperature. Embryos were then incubated until hatching in a closed system in the Laboratory of Developmental Neurobiology (KU Leuven), Belgium at 18.5 °C (24). Embryos were staged daily until reaching stage XIX.1, after which they were kept in darkness for eight days. Exposure to light during this period can induce premature hatching, so the embryos were maintained in the dark until the end of the incubation period. Hatching was then triggered by exposure to light. At this stage, the yolk had been completely consumed, indicating normal development. Hatchlings displayed normal swimming behavior and were used within 24 hours.

### Institutional permissions

All procedures involving adults and hatchlings were approved by the ethical board and competent authority on animal experimentation from CSIC (permit CEIBA 1377-2023 and 1610-2024) and KU Leuven (permit P080/2021), in compliance with Directive 2010/63/EU. Readers will need to acquire permission from their respective institutions.

### Prepare Solutions

**Timing: 2 hours, to be completed at least a day before the experiment**.

1. Prepare octomedia Refer to Materials and Equipment for recipe.
2. Prepare highK^+^ octomedia Refer to Materials and Equipment for recipe.
3. Prepare acetylcholine stock solution
  a. Acetylcholine hydrochloride (molecular weight: 181.66 g/mol) is stored at 4°C.
  b. First make a 1M stock solution by adding 2 ml milliQ water and 363.32 mg acetylcholine.
  c. Mix briefly with a vortex.
  d. Store at -20°C as aliquots of 100 µl.
4. Prepare CAL-520 stock solution
  a. CAL-520 (molecular weight: 1102.95 g/mol) was purchased as 50 µg aliquots and stored at - 20°C.

**Note:** Dry aliquots of CAL-520 have a longer storage life so only do the following steps when you plan to do experiments within the next month.

b. Add 9 µl DMSO to a 50 µg aliquot to generate a 5 mM stock solution.

**CRITICAL:** Protect CAL-520 from light by covering the tube in tinfoil.

**CRITICAL:** Place aliquot on a vortex for 30 minutes to ensure the solution has fully dissolved.

c. Store CAL-520 5 mM stock solution in the freezer for up to 1 month.

**CRITICAL:** Avoid freeze thaw cycles of 5 mM solution, but we observed refreezing once or twice did not dramatically reduce the fluorescent signal.

**Optional:** Pluronic F-127 can be added to improve the solubility of CAL-520, but we obtained good signal without it.

### Key resources table

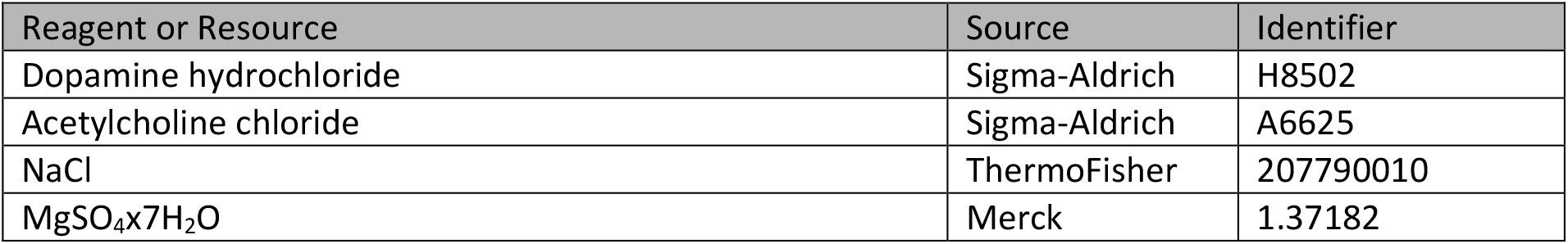
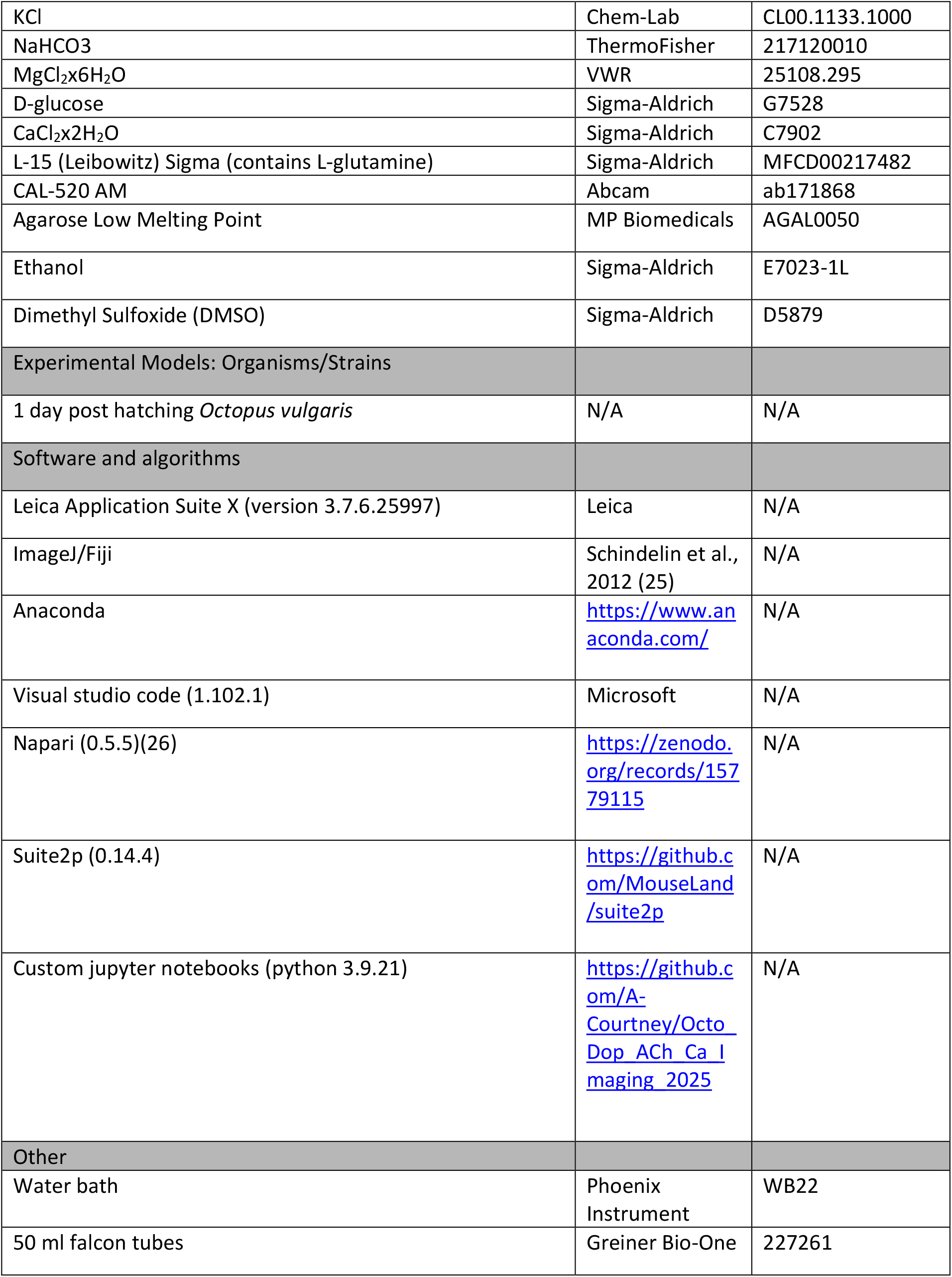
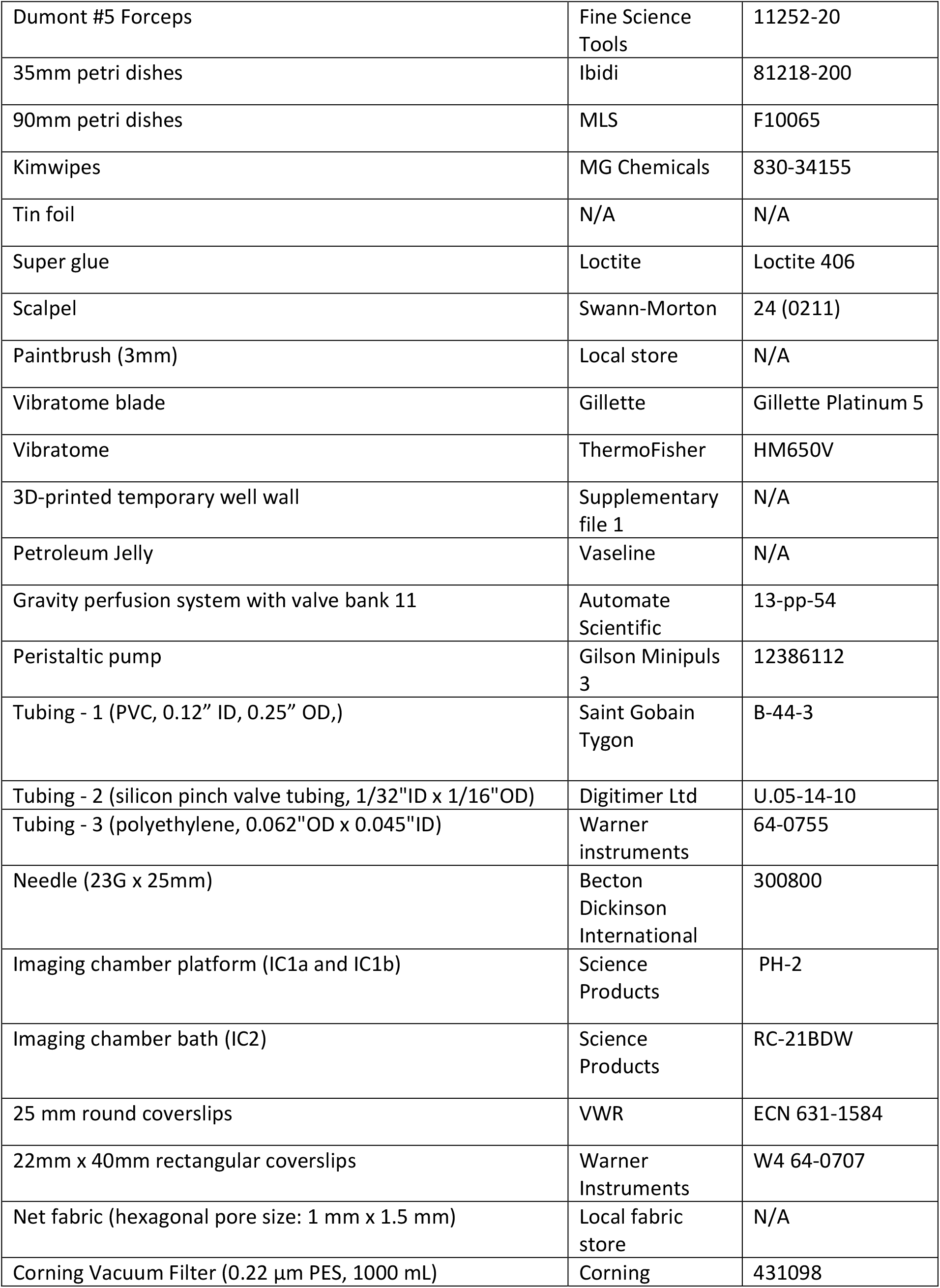
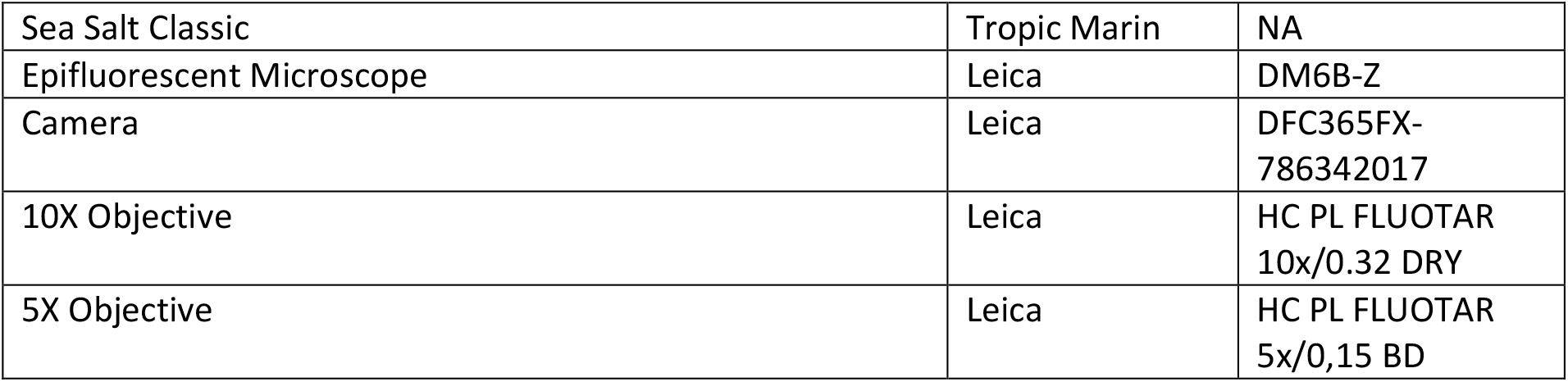

## Materials and equipment setup

1. Assemble the perfusion system as seen in Figure 3A, and per the supplier manual.
2. Generate automated protocols for the perfusion system, as per the supplier manual. **Note:** Our protocol involves 90s octomedia (Channel 1, 0-90s), 45s ligand (Channel 2 or Channel 3, 90-135s), 135s octomedia (Channel 1, 135-270s) and 90s highK^+^ octomedia (Channel 8, 270-360), 6 minutes total.
3. Generate 3D printed temporary well. **Note**: The stl file is available as supplemental file 1. We used a Creality Ender 3 Pro 3D printer (nozzle size: 0.4mm) and GEETECH 1.75mm polylactic acid (PLA) filament in ‘Silk Copper’. However, equivalent 3D printers/filament will also work.

**4. Octomedia**

- Add 500 ml L15 to 1 L beaker
- Stir continuously with a magnetic stirrer to ensure everything has dissolved
- Add the following:

| Reagent |  | Amount to add (g) | In mM | Final concentration including L15 composition (mM) |
| --- | --- | --- | --- | --- |
| NaCl | Sodium chloride | 6.25 | 214.04 | 351.97 |
| MgSO<sub>4</sub>·7H<sub>2</sub>O | Magnesium sulfate heptahydrate | 3.225 | 26.17 | 26.984 |
| KCl | Potassium chloride | 0.172 | 4.61 | 9.94 |
| NaHCO<sub>3</sub> | Sodium bicarbonate | 0.096 | 2.29 | 2.29 |
| MgCl<sub>2</sub>·6H<sub>2</sub>O | Magnesium chloride hexahydrate | 2.85 | 28.04 | 29.026 |
| D-glucose |  | 3.43 | 38.08 | 38.08 |
| CaCl<sub>2</sub>·2H<sub>2</sub>O | Calcium chloride dihydrate | 0.744 | 10.12 | 11.38 |
| C<sub>3</sub>H<sub>3</sub>NaO<sub>3</sub> | Sodium Pyruvate | 0 | 0 | 5 |
| KH<sub>2</sub>PO<sub>4</sub> | Potassium phosphate monobasic | 0 | 0 | 0.44 |
| Na<sub>2</sub>HPO<sub>4</sub> | Sodium phosphate dibasic | 0 | 0 | 1.34 |

- Adapt pH to 7.6-7.7 with KOH
- Filter with vacuum pump and Corning system

[Store at 4°C for up to 1-2 weeks]

**5. HighK^+^ octomedia**

- Add 500 ml L15 to 1L beaker
- Stir continuously with a magnetic stirrer to ensure everything has dissolved
- Add the following:

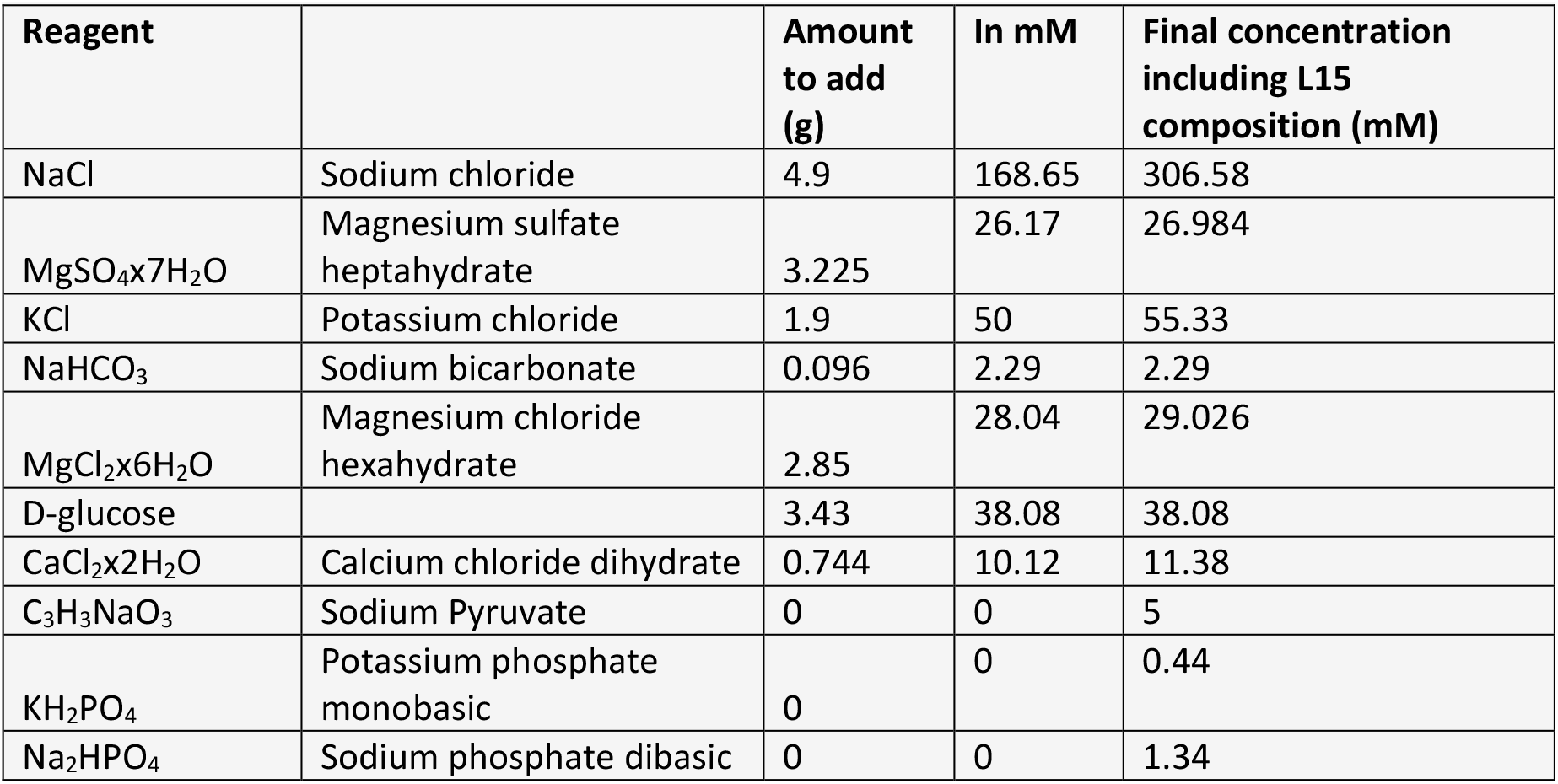

- Adapt pH to 7.6-7.7 with KOH
- Filter with vacuum pump and Corning system

[Store at 4°C for up to 1-2 weeks]

**6. Filtered Artificial Seawater (FASW)**

- Add 1L milliQ water to 35g sea salt
- Stir continuously with a magnetic stirrer to ensure everything has dissolved
- Filter with vacuum pump and Corning system

[Store at room temperature for up to 3 months]

**7. 2% ethanol in Filtered Artificial Seawater**

| Reagent | Amount to add (ml) |
| --- | --- |
| Filtered artificial seawater | 49 |
| Ethanol | 1 |
| **Total** | **50** |

[Store at room temperature for up to 1 week]

**8. Leica settings**

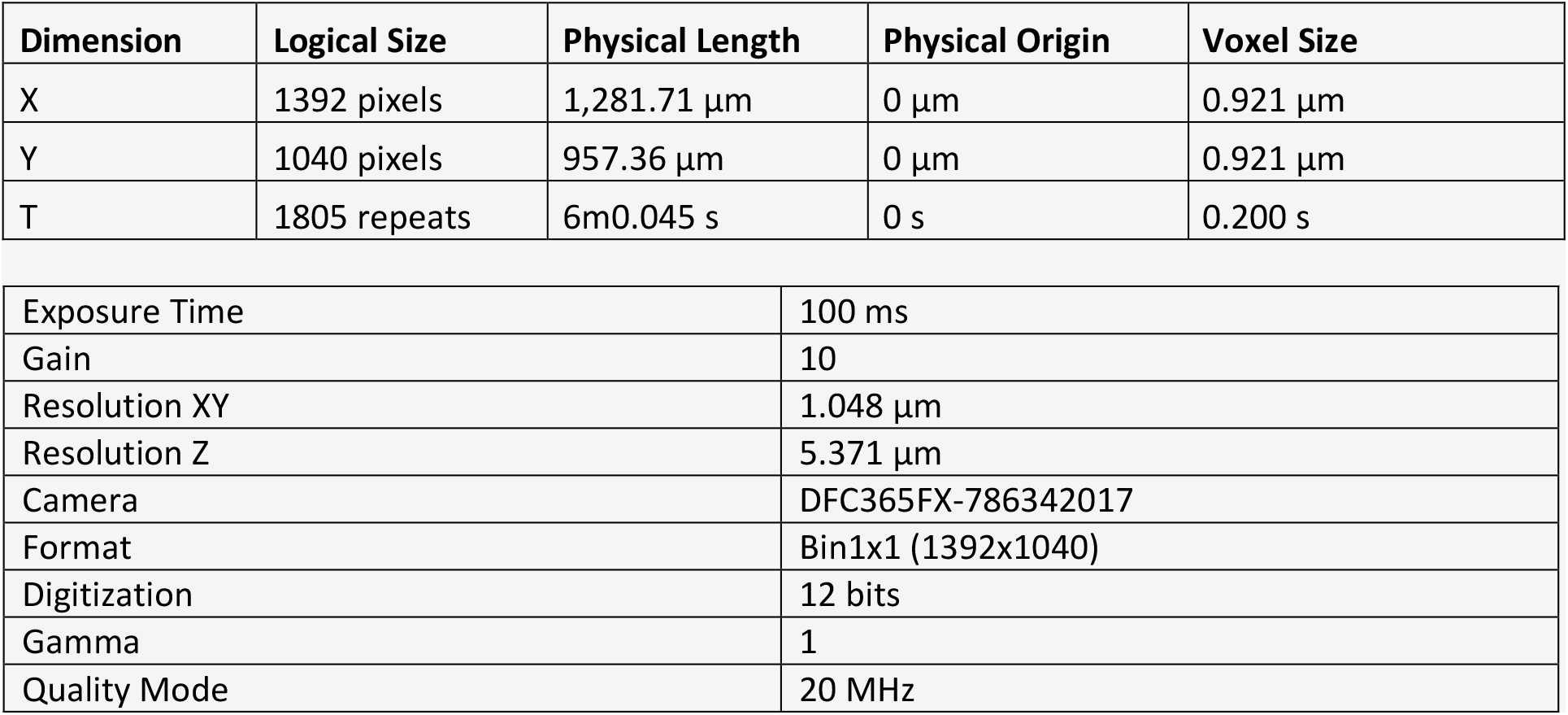

| Dimension | Logical Size | Physical Length | Physical Origin | Voxel Size |
| --- | --- | --- | --- | --- |
| X | 1392 pixels | 1,281.71 $\mu\text{m}$ | 0 $\mu\text{m}$ | 0.921 $\mu\text{m}$ |
| Y | 1040 pixels | 957.36 $\mu\text{m}$ | 0 $\mu\text{m}$ | 0.921 $\mu\text{m}$ |
| T | 1805 repeats | 6m0.045 s | 0 s | 0.200 s |

|  |  |
| --- | --- |
| Exposure Time | 100 ms |
| Gain | 10 |
| Resolution XY | 1.048 $\mu\text{m}$ |
| Resolution Z | 5.371 $\mu\text{m}$ |
| Camera | DFC365FX-786342017 |
| Format | Bin1x1 (1392x1040) |
| Digitization | 12 bits |
| Gamma | 1 |
| Quality Mode | 20 MHz |

## Step-by-step method details

**Note:** This protocol is separated into three major steps; step one is the brain slice preparation, step two is the calcium imaging, and step three is the image analysis. Step one and two need to be performed on the same day and step 3 can be performed any time after that. Performing step one and two on a single day takes ∼ 14 hours for 8 brains. We therefore suggest that these experiments are done in tandem with two experimenters. The first experimenter does the ‘morning shift’ and prepares the brain slices, while the second experimenter does the ‘afternoon/evening shift’ and collects the calcium imaging data.

### Prepare 3% agarose in octomedia

**Timing: 10 minutes**

1. Fill water bath with water and heat up to 42 °C.
2. Get two 50 ml falcon tubes, fill one with 50 ml octomedia.
3. Weigh out 1.75 g agarose and mix with octomedia back and forth between the two falcon tubes.

**CRITICAL:** Ensure that you use an agarose with a low melting temperature (26 °C) and a high strength (500 g/cm^2^), as other types of agarose we tested did not work.

4. Put the falcon tubes in an empty 200 ml beaker with the lids off and put in the microwave for 20 seconds, then mix back and forth, then another 20 seconds, and mix back and forth, until the agarose has dissolved but before the solution boils.

**CRITICAL:** Do not let the solutions boil. You want to make sure the agarose has fully dissolved in the octomedia, but you also don’t want to generate too many bubbles.

5. Have equal amounts of solution in both falcons, place the lids on the falcons and place them in the water bath lying down.

**CRITICAL:** If there are too many bubbles in the agarose the brain will be more likely to pop out of the agarose during vibratome sectioning. The longer the agarose is left in the water bath the less bubbles you’ll have, so do this step as early as possible.

**Note:** Any agarose you don’t use can be stored in the fridge and re-microwaved the next day.

### Brain dissection

**Timing: 1-2 hours for 8 brains, this may take longer for inexperienced users**.

6. Gather tools for dissection:
  a. 2 sharpened forceps for dissecting
  b. 1 old/blunt forceps for adding holes to the foil
  c. Glass pipette with opening larger than a hatchlings for transferring hatchlings/brains
  d. Foil
  e. Kimwipes
  f. 35 mm petri dishes
  g. 90 mm petri dishes
  h. 200 ml octomedia chilled on ice
7. Cut eight 1 cm x 1 cm foil squares and add holes using the blunt forceps.
Lay out 35 mm petri dishes (which will be filled with agarose later), label each as 1 to 8 (number of brains you intend to dissect).
8. Put 2% ethanol/ASW solution in 90 mm petri dish (anesthesia bath).

**Note:** EtOH is well established as an effective anesthesia for cephalopods (27).

10. Transfer one hatchling to anesthesia bath for at least 5 minutes using the glass pipette.

**CRITICAL:** You need to make sure they are fully anesthetized before the dissection (Figure 1A-C). Live animals will swim around the petri dish and have an orange/yellow colour to their skin. Anesthetized animals will be fully immobilized, and their skin will have become almost completely transparent.

**Figure 1.**
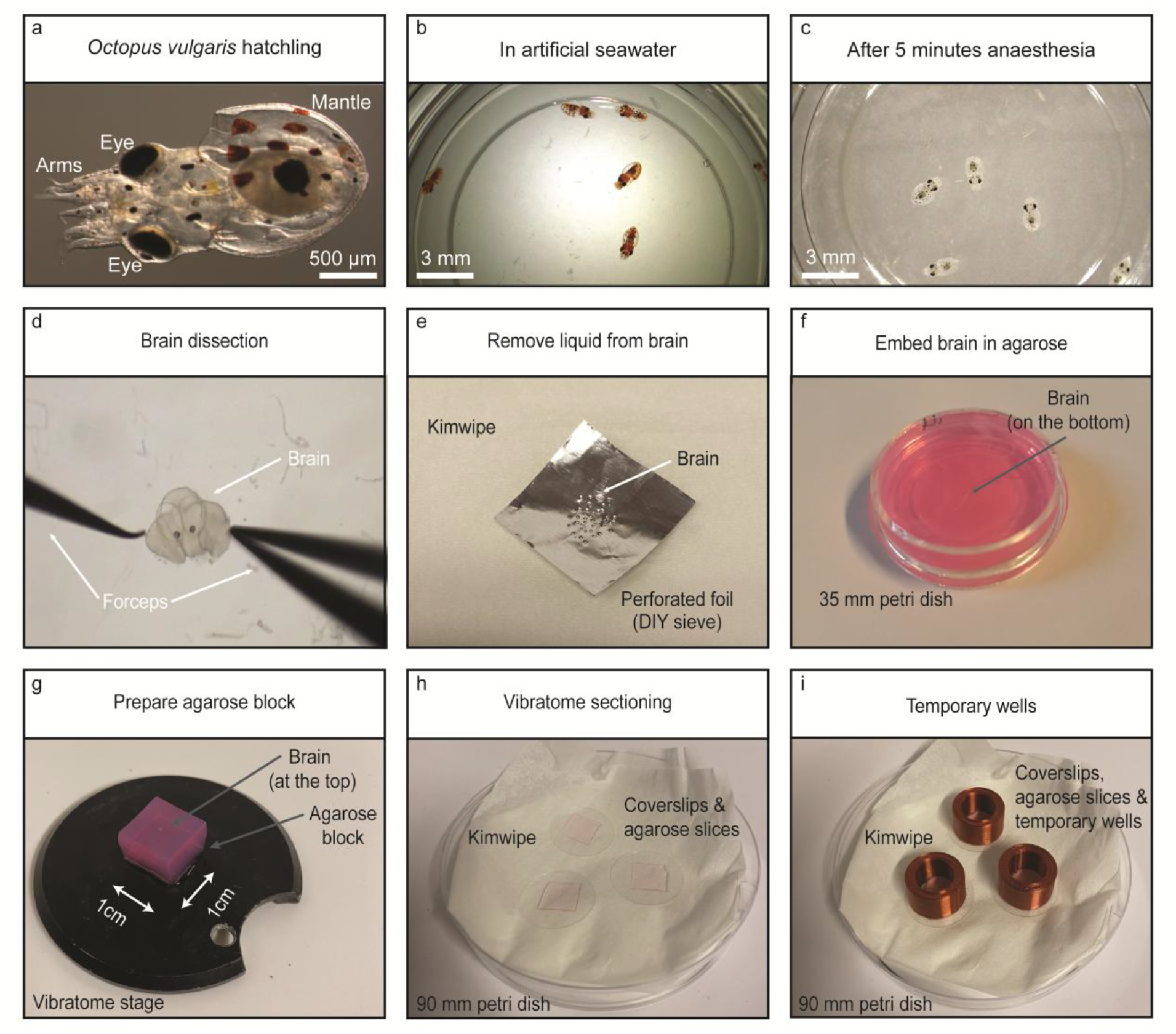
Experimental workflow for preparing acute brain slices from *Octopus vulgaris* hatchlings. (**A**) *Octopus vulgaris* 1 day post hatching (“hatchling”). (**B**) *Octopus vulgaris* hatchlings in artificial seawater. (**C**) *Octopus vulgaris* hatchlings after 5 minutes in anesthesia solution (2 % EtOH in FASW).(**D**) Dissected *Octopus vulgaris* hatchling brain. (**E**) *Octopus vulgaris* hatchling brain on perforated foil on kimwipes to remove liquid. (**F**) *Octopus vulgaris* hatchling brain embedded in 3 % agarose/octomedia in 35mm petri dish. (**G**) *Octopus vulgaris* hatchling brain embedded in 3 % agarose/octomedia as a 1 cm x 1 cm block on the vibratome stage. (**H**) 200 µm vibratome sections (*Octopus vulgaris* hatchling brain embedded in 3% agarose/octomedia) on circular coverslips. (**I**) 3D printed temporary wells placed around the vibratome sections.

11. Prepare 90 mm petri dish filled with chilled octomedia, on ice, under the dissecting microscope.
12. Perform the brain dissection:

**Note:** A video of an example dissection can be found in supplementary file 2.

a. Hold the arms and peel off the top layer of skin and mantle.
b. Then remove all the other arms and mouth.
c. Carefully remove the eyes by pinching the optic nerves and pulling the eye away gently.

**Note:** You can leave the eyes on, but we observed contractions of the eye muscles during calcium imaging (when we applied the highK^+^ solution), and thus we removed them for all our experiments.

**Note**: The two optic lobes, central brain (suboesophageal/supraoesophageal mass) and statocyst will be clearly visible (Figure 1D).

**CRITICAL:** Ensure as much skin as possible has been removed. Any remaining skin or musculature surrounding the brain may contract during calcium imaging.

### Agarose embedding

**Timing: 1 hour**

**CRITICAL:** brains should be as dry as possible before embedding them in agarose. If there is too much water around the brain when it goes into the agarose it will pop out during vibratome sectioning.

13. Place foil square with holes on kimwipe.
14. Transfer the brain from the dissection dish to the foil square using a glass pipette. This essentially creates a small DIY sieve and allows you to remove some of the water surrounding the brain.

**CRITICAL:** The brains stick to plastic pipettes so when you’re moving the brains make sure you use glass, or alternatively, coat the plastic pipettes with 0.2% tween to prevent the brains from sticking.

15. Move the foil square to a dry part of the tissue and continue to add more holes close to the brain to remove as much water as possible (Figure 1E).
16. Take one of the falcon tubes containing melted agarose from the water bath.

**Note:** The temperature of the agarose will start to drop as soon as you remove it from the water bath, so you only have a few minutes until the agarose will harden.

17. Fill a 35 mm petri dish with agarose to the top.
18. Place the falcon tube back in the water bath.
19. Transfer brain from foil to agarose by gently pushing the brain off the foil using a forceps, or dipping the foil into the agarose.
20. Under the dissection microscope swirl the brain ∼10 times in the agarose to remove additional water, you do this by moving your forceps through the agarose around the brain, you don’t have to directly touch the brain to move it around.
21. Under the dissection microscope position the brain on the bottom of the petri dish in the center (Figure 1F).

**Note:** To achieve either coronal or transversal brain sections position the brain as shown in Figure 2 A-E. The optic lobe sections look the same in either coronal or transversal sections, but the central brain sections look very different. Choosing coronal or transversal sections is more relevant for cases where you want to record from central brain cells (Figure 2F/G).

**Figure 2.**
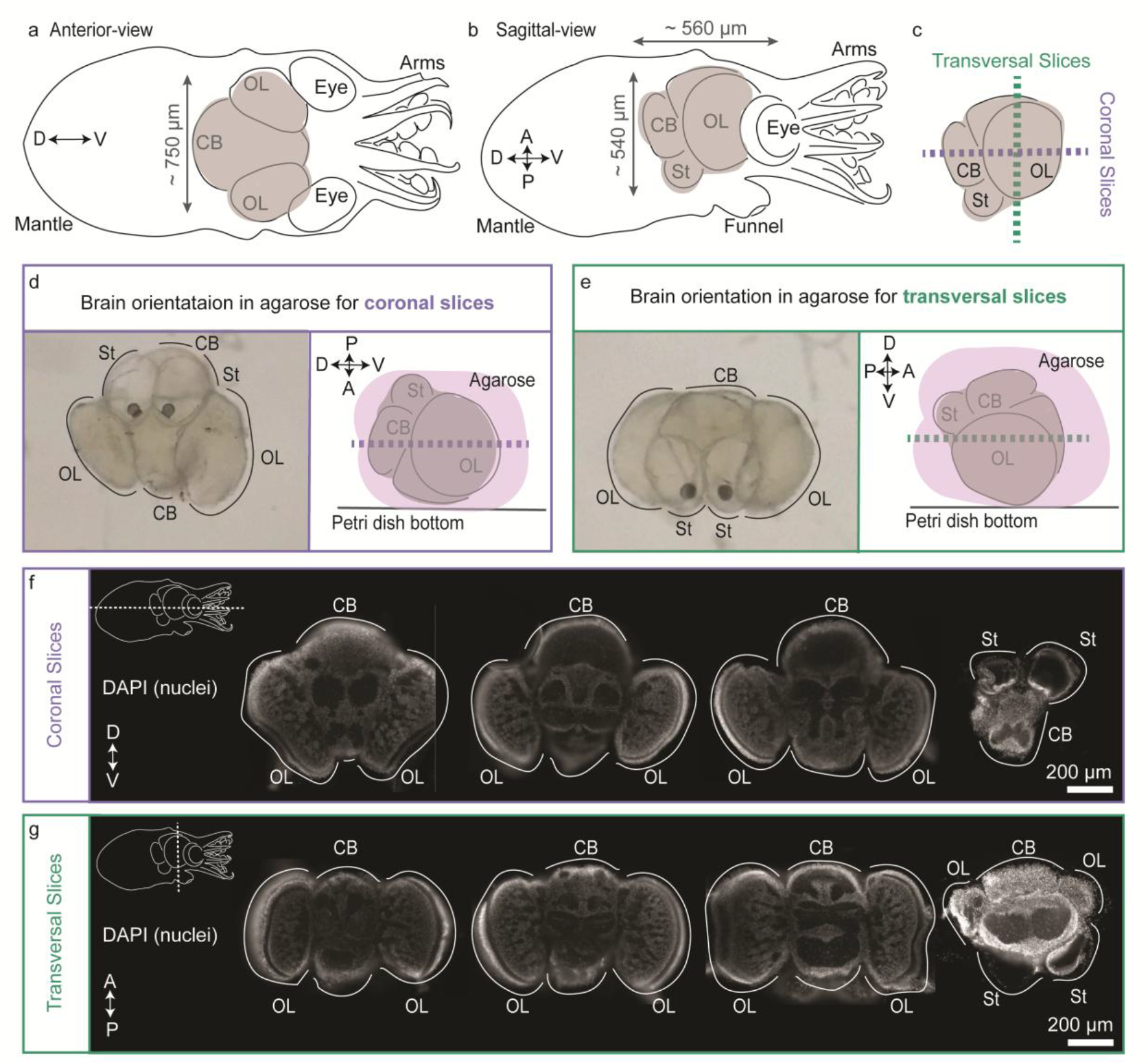
Preparing coronal and transversal brain slices from *Octopus vulgaris* hatchlings. (**A**) Schematic of *Octopus vulgaris* 1 day post hatching (‘hatchling’), viewed anteriorly with major brain regions highlighted. (**B**) Schematic of the *Octopus vulgaris* hatchling, viewed sagittally. (**C**) Schematic of the *Octopus vulgaris* hatchling brain, viewed sagittal, with coronal and transversal cutting planes highlighted. (**D**) Left: dissected *Octopus vulgaris* hatchling brain in the orientation required to achieve coronal slices in agarose, when viewing from above. Right: schematic of the same brain in this orientation when viewed sagittally. (**E**) Left: dissected *Octopus vulgaris* hatchling brain in the orientation required to achieve transversal slices in agarose, when viewing from above. Right: schematic of the same brain in this orientation when viewed sagittally. (**F**) Representative 200 µm coronal brain slices taken from different *Octopus vulgaris* hatchlings. The brains were sliced live using a vibratome (as described in this protocol) but were then fixed in 4% paraformaldehyde and the nuclei were stained with DAPI, and imaged on the same epifluorescent microscope used for calcium imaging (described here). This allowed visualization of morphology more clearly. (**G**) Representative 200 µm transversal brain slices taken from different *Octopus vulgaris* hatchlings. The brains were prepared as described in F. D: dorsal, V: ventral, P: posterior, A: anterior, CB: central brain, OL: optic lobe, St: statocyst.

**Figure 3.**
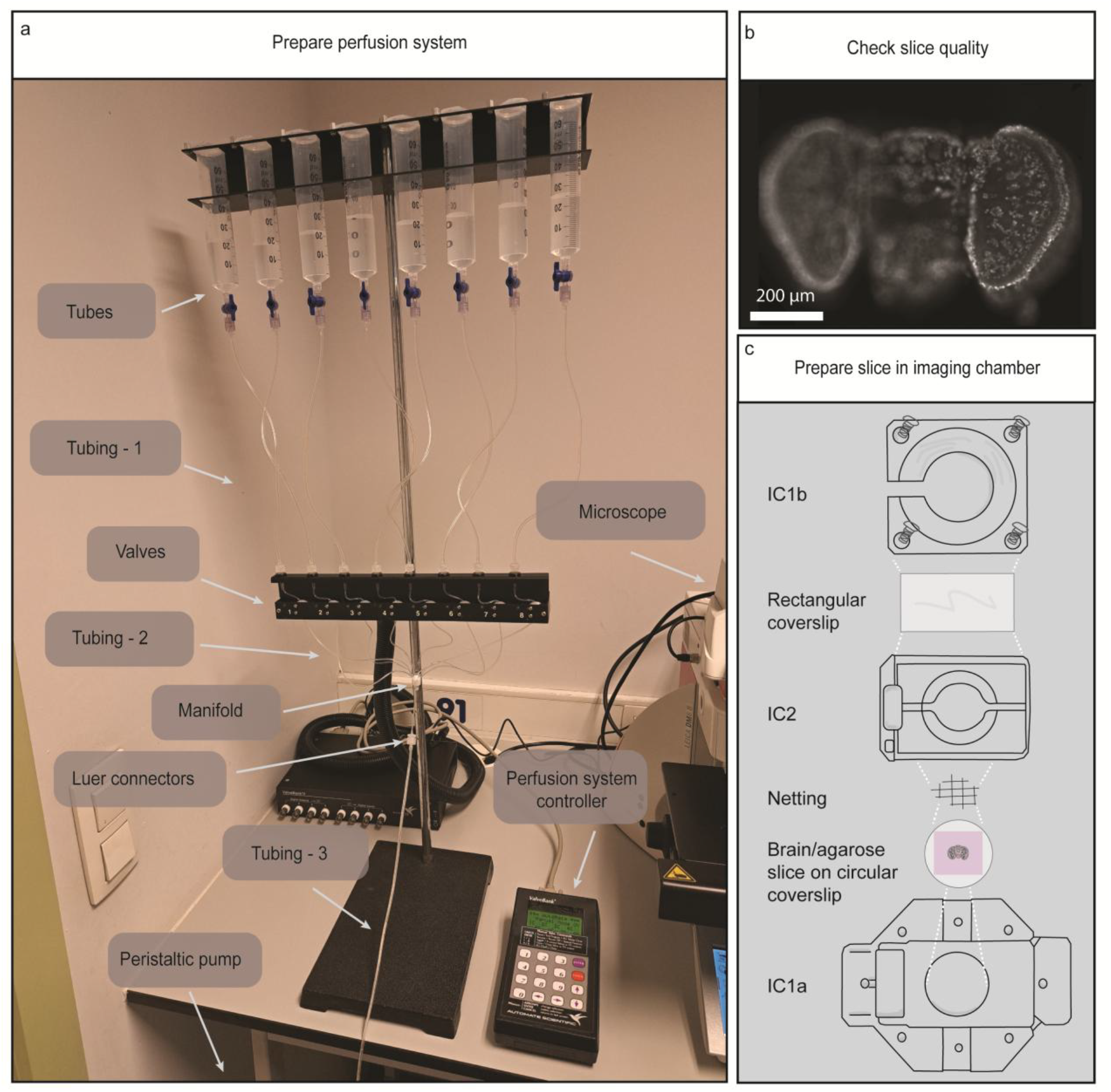
Setting up the perfusion system and preparing slices for imaging. (**A**) How to assemble the perfusion system with all key components labelled. (**B**) Live *Octopus vulgaris* hatchling brain slice stained with CAL-520 calcium indicator. (**C**) How to prepare the *Octopus vulgaris* hatchling brain/agarose slices in the imaging chamber.

22. Gently push the agarose dish to the side and let it solidify for 5 minutes at room temperature.
23. Then place the dish on ice until you’re ready to section it on the vibratome. Cover the ice box with foil to protect from light.

**Note:** We haven’t been strict with how long the brains are on ice before sectioning. It is usually 10-30 minutes.

24. Repeat the brain dissection and agarose embedding steps until you have enough brains (aim for 8 per day).

### Vibratome sectioning

**Timing: 2 hours**

25. Prepare the vibratome:
  a. Add the blade.
  b. Add chilled octomedia to the bath.
  c. Apply the following settings: speed: 5, amplitude 0.7, slice thickness: 200 µm.
26. Make 5 µM CAL-520 working solution:
  a. Add 9 µl 5mM CAL-520 (from the -20°C) to 9 ml octomedia in a 15 ml falcon tube.
  b. Protect from light with foil.
  c. Keep at room temperature.
27. Prepare 90 mm petri dishes with a kimwipe and three circular coverslips, label each petri dish 1 to 8 (for each brain).
28. Prepare 1 cm x 1 cm agarose block:
  a. Take the 35mm petri dish containing dissected brain in agarose.
  b. Use a scalpel to remove a chunk from the side of the agarose in the petri dish and then gently remove the rest of the agarose block from the petri dish using the opening you just created.
  c. Remove the agarose surrounding the brain using a scalpel so that the agarose measures a 1cm x 1cm square (Figure 1G).
  d. The brain was placed at the bottom of the petri dish and then solidified in the agarose (step 21 above). The agarose block should now be flipped, so that the brain is now at the top. The height of the agarose block should be the height of the 35mm petri dish (if filled to the top) and will therefore be consistent across agarose blocks.
29. Glue the agarose block to stage, wait at least one minute for the glue to dry.
30. Run vibratome protocol:
  a. Define the window containing the agarose block.
  b. Press run, use high speed before the blade reaches the brain, use the lowest speed (1-5), as the blade reaches the brain.

**Note:** Each 1 dph brain usually yields three slices.

31. Place the brain/agarose slice on a circular coverslip using a paintbrush (Figure 1H)

**Note:** in earlier versions of this protocol we left the brain/agarose slices free floating in 12 well plates. However, they were difficult to manipulate and as the brains are so small, we often lost them. The quality of the slices was less reliable too. To overcome this see we decided to place the slices directly on circular coverslips (which are compatible with the imaging chamber used later in the protocol), and creating a temporary well around them to enable incubation in CAL-520 (step 33 below).

32. Place the slice on the coverslip so the surface of the section that was cut is now facing up.

**Note:** This is important for imaging later as we will use an upright objective and we want the cut surface to be facing the objective. The first slice will be cut only on its under surface, the middle slice will have been cut on both surfaces, the last slice will have been cut on its top-most surface only.

33. Create temporary well around each slice:
  a. Remove excess liquid from each of the three slices using a P200 pipette.
  b. Coat the bottom of the 3D-printed well wall with Vaseline.
  c. Place the 3D printed well wall around the slice on the coverslip and push down to create a seal (Figure 1I).

### Cal-520 incubation

**Timing: 1 hour**

34. Add 150 µl of CAL-520 to each temporary well.
35. Incubate for at least an hour in the dark.
36. After 1 hour add another 150 µl octomedia to provide more nutrients for the slice.

**Note:** Leave the slice in this solution until imaging. Longer periods in CAL-520 did not seem to negatively affect the slice.

**Note:** From this point forward the slice and all solutions are kept at room temperature and in the dark.

**Pause point:** You can take a break for 30 minutes-1 hour here while the slices are incubating in CAL-520. If you are performing this protocol with two researchers, the second researcher can now take over.

### Preparing the perfusion system

**Timing: 30 minutes**

37. Prepare 100 µM dopamine:
  a. Dopamine (molecular weight: 189.64 g/mol) is stored at 4°C.
  b. Add 1.8 mg dopamine and to 100 ml octomedia.

**CRITICAL:** You need to prepare dopamine solutions on the day of the experiment as you cannot freeze stock solutions. This does not apply to acetylcholine.

38. Prepare 100 µM acetylcholine by combining 10 µl 1 M ACh and 100 ml octomedia.
39. Prepare the perfusion system with all solutions as seen in Figure 3A:

**Note:** all solutions are maintained at room temperature.

a. Add 50 ml octomedia to tube 1.
b. Add 10 ml octomedia to tube 4, 5, 6, and 7.
c. Add 30 ml 100 µM dopamine in octomedia to tube 2.
d. Add 30 ml 100 µM acetylcholine in octomedia to tube 3.
e. Add 50 ml highK^+^ octomedia to tube 8.
f. Wash all liquids through the system by adding a tube from the peristaltic pump to the manifold on the perfusion system.
g. Turn on the peristaltic pump to the highest speed.
h. Open all valves via the control panel.
i. Once there is liquid and no bubbles in the perfusion system tubing, close all valves via the control panel.
j. Remove the tube to the peristaltic pump from the manifold.

40. Check each valve is working correctly:
  a. Place the manifold opening in a canonical flask so you can observe the flow.
  b. Open and close each valve individually to ensure the opening/closely is working effectively and the flow is relatively consistent across valves.

**Note:** If the liquid is not flowing within one valve it’s likely due to trapped air so squeezing the tubes and opening/closing the valves will eventually get the liquid moving.

c. Then attach the tubing that will attach the imaging chamber to the manifold and open valve 1 (octomedia) until it fills with liquid to the end of the inlet tubing.

**Note:** The time it takes for liquid to travel from the manifold through this tubing is approximately the timing of the delay between valve opening and the solution reaching the imaging chamber. Keeping the inlet tubing as short as possible will reduce this delay period.

### Setting up the epifluorescent microscope

#### Alternative

for calcium imaging we decided to use an upright epifluorescent microscope with an air objective. Alternatives could also include (although we haven’t explicitly tested them), a water dipping upright objective, and inverted epifluorescent microscope, or a 2-photon microscope in either an upright or inverted configuration.

**Timing: 15 minutes**

41. Turn on the computer, microscope and light source.
42. Turn the knob on the light source to the lowest intensity value.
43. Open the Leica Las X software on the computer.
44. Apply the settings listed in table 8.
45. Keep exposure as high as possible while allowing five frames per second.

**CRITICAL**: Exposing the tissue to too much light will result in photobleaching and analysing changes in fluorescence over time due to neural activity will be challenging. We reduced photobleaching by reducing the iris size and having the light source at the lowest intensity. To determine the rate of photobleaching, maintain the slice in octomedia, record a video for 5 minutes, and plot the intensity of individual cells over time.

### Imaging

**Timing: 4-10 hours for 8 brains**

46. Place a coverslip with a brain/agarose slice and temporary well on the bottom of the imaging chamber (IC1a) and place it on the epifluorescent microscope stage.
47. Use the 5X objective and the iris fully open, observe the quality of the slice (Figure 3B).

**Note:** A ‘good quality’ slice has relatively bright fluorescence, normal brain morphology and many cells in focus in one plane. The middle slice is usually the best slice from each brain, and we usually get 1-2 ‘good quality’ slices from each brain. Assessing the ‘quality’ of the slices is somewhat subjective but it becomes obvious as the experimenter gains experience.

48. If the slice is ‘good quality’, then proceed with the following steps, if the slice is ‘poor quality’ then check another slice.
49. Remove the imaging chamber from the microscope stage, and remove the temporary well from the coverslip.
50. Prepare the slice in the imaging chamber (Figure 3C):
  a. Use a dissecting microscope (with as little light as possible) to place the netting on top of the slice.

**Note:** The netting should have a pore size large enough to create a window around your brain tissue, while the fibers sit on the agarose. We used a netting with a hexagonal pore size of 1 mm x 1.5 mm.

b. Add the IC2 bath chamber on top.

**Note:** The overall size of the netting should be slightly larger than the opening in IC2, so when you place IC2 on top it clamps the brain/agarose slice in place.

c. Add the rectangular coverslip on top of IC2.
d. Add IC1b on top and screw down the four screws.

**Note:** The rectangular coverslip acts as a transparent lid and makes this a closed imaging chamber. As we are imaging with a dry upright objective, we found if we don’t add this lid then changes in the water level during perfusion created a variable lens effect and we could not image the slices clearly.

e. Fill the imaging chamber with octomedia by putting the inlet tubing into the imaging chamber, holding the imaging chamber in your hand at an angle, opening valve 1 via the control panel and waiting for the chamber to fill with octomedia. Then close valve 1 again.

**Note**: Make sure to do this step at an angle otherwise you will get bubbles in the chamber.

51. Place the imaging chamber on the epifluorescent microscope stage with the inlet tubing on the right.
52. The outlet tubing is attached to the peristaltic pump and contains a 23G x 25mm needle. Place the needle into the reservoir opening in IC2 bath chamber on the left.

**Note:** the imaging chamber and inlet/outlet tubings had to be placed on the stage in this orientation to not interfere with the position of the other objectives, and thus this could be orientated differently on your system.

53. Turn on the peristaltic pump.
54. Open valve 1 (octomedia) via the control panel and make sure the solutions are flowing without leakage.

**Note:** The ideal speed of the peristaltic pump will depend on the rate at which the solutions are flowing into the chamber. Ideally it will come in and be removed at a constant rate. Start at the highest speed setting and work down to the slowest speed that prevents overflow. The first time you do this, test it on the bench without any brain/agarose slice.

**CRITICAL:** do not let your imaging chamber overflow as solution leakage could damage your microscope.

55. With the 5X objective still in place, center the brain slice in the field of view. Then change to the 10X objective and recenter the slice. Adjust the focus to get as many cells as possible in the optic lobes in focus. Reduce the iris size as much as possible.
56. Leave the octomedia running for at least 10 minutes before recording to make sure the slice is acclimatized to the imaging chamber.

**Note:** Ensure the fluorescence is off to avoid photobleaching when not imaging.

57. Run the appropriate pre-programmed perfusion protocol by pressing start on the control panel and starting the video in the Las X software at the same time.
58. Once the imaging has finished, open valve 1 and run octomedia for 1 minute to ensure the inlet tubing is filled with octomedia again.
59. Then close all valves.
60. Remove the imaging chamber from the stage, dispose of the slice, rinse the chamber with water.
61. Repeat steps 46-60 with another slice until all slices have been checked and/or imaged.

### Clean-up

**Timing: 10 minutes**

62. Remove the inlet tubing from the manifold on the perfusion system.
63. Add tubing from the peristaltic pump to the manifold (as seen in Figure 3A).
64. Turn on the peristaltic pump to the highest speed.
65. Open all valves via the control panel.
66. Once all the solutions have been removed, close all valves via the control panel.
67. Add 30 ml water to all tubes.
68. Open all valves via the control panel and wait for all water to wash through.
69. Remove the peristaltic pump tubing from the manifold on the perfusion system.
70. Rinse imaging chamber and inlet tubing with water and dry with tissue.
71. Dispose of all solutions.
72. Wash all glassware.
73. Save your data and turn everything off.

**Pause point:** The following data analysis steps can be performed later.

### Image analysis

**Note:** The steps of our image analysis pipeline are outlined in Figure 4. Registration, region of interest (ROI) detection, and extracting mean intensity over time was performed in Suite2p. While custom jupyter notebooks were then used to manually define anatomical regions, filter ROIs, categorize ligand responses, define their dynamics and finally visualization. We decided to normalize the response from each ROI using min/max values as we wanted the highK^+^ response to act as the ‘maximum’ value for each cell. We could then examine the ligand response relative to that. However, calcium imaging data from epifluorescent microscopes have blurry background contamination from neuropil and out of focus cells, and as our cell segmentation was performed using an anatomical approach, we wanted to filter out any segmented cells that predominantly showed this contamination and thus were therefore not a ‘real’ cell. We manually defined ‘Neuron’ and ‘Neuropil’ ROIs and calculated the max slopes and found that ‘Neuropil’ ROIs have a lower max slope compared to ‘Neuron’ ROIs. We then defined a ‘slope threshold’ to filter out these cells. More details can be found in Courtney et al., 2025 (13). This slope threshold needs to be recalculated for each dataset (step 77/78). The following steps outline how to install and run this pipeline on our sample dataset (10.5281/zenodo.18981973) and how to adapt it for the readers dataset.

**Figure 4.**
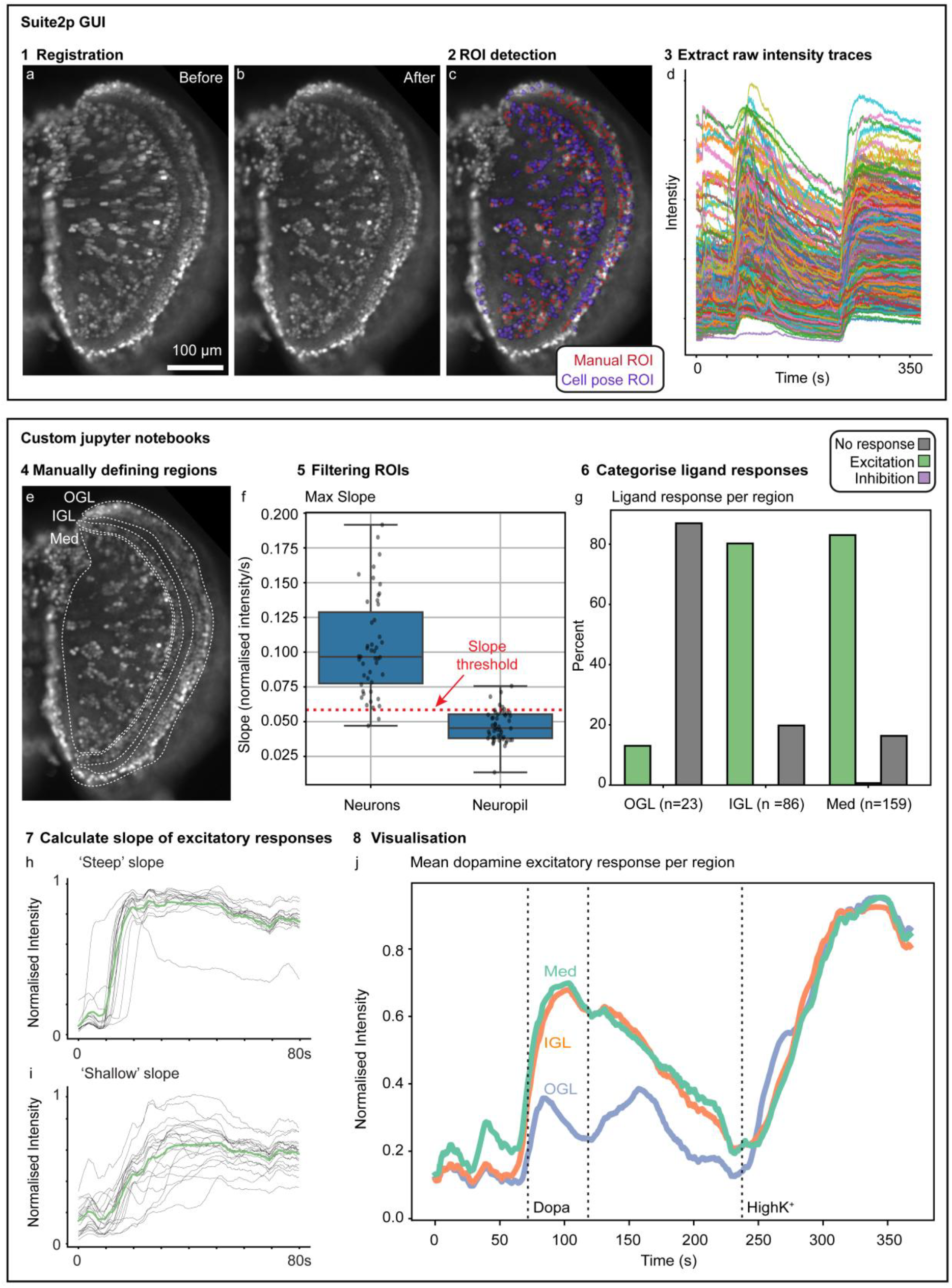
Key steps in the calcium imaging analysis pipeline using Suite2p and custom Jupyter notebooks. (**A**) Maximum intensity projection of *Octopus vulgaris* hatchling brain slice (optic lobe) stained with CAL-520 (calcium indicator) and imaged over the course of 6 minutes during application of 100 µM dopamine and highK^+^ solution, before suite2p registration was applied. (**B**) same as A, with suite2p registration applied. (**C**) Same as B, with Cellpose and manual ROIs highlighted. (**D**) Raw intensity traces for all ROIs extracted using Suite2p. (**E**) Same as B, with anatomical regions highlighted that were manually defined using Napari in custom Jupyter notebooks. (**F**) Max slope of the normalized intensity from manually defined neuron and neuropil ROIs, to determine the slope threshold (neuropil mean + standard deviation), calculated using custom Jupyter notebooks. (**G**) Dopamine response categories by region for a representative slice, calculated using custom Jupyter notebooks. (**H**) subset of dopaminergic responses with ‘steep’ slope (> 0.075 normalized intensity/s). (**I**) A subset of dopaminergic excitatory responses with “shallow” slopes (between 0.0125 and 0.0375 normalized intensity/s). For H and I, the mean trace is shown in green and they were calculated using custom Jupyter notebooks. (**J**) Mean traces from a representative slice per optic lobe layer, visualized using custom Jupyter notebooks. ROI: region of interest, OGL: outer granular layer (of the optic lobe), IGL: inner granular layer (of the optic lobe), Med: medulla (of the optic lobe), Dopa: dopamine.

**Timing: hours to days**

74. Preparing the data

**Note:** if you are using our sample dataset, you can skip this step.

a. Leica saves the videos as a .lif project, convert each video to .tif file using Leica LAS X software.
b. Prepare a main working directory.
c. Save each .tif file in a separate folder within the main working directory. For example, ‘12Dec25_Slice6_Dop’ or ‘12Dec25_Slice7_ACh’, as seen in our sample dataset.

75. Software installation
  a. Install anaconda as per instructions on their website: https://www.anaconda.com/download
  b. Install visual studio code (VSC) as per the instructions on their website: https://code.visualstudio.com/download
  c. Download custom jupyter notebooks and environment file from github (
https://github.com/A-Courtney/Octo_Dop_ACh_Ca_Imaging_2025) or from supplemental file 3.
  d. Open Anaconda Prompt and install mamba:

`>conda install -n base -c conda-forge mamba`
  e. Navigate to the folder containing the custom jupyter notebooks and the environment file.
  f. Create environment using the environment.yml file:

`>mamba env create -f environment.yml --channel-priority flexible`
  g. Activate the environment:

`>conda activate Octo-Ca-Im`
  h. Open VSC and install the python and the jupyter extensions.
  i. Open the folder containing the custom jupyter notebooks in VSC.
  j. Open the first notebook, select kernel and choose the ‘Octo-Ca-Im’ environment.

**Note:** the custom jupyter notebooks are now ready to be used in steps 78-81 below.

k) Install suite2p as per the instructions on their GitHub page: https://github.com/MouseLand/suite2p

**Note:** Suite2p is now ready to be used in step 76 below.

l) Install Fiji as per the instructions on their website: https://imagej.net/software/fiji/downloads

**Note:** Fiji is now ready to be used in step 77 below.

76. Registration and semi-automated ROI detection using Suite2p

**Note:** If you are working with our sample dataset you can skip this step.

a. Open anaconda prompt and type the following:

`>conda activate suite2p`

`>suite2p`
b. Within the GUI click ‘File>Run suite2p’ and another window will open.
c. Click ‘Load ops file’ and navigate to our sample dataset. Within ‘12Dec25_Slice6_Dop/suite2p/plane0’ you will find ‘ops.npy’. Choose this file to apply the settings that were optimized for our sample dataset.

**Note:** when running your own data, start with the default settings and then adapt for your videos. Also refer to suite2p documentation for more details.

d) Click ‘Add directory to data_path’ and choose the folder containing your videos in .tif format.
e) Click ‘RUN SUITE2P’.
f) Once the program has finished running, you will see the result in the main Suite2p GUI.

**Note:** the tif has been registered and is saved in ‘12Dec25_Slice6_Dop/suite2p/plane0/reg_tif’.

g) In the main Suite2p GUI click ‘T:max projection’.
h) Look at the ‘cells’ view of your slice, right click on any ROIs that are not a cell or are defining two cells, they will no longer be visible and no longer included in any downstream analysis.
i) Click File>Manual labelling.
j) Change the ROI size to 6.

**Note:** This was the ideal size for the resolution of our images/the size of the cells. This may be different for your data.

k) Click alt and right click to add ROIs on cells.
l) When you’re finished click ‘extract ROIs’ and wait for the traces to be plotted
m) The click ‘save and close’ and then you can close this window
n) Now all the ROIs will be visible in the ‘cells’ view

**Note:** this is automatically saved by Suite2p

77. Generate ground truth ‘Neuron’ and ‘Neuropil’ data using Fiji

**Note:** If you are working with our sample dataset you can skip steps 77/78.

a. Go to this folder ‘suite2p\plane0\reg_tif’
b. Load all registered tifs into Fiji
c. ‘Image>Stacks>Tools>Concatenate’ and combine the tifs in order
d. ‘Analyze>Tools>ROI Manager’
e. Use the oval tool to choose a ROI
f. ‘Image>Stacks>Plot Z-axis profile’, this allows you to see the intensity over time. Click live so this plot updates as you choose a new ROI.
g. Define ROIs that visually look like a neuron (bright intensity/circular morphology) and have a trace with a steep response during the ligand or highK^+^ response.
h. Add appropriate ROIs to the ROI manager, aim for ∼15 ROIs.
i. Within the ROI manager ‘More>Save…’ and save the ROIs in case you need to return to them later. Save in the ‘Neuropil_Filter’ folder in the main folder.
j. Within the ROI manager ‘More>Multi Measure’ and save the results table and make sure it ends in ‘_Neurons.csv’. Make sure each column is the mean intensity of an ROI and each row is a slice (Analyze>Set Measurements…). Save in the ‘Neuropil_Filter’ folder in the main folder.
k. Repeat for neuropil regions, keeping the ROI to approximately the same size as the ‘Neuron’ ROIs. Make sure the results file ends with ‘_Neuropil.csv’.
l. Repeat on all videos. If you have a large set of videos choosing a representative subset should be sufficient.

78. Run the ‘Octo_CaImaging_Part0_DetermineSlopeFilter.ipynb’ jupyter notebook

**Note:** This notebook takes the manually defined ‘Neuron’ and ‘Neuropil’ data saved in the

‘Neuropil_Filter’ folder and calculates/plots the max slope for all ROIs. The ‘Neuron’ ROIs should have a higher max slope compared to the Neuropil ROIs. It then saves the mean + standard deviation max slope for the ‘Neuropil’ ROIs in ‘Neuropil_Filter_max_slope_metrics_Normalised.xlsx’. This value is later used in step 79 to filter out ROIs. If you are working with 2P data you will not need to do this.

More details can be found in Courtney et al., 2025 (13).

79. Run the ‘Octo_CaImaging_Part1_InitialProcessing.ipynb’ jupyter notebook

**Note:** This notebook loads raw intensity traces from Suite2p, the user manually defines the anatomical regions, each ROI is normalized using min/max values, and filtered by the slope threshold calculated in step 77/78 above. It saves the ‘accepted’ traces, per anatomical region, for subsequent notebooks to use. All outputs will appear in the same folder as the video.

a. Define the main working directory containing all your data, and the folder containing the video you want to process.
b. Run the subsequent cells.
c. When the Napari GUI opens, manually define the anatomical regions by clicking ‘New shapes layer’, ‘Add polygons (P)’ and draw a shape around the outer granular layer (OGL), inner granular layer (IGL), and medulla of the optic lobe, as well as the central brain, if the cells are also in focus.

**Note:** If you are working with our sample dataset it will load the previously defined regions in the folder and you won’t need to do anything.

d) Add the names for each region to ‘CaImaging_PolygonAnnotation_V1.xlsx’ in the main folder. If you are working with our sample dataset this has already been done.
e) Run the subsequent cells.
f) Repeat for next video.

80. Run the ‘Octo_CaImaging_Part2_Categorisation.ipynb’ jupyter notebook

**Note:** This notebook loads the filtered data from above and categorizes each ROI’s ligand response as excitation (20% above baseline), inhibition (20% below baseline) or no response. It also generates heatmaps and other visualisations. All outputs will appear in the same folder as the video.

a. Define the main working directory containing all your data, and the folder containing the video you want to process.
b. Run the subsequent cells.
c. Repeat for next video.

81. Run the ‘Octo_CaImaging_Part3_ExcitatorySlope.ipynb’ jupyter notebook

**Note:** This notebook loads all excitatory responses and calculates the slope of the response. In our analysis this was used for videos involving dopamine stimulation, acetylcholine stimulation didn’t show many excitatory responses. All outputs will appear in the same folder as the video.

a. Define the main working directory containing all your data, and the folder containing the video you want to process.
b. Run the subsequent cells.
c. Repeat for next video.

**Note:** Once steps 79-81 have been run on all videos proceed to step 82 to combine all data.

82. Run the ‘Octo_CaImaging_Part4_CompilingData.ipynb’ jupyter notebook

**Note:** this notebook compiles all dopamine and acetylcholine data, across all videos, and plots the results.

a. Define the main folder with all the folders containing videos and where you want the outputs to be saved.
b. Cell 3, 7, and 11 will load all data from folders containing ‘Dop’ or ‘ACh’, if you are working with our sample dataset don’t change anything, if you are running your own data change this to whatever string defines your group stimulus.

### Expected outcomes

Here we present a protocol to record and quantify the calcium activity from cells in *Octopus vulgaris* hatchling optic lobe slices during application of neurotransmitters. This protocol was applied in Courtney et al., 2025 (13) to understand the role dopamine and acetylcholine are playing in the *Octopus vulgaris* hatchling optic lobe. We specifically focused on cells within the three layers of the optic lobe; OGL, IGL and medulla. Figure 5A shows maximum intensity projections of five representative optic lobe slices from *Octopus vulgaris* hatchlings when the CAL-520 calcium indicator is imaged over time during application of dopamine or acetylcholine and highK^+^. When the user has become experienced with the brain dissection and slice preparation, they will be able to achieve a fully in-focus optic lobe resembling these example images, with single cells clearly visible. During imaging, we first apply octomedia and regularly observed spontaneous activity in many cells. We then apply either dopamine (Figure 4A) or acetylcholine (Figure 5A/B), we then apply octomedia again and then finally the imaging protocol ends with robust activation of all alive cells using a highK^+^ solution. During dopamine application, we observe robust excitatory responses (at least 20% above baseline) in all layers of the optic lobe. When we quantify the proportion of cells that were excited by dopamine, we generally see higher proportions of excitatory responses in the IGL and medulla compared to the OGL. Data across multiple slices is presented in Courtney et al., 2025 (13) and a representative slice is presented here in Figure 5A. For the experiments involving acetylcholine we previously observed that most cells either have no response to acetylcholine or they are inhibited by acetylcholine (at least 20% below baseline). For the results across multiple slices see Courtney et al., 2025 (13) and here we present two representative slices, one with mainly no response to acetylcholine (Figure 6A) and another with mainly inhibitory responses to acetylcholine (Figure 6B).

**Figure 5.**
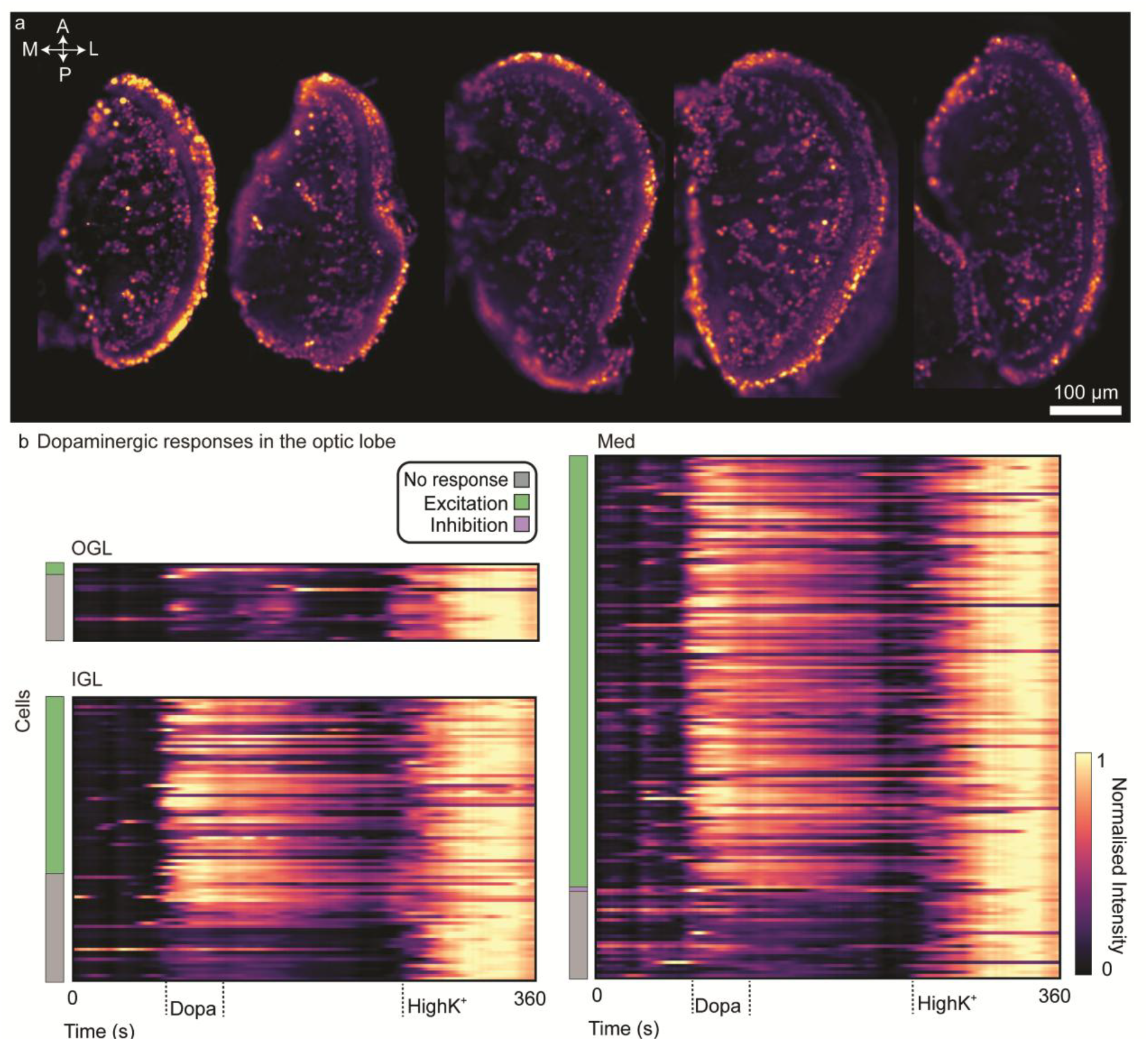
Expected results for calcium imaging of acute brain slices (optic lobe) of *Octopus vulgaris* hatchlings during application of 100 µM dopamine. (**A**) Maximum intensity projection of *Octopus vulgaris* hatchling brain slice (optic lobe) stained with CAL-520 (calcium indicator) and imaged over the course of 6 minutes during application of 100 µM dopamine or acetylcholine and highK^+^ solution, after registration using suite2p. (**B**) A heat map of a representative slice during calcium imaging over time when exposed to 100 µM dopamine and highK^+^ solutions. The lines correspond to individual cells (ROI) in the indicated layer of the optic lobe, and are ordered by ligand response category. The intensity for each ROI was smoothed using a rolling average window over 10 seconds and normalized using min/max values per ROI. A: anterior, P: posterior, M: medial, L: lateral, Dopa: dopamine, OGL: outer granular layer (of the optic lobe), IGL: inner granular layer (of the optic lobe), Med: medulla (of the optic lobe).

**Figure 6.**
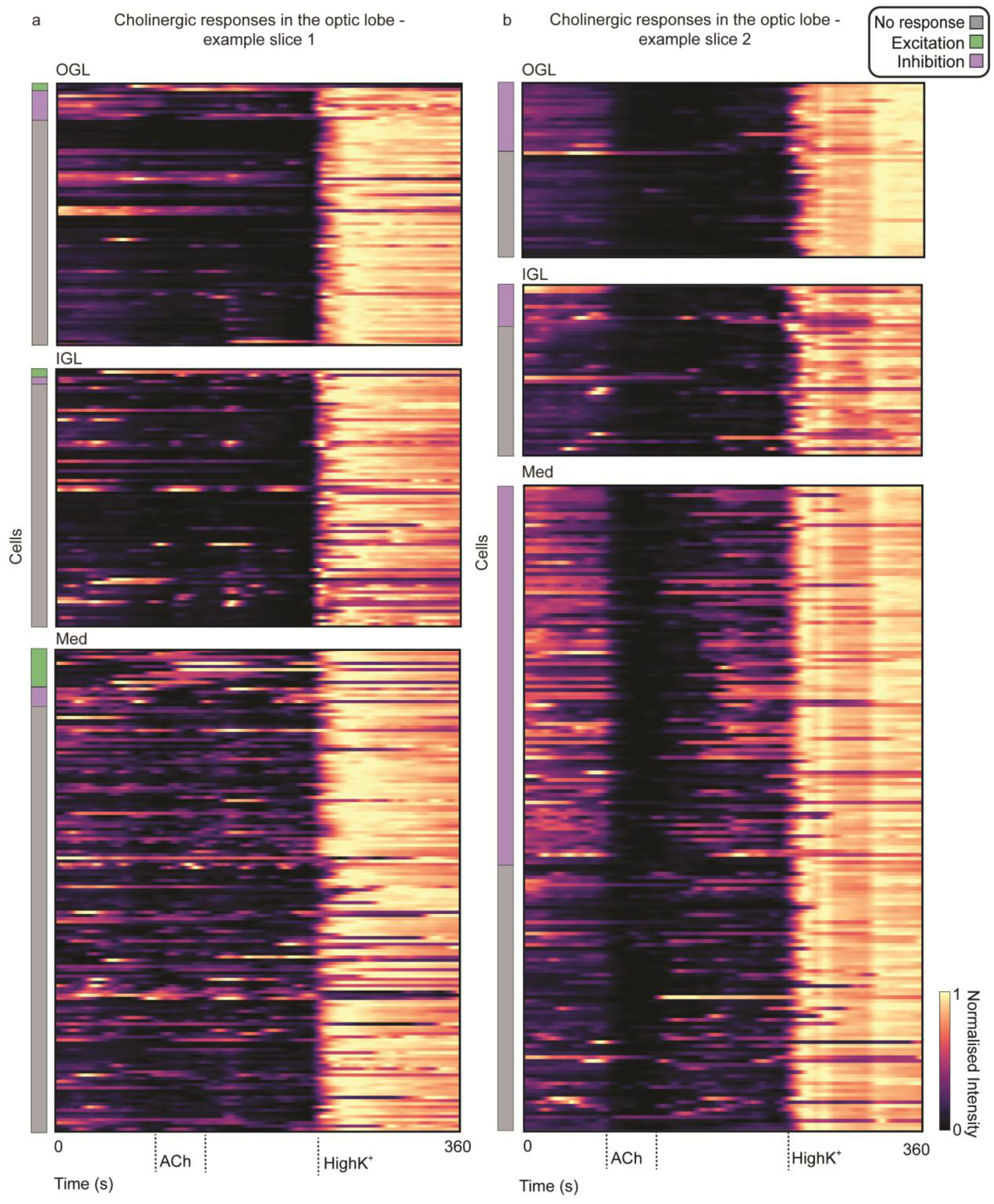
Expected results for calcium imaging of acute brain slices (optic lobe) of *Octopus vulgaris* hatchlings during application of 100 µM acetylcholine. (**A**) Heat map of a representative brain slice (optic lobe), with lines corresponding to individual cells in the indicated layer, and ordered by ligand response category. The fluorescence intensity for each ROI was smoothed using a rolling average window over 10 second periods and normalized using min/max values per ROI. This slice predominantly displayed no response to acetylcholine, with very little excitation or inhibition. (**B**) Same as A, but this slice mainly displayed inhibition across all layers when acetylcholine was applied. ACh: acetylcholine.

### Limitations

While this protocol has enabled us to observe the functional roles of dopamine and acetylcholine in the *Octopus vulgaris* hatchling optic lobe, and has immense potential as a tool for asking further questions about the neurochemical signaling mechanisms in the brains of these animals, several limitations exist that could be improved in the future. First, in inexperienced hands, the quality of the brain slices are unreliable, however, this improves quickly as the experimenter gains experience. The factors that contribute to low quality slices are discussed in the troubleshooting section below. Secondly, this protocol needs to be performed within a single day, to ensure optimal brain slice health. You therefore need two experimenters; one to prepare the slices early in the day and a second to stay later into the evening and do the imaging. This limitation could be overcome by maintaining the brain slices over multiple days, however, optimization and assessment of viability of the tissue would need to be carried out first. Thirdly, each stimulation with either dopamine or acetylcholine was performed on separate slices, as we didn’t want the impact of the first stimulus to affect the response of the second one. You could potentially reuse slices after a sufficient recovery period, however we didn’t test this. Another limitation involves the use of an epifluorescent microscope with an upright dry objective and closed imaging chamber. While this provided sufficient performance for our experiments, and offers a readily accessible setup, it is not the optimal configuration. Two-photon microscopy remains the gold standard and would improve optical sectioning, signal-to-noise, and minimize background contamination from out of focus cells. Another thing to consider is that this protocol involves applying the ligands to the entire bath and thus all cells are stimulated almost simultaneously. We therefore are observing the activity of both the primary activating neurons (which express receptors for these molecules) and the secondary activating neurons that are downstream of the primary activating neurons. Future optimization could tease this relationship apart by looking at the response onset timings, by applying the neurotransmitters locally, or by applying pharmacological agents (such as tetrodotoxin) to selectively inhibit synaptic signaling. Another limitation relates to the central brain lobes. While we have reliably imaged from the optic lobes, we haven’t yet established a robust approach for identifying and reliably recording from specific lobes within the central brain. *Octopus vulgaris* adults have ∼40 central brain lobes (28) which also appear to be present at hatching. This could be overcome through post-imaging fixation and morphological or molecular identification of the central brain lobes. The final limitation is that we do not know the molecular identity of the individual cells within different layers of the optic lobe that we have recorded from. Performing cell-specific knock-ins are currently not possible in *Octopus vulgaris* hatchlings (or other cephalopods).However fluorescent *in situ* hybridization chain reaction (HCR) is routinely used to spatially localize the molecular identity of cells in *Octopus vulgaris* hatchlings (7,13). Future optimization could involve coupling calcium imaging with subsequent HCR to link the functional and molecular identity of neurons.

### Troubleshooting

#### Problem 1: Tissue popping out of agarose during vibratome sectioning

During vibratome sectioning, the tissue may sometimes detach from the agarose. Because the tissue is very small, the affected slice is then lost. Keeping the brain embedded within the surrounding agarose is important for maintaining slice orientation and for immobilization during imaging.

##### Potential solutions

- We tested multiple types of agarose and found low gelling temperature (26°C) and high gel strength (500 g/cm^2^) were critical. Use the agarose listed in the key resources table.
- Air bubbles in the agarose should be prevented. Prepare the agarose early in the day and keep it in a water bath to allow bubbles to dissipate slowly over time (steps 4/5).
- Residual liquid around the brain at the time of solidification increases the chance of detachment. Remove as much liquid as possible using the perforated foil method, and swirl the brain several times in the agarose before it solidifies (step 20).
- Start with the vibratome settings described here, but optimizing the speed and amplitude for your instrument may improve results (step 30).
- We found that 3% agarose worked well for our tissue, but for your tissue different concentrations could work better.

#### Problem 2: Many ‘dead’ or dying cells

Signs of an unhealthy slice include absence of spontaneous activity, weak or absent responses to ligands or highK^+^, and no fluorescence changes over time (only photobleaching).

##### Potential solutions

- Avoid excessive mechanical damage during dissection (step 12). Practice will improve consistency and tissue viability.
- Minimize the time that agarose-embedded brains are kept on ice. Test whether room temperature handling is sufficient (step 23).
- Keep the interval between dissection and imaging under 12 hours. If you go beyond this replace octomedia more frequently.

#### Problem 3: Spontaneous activity present but no response to stimulants

If spontaneous activity is visible but there is no response to ligands or highK^+^, consider the following solutions.

##### Potential solutions

- Check that perfusion valves are not blocked and that flow rate is adequate (step 40b).
- Ensure the perfusion inflow tubing is not excessively long, which can delay solution exchange at the slice (step 40c).
- Ensure that the ligands are fully dissolved.

#### Problem 4: Few cells in focus within a single plane

Ideally, both optic lobes and central brain should be in focus. In some cases, only part of the brain is visible in a single focal plane.

##### Potential solutions

- Improve brain dissection technique to minimize tissue distortion or damage (step 12).
- Optimize vibratome settings (step 30).
- Avoid movement of the brain/agarose slice on the circular coverslip after vibratome sectioning.
- Confirm that imaging is performed from the cut surface of the slice (step 32).

#### Problem 5: Low signal-to-noise ratio during calcium imaging

Low signal-to-noise makes fluorescence changes difficult to quantify at the single-cell level.

##### Potential solutions

- For 200 µm slices, incubation in CAL-520 provided adequate signal. For thicker sections, or tissue from different species/organs, they may require deeper dye delivery (e.g. pressure injection as described in Pungor et al., 2023 (14)), longer incubation, and or higher dye concentration.
- Minimize photobleaching by keeping slices in the dark whenever possible.
- Ensure that the CAL-520 stock has not degraded over time.

## Resource availability

### Lead Contact

Further information and requests for any resources and reagents should be directed to and will be fulfilled by the lead contact, Eve Seuntjens.

### Technical contact

Questions regarding technical specifics of the protocol should be directed to the technical contact, Amy Courtney.

### Materials availability

This study did not generate new unique reagents.

## Data and code availability

The data can be found here: 10.5281/zenodo.18981973. The code is available here: https://github.com/A-Courtney/Octo_Dop_ACh_Ca_Imaging_2025 and in supplemental file 3.

## Acknowledgments

The authors gratefully acknowledge Denise Walker and all other past and present members of the Schafer and Seuntjens labs for technical assistance and helpful discussions. The authors also wish to thank the Arckens lab for kindly lending us equipment. The authors wish to thank Joshua Rosenthal and the Cephalopods Program at the Marine Biological Laboratory for their support in protocol development. The authors wish to thank all support and administrative staff at the LMB and KU Leuven. This work was kindly supported by the Medical Research Council MC-A023-5PB91 (to W.R.S.), the Research Foundation – Flanders G079521N (to W.R.S.), G040124N (to E.S.) and fellowship 1181025N (to M.V.D.), and KU Leuven BOF C14/16/049 (to E.S. and W.R.S.) and ID-N/20/007 (to E.S.).IEO was funded by the European Maritime, Fisheries and Aquaculture Fund.

## Author contributions

A.C., R.S., W.R.S and E.S., conceptualized the project. E.A. supplied the *Octopus vulgaris* embryos. A.C., M.V.D., R.S., and E.S. optimized the calcium imaging experiments, while A.C. and M.V.D. carried out the experiments and documented the experimental workflow. A.C. and H.A.O. developed the calcium image analysis pipeline, while A.C. carried out the analysis and prepared all figures. A.C. wrote the original manuscript, and all authors read and critically revised the manuscript. W.R.S and E.S. supervised the project and provided funding/infrastructure.

