## Supplemental file 3 for "Protocol for calcium imaging of acute brain slices from *Octopus vulgaris* hatchlings during application of neurotransmitters": environment.yml

```
name: Octo-Ca-Im
channels:
  - conda-forge
  - defaults
  - https://repo.anaconda.com/pkgs/main
  - https://repo.anaconda.com/pkgs/r
  - https://repo.anaconda.com/pkgs/msys2
dependencies:
  - _libavif_api=1.1.1
  - alabaster=0.7.16
  - annotated-types=0.7.0
  - anyio=4.7.0
  - aom=3.9.1
  - app-model=0.3.0
  - appdirs=1.4.4
  - argon2-cffi=23.1.0
  - argon2-cffi-bindings=21.2.0
  - arrow=1.3.0
  - asciitree=0.3.3
  - asttokens=3.0.0
  - async-lru=2.0.4
  - attrs=24.3.0
  - babel=2.16.0
  - beautifulsoup4=4.12.3
  - blas=1.0
  - bleach=6.2.0
  - blosc=1.21.3
  - bottleneck=1.4.2
  - brotli=1.0.9
  - brotli-bin=1.0.9
  - brotli-python=1.0.9
  - bzip2=1.0.8
  - ca-certificates=2024.12.14
  - cached-property=1.5.2
  - cached_property=1.5.2
  - cachey=0.2.1
  - certifi=2024.12.14
  - cffi=1.17.1
  - cfitsio=3.470
  - charls=2.2.0
  - charset-normalizer=3.4.0
  - cloudpickle=3.1.0
  - cmasher=1.8.0
  - colorama=0.4.6
  - colorspacious=1.1.2
  - comm=0.2.2
  - contourpy=1.2.0
  - cpython=3.9.21
  - cycler=0.11.0
  - cytoolz=1.0.1
  - dask-core=2024.8.0
  - dav1d=1.2.1
  - debugpy=1.8.11
  - decorator=5.1.1
  - defusedxml=0.7.1
  - docstring_parser=0.16
  - docutils=0.21.2
  - entrypoints=0.4
  - exceptiongroup=1.2.2
  - executing=2.1.0
  - fasteners=0.19
  - flexcache=0.3
  - flexparser=0.4
  - fonttools=4.51.0
  - fqdn=1.5.1
  - freetype=2.12.1
  - freetype-py=2.5.1
  - giflib=5.2.2
  - h11=0.14.0
  - h2=4.1.0
  - heapdict=1.0.1
  - hpack=4.0.0
  - hsluv=5.0.4
  - httpcore=1.0.7
  - httpx=0.28.1
  - hyperframe=6.0.1
  - icc_rt=2022.1.0
  - icu=73.1
  - idna=3.10
  - imagecodecs=2024.9.22
  - imageio=2.36.1
  - imagesize=1.4.1
  - importlib-metadata=8.5.0
  - importlib_metadata=8.5.0
  - importlib_resources=6.4.0
  - in-n-out=0.2.1
  - intel-openmp=2023.1.0
  - ipykernel=6.29.5
  - ipython=8.18.1
  - isoduration=20.11.0
  - jedi=0.19.2
  - jinja2=3.1.4
  - joblib=1.4.2
  - jpeg=9e
  - json5=0.10.0
  - jsonpointer=3.0.0
  - jsonschema=4.23.0
  - jsonschema-specifications=2024.10.1
  - jsonschema-with-format-nongpl=4.23.0
  - jupyter-lsp=2.2.5
  - jupyter_client=8.6.3
  - jupyter_core=5.7.2
  - jupyter_events=0.10.0
  - jupyter_server=2.14.2
  - jupyter_server_terminals=0.5.3
  - jupyterlab=4.3.3
  - jupyterlab_pygments=0.3.0
  - jupyterlab_server=2.27.3
  - kiwisolver=1.4.4
  - krb5=1.20.1
  - lazy-loader=0.4
  - lazy_loader=0.4
  - lcms2=2.16
  - lerc=4.0.0
  - libaec=1.1.3
  - libavif=1.1.1
  - libavif16=1.1.1
  - libbrotlicommon=1.0.9
  - libbrotlidec=1.0.9
  - libbrotlienc=1.0.9
  - libclang=14.0.6
  - libclang13=14.0.6
  - libdeflate=1.22
  - libpng=1.6.39
  - libpq=17.2
  - libsodium=1.0.18
  - libtiff=4.5.1
  - libwebp-base=1.3.2
  - libzopfli=1.0.3
  - llvmlite=0.43.0
  - locket=1.0.0
  - lz4-c=1.9.4
  - magicgui=0.9.1
  - markdown-it-py=3.0.0
  - markupsafe=3.0.2
  - matplotlib=3.9.2
  - matplotlib-base=3.9.2
  - matplotlib-inline=0.1.7
  - mdurl=0.1.2
  - mistune=3.0.2
  - mkl=2023.1.0
  - mkl-service=2.4.0
  - mkl_fft=1.3.11
  - mkl_random=1.2.8
  - msgpack-python=1.1.0
  - napari=0.5.5
  - napari-base=0.5.5
  - napari-console=0.1.3
  - napari-plugin-engine=0.2.0
  - napari-plugin-manager=0.1.4
  - napari-svg=0.2.0
  - nbclient=0.10.1
  - nbconvert-core=7.16.4
  - nbformat=5.10.4
  - nest-asyncio=1.6.0
  - networkx=3.2.1
  - notebook-shim=0.2.4
  - npe2=0.7.7
  - numba=0.60.0
  - numcodecs=0.12.1
  - numexpr=2.10.1
  - numpydoc=1.8.0
  - openjpeg=2.5.2
  - openssl=3.4.0
  - overrides=7.7.0
  - packaging=24.2
  - pandas=2.2.3
  - pandocfilters=1.5.0
  - parso=0.8.4
  - partd=1.4.2
  - pickleshare=0.7.5
  - pillow=11.0.0
  - pint=0.24.4
  - pkgutil-resolve-name=1.3.10
  - platformdirs=4.3.6
  - ply=3.11
  - pooch=1.8.2
  - prometheus_client=0.21.1
  - prompt-toolkit=3.0.48
  - psutil=6.1.0
  - psygnal=0.11.1
  - pure_eval=0.2.3
  - pybind11-abi=5
  - pyconify=0.2
  - pycparser=2.22
  - pydantic=2.10.4
  - pydantic-compat=0.1.2
  - pydantic-core=2.27.2
  - pygments=2.18.0
  - pynndescent=0.5.10
  - pyopengl=3.1.6
  - pyparsing=3.2.0
  - pyproject_hooks=1.2.0
  - pyqt=5.15.10
  - pyqt5-sip=12.13.0
  - pysocks=1.7.1
  - python=3.9.21
  - python-build=1.2.2.post1
  - python-dateutil=2.9.0post0
  - python-dotenv=1.0.1
  - python-fastjsonschema=2.21.1
  - python-json-logger=2.0.7
  - python-tzdata=2023.3
  - python_abi=3.9
  - pytz=2024.1
  - pywavelets=1.6.0
  - pywin32=307
  - pywinpty=2.0.14
  - pyyaml=6.0.2
  - pyzmq=26.2.0
  - qt-main=5.15.2
  - qtconsole-base=5.6.1
  - qtpy=2.4.2
  - rav1e=0.6.6
  - referencing=0.35.1
  - requests=2.32.3
  - rfc3339-validator=0.1.4
  - rfc3986-validator=0.1.1
  - rich=13.9.4
  - rpds-py=0.22.3
  - scikit-image=0.24.0
  - scikit-learn=1.5.1
  - scipy=1.13.1
  - seaborn=0.13.2
  - send2trash=1.8.3
  - shellingham=1.5.4
  - sip=6.7.12
  - six=1.16.0
  - snappy=1.2.1
  - sniffio=1.3.1
  - snowballstemmer=2.2.0
  - soupsieve=2.5
  - sphinx=7.4.7
  - sphinxcontrib-applehelp=2.0.0
  - sphinxcontrib-devhelp=2.0.0
  - sphinxcontrib-htmlhelp=2.1.0
  - sphinxcontrib-jsmath=1.0.1
  - sphinxcontrib-qthelp=2.0.0
  - sphinxcontrib-serializinghtml=1.1.10
  - sqlite=3.45.3
  - stack_data=0.6.3
  - superqt=0.7.0
  - svt-av1=2.3.0
  - tabulate=0.9.0
  - tbb=2021.8.0
  - terminado=0.18.1
  - threadpoolctl=3.5.0
  - tinycss2=1.4.0
  - tomli=2.0.1
  - tomli-w=1.1.0
  - toolz=1.0.0
  - tornado=6.4.2
  - tqdm=4.66.5
  - traitlets=5.14.3
  - typer=0.15.1
  - typer-slim=0.15.1
  - typer-slim-standard=0.15.1
  - types-python-dateutil=2.9.0.20241206
  - typing-extensions=4.12.2
  - typing_extensions=4.12.2
  - typing_utils=0.1.0
  - tzdata=2024b
  - ucrt=10.0.22621.0
  - umap-learn=0.5.4
  - unicodedata2=15.1.0
  - uri-template=1.3.0
  - urllib3=2.2.3
  - vc=14.40
  - vc14_runtime=14.42.34433
  - vispy=0.14.3
  - vs2015_runtime=14.42.34433
  - wcwidth=0.2.13
  - webcolors=24.11.1
  - webencodings=0.5.1
  - websocket-client=1.8.0
  - win_inet_pton=1.1.0
  - winpty=0.4.3
  - wrapt=1.17.0
  - xz=5.4.6
  - yaml=0.2.5
  - zarr=2.18.2
  - zeromq=4.3.5
  - zfp=1.0.1
  - zipp=3.21.0
  - zlib=1.2.13
  - zlib-ng=2.0.7
  - zstandard=0.23.0
  - zstd=1.5.6
  - pip:
      - ansi2html==1.9.2
      - blinker==1.9.0
      - cellpose==3.1.0
      - click==8.1.7
      - dash==2.18.2
      - dash-core-components==2.0.0
      - dash-html-components==2.0.0
      - dash-table==5.0.0
      - et-xmlfile==2.0.0
      - fastremap==1.15.0
      - filelock==3.16.1
      - flask==3.0.3
      - fsspec==2024.10.0
      - ipyfilechooser==0.6.0
      - ipywidgets==8.1.5
      - itsdangerous==2.2.0
      - jupyter-dash==0.4.2
      - jupyterlab-widgets==3.0.13
      - mpmath==1.3.0
      - natsort==8.4.0
      - notebook==7.3.1
      - numpy==1.26.4
      - opencv-python-headless==4.10.0.84
      - openpyxl==3.1.5
      - pandas-flavor==0.6.0
      - patsy==1.0.1
      - pingouin==0.5.5
      - pip==26.0
      - plotly==5.24.1
      - qgrid==1.3.1
      - rastermap==1.0
      - retrying==1.3.4
      - roifile==2024.9.15
      - scanimage-tiff-reader==1.4.1.4
      - setuptools==80.10.2
      - statsmodels==0.14.4
      - suite2p==0.14.4
      - sympy==1.13.1
      - tenacity==9.0.0
      - tifffile==2024.8.30
      - torch==2.5.1
      - werkzeug==3.0.6
      - wheel==0.46.3
      - widgetsnbextension==4.0.13
      - xarray==2024.7.0
prefix: C:\ProgramData\miniconda3\envs\Octo-Ca-Im
```
