## Supplemental file 3 for "Protocol for calcium imaging of acute brain slices from *Octopus vulgaris* hatchlings during application of neurotransmitters": Octo_CaImaging_Part0_DetermineSlopeFilter.ipynb

```
# %% [code]
import os
import numpy as np
import pandas as pd
import matplotlib.pyplot as plt
import seaborn as sns
from scipy.signal import savgol_filter
from scipy.stats import ttest_ind

master_folder = r"C:/Users/Amy Courtney/Documents/Temp/Calcium_Imaging_Paper/Biorxiv_Submission1_Mar26/Supp_Files/Courtney_et_al_2026_Supp_File_ 3_Sample_Dataset/Calcium_Imaging_Analysis/Neuropil_Filter"

if not os.path.isdir(master_folder):
    raise FileNotFoundError(f"master_folder does not exist: {master_folder}")

lowest_folder_name = os.path.basename(os.path.normpath(master_folder))

# Fixed parameters
FR = 5
Fil_Win = 50
Sp_Win = 10

def dataset_id_from_filename(filename: str) -> str:
    if filename.endswith("_Neuropil.csv"):
        return filename[:-len("_Neuropil.csv")]
    if filename.endswith("_Neurons.csv"):
        return filename[:-len("_Neurons.csv")]
    return os.path.splitext(filename)[0]

# Find all CSV files ending in _Neuropil and _Neurons
files = []
for root, _, filenames in os.walk(master_folder):
    for filename in filenames:
        if filename.endswith("_Neuropil.csv") or filename.endswith("_Neurons.csv"):
            group = "Neuropil" if filename.endswith("_Neuropil.csv") else "Neurons"
            dsid = dataset_id_from_filename(filename)
            files.append((
                os.path.join(root, filename),
                group,
                dsid
            ))

# Sort for reproducibility
files = sorted(files, key=lambda x: x[0])

print(f"\nCSV files used from: {master_folder}\n")
for f, grp, dsid in files:
    print(f"{grp}: {f}   Dataset: {dsid}")

print(
    f"\nTotal CSV files used: {len(files)} "
    f"({sum(g == 'Neuropil' for _, g, _ in files)} Neuropil, "
    f"{sum(g == 'Neurons' for _, g, _ in files)} Neurons)\n"
)

input_files_df = pd.DataFrame(files, columns=["File_Path", "Group", "Dataset_ID"])

# Prepare data
features = []

for file, group, dsid in files:
    data = pd.read_csv(file)

    # Keep only columns that include 'Mean'
    mean_cols = [col for col in data.columns if "Mean" in col]
    data = data[mean_cols]

    if data.shape[1] == 0:
        print(f"WARNING: No 'Mean' columns found in {file}. Skipping.")
        continue

    # Normalize each trace (column-wise), protect against flat traces
    denom = (data.max() - data.min()).replace(0, np.nan)
    data = (data - data.min()) / denom
    data = data.fillna(0)

    # Window checks
    if Fil_Win >= len(data):
        print(f"WARNING: Fil_Win ({Fil_Win}) >= number of frames ({len(data)}) in {file}. Skipping.")
        continue

    # Savitzky-Golay filter
    smoothed_data = data.apply(
        lambda x: savgol_filter(x, window_length=Fil_Win, polyorder=3),
        axis=0
    )

    if Sp_Win >= len(smoothed_data):
        print(f"WARNING: Sp_Win ({Sp_Win}) >= number of frames ({len(smoothed_data)}) in {file}. Skipping.")
        continue

    slopes = []
    for start in range(0, len(smoothed_data) - Sp_Win):
        subset = smoothed_data.iloc[start:start + Sp_Win]
        slope = ((subset.iloc[-1] - subset.iloc[0]) / (len(subset) - 1)) * FR
        slopes.append(slope)

    if len(slopes) == 0:
        print(f"WARNING: Not enough frames to compute slopes in {file}. Skipping.")
        continue

    slopes_max = np.max(slopes, axis=0)

    features.append(pd.DataFrame({
        "Slope_Max": slopes_max,
        "Group": group,
        "Dataset_ID": dsid,
        "Source_File": os.path.basename(file)
    }))

if len(features) == 0:
    raise RuntimeError("No valid data found to analyze. Check your CSV files and parameters.")

features = pd.concat(features).reset_index(drop=True)

# Split groups for global summary
neuropil_features = features[features["Group"] == "Neuropil"]
neuron_features = features[features["Group"] == "Neurons"]

neu_mean = neuropil_features["Slope_Max"].mean()
neu_std = neuropil_features["Slope_Max"].std()

# Output paths: all into master_folder
output_tiff_path = os.path.join(
    master_folder,
    f"{lowest_folder_name}_max_slope_distribution_plot_Normalised.tiff"
)
output_pdf_path = os.path.join(
    master_folder,
    f"{lowest_folder_name}_max_slope_distribution_plot_Normalised.pdf"
)
excel_output_path = os.path.join(
    master_folder,
    f"{lowest_folder_name}_max_slope_metrics_Normalised.xlsx"
)

# Global plot
fig, ax = plt.subplots(figsize=(6, 5))
sns.boxplot(x="Group", y="Slope_Max", data=features, showfliers=False, ax=ax)
sns.stripplot(x="Group", y="Slope_Max", data=features, color="black", alpha=0.5, size=5, ax=ax)

ax.axhline(neu_mean + neu_std, linestyle=":", linewidth=1.5, label=f"+1 SD = {(neu_mean + neu_std):.3f}")
ax.set_title("Max Slope")
ax.set_xlabel("Group")
ax.set_ylabel("Slope")
ax.grid(True)
ax.legend(frameon=False, fontsize=9)

plt.tight_layout()
plt.savefig(output_tiff_path, format="tiff", dpi=300)
plt.savefig(output_pdf_path, format="pdf")
plt.close()

# Global statistics
if len(neuropil_features) == 0 or len(neuron_features) == 0:
    p_value = np.nan
    significance = "n/a"
    print("WARNING: One or both groups are empty. t-test not performed.")
else:
    ttest = ttest_ind(neuropil_features["Slope_Max"], neuron_features["Slope_Max"])
    p_value = ttest.pvalue
    significance = "***" if p_value <= 0.001 else (
        "**" if p_value <= 0.01 else (
            "*" if p_value <= 0.05 else "n.s."
        )
    )

stat_summary = [{
    "Metric": "Slope_Max",
    "Neuropil_Mean": neu_mean,
    "Neuropil_Std": neu_std,
    "Neuropil_Mean_plus_Std": neu_mean + neu_std,
    "Neuropil_Mean_minus_Std": neu_mean - neu_std,
    "Neuron_Mean": neuron_features["Slope_Max"].mean() if len(neuron_features) > 0 else np.nan,
    "Neuron_Std": neuron_features["Slope_Max"].std() if len(neuron_features) > 0 else np.nan,
    "P_Value": p_value,
    "Significance": significance
}]

neuropil_summary = pd.DataFrame({
    "Metric": ["Slope_Max"],
    "Mean": [neu_mean],
    "Std": [neu_std],
    "Mean_plus_Std": [neu_mean + neu_std],
    "Mean_minus_Std": [neu_mean - neu_std]
})

# Per dataset plots folder (still inside master_folder)
per_dataset_plot_dir = os.path.join(master_folder, "Per_Dataset_Plots")
os.makedirs(per_dataset_plot_dir, exist_ok=True)

# Make one plot per dataset (Neurons vs Neuropil)
datasets = sorted(features["Dataset_ID"].unique())

per_dataset_stats = []

for dsid in datasets:
    df_ds = features[features["Dataset_ID"] == dsid].copy()
    if df_ds.empty:
        continue

    # Only plot if both groups exist for this dataset
    groups_present = set(df_ds["Group"].unique())
    if not ("Neurons" in groups_present and "Neuropil" in groups_present):
        print(f"WARNING: Dataset {dsid} does not contain both groups. Skipping per dataset plot.")
        continue

    # Stats per dataset
    neu_ds = df_ds[df_ds["Group"] == "Neuropil"]["Slope_Max"]
    neu_ds_mean = neu_ds.mean()
    neu_ds_std = neu_ds.std()

    neuN_ds = df_ds[df_ds["Group"] == "Neurons"]["Slope_Max"]
    if len(neu_ds) > 0 and len(neuN_ds) > 0:
        ttest_ds = ttest_ind(neu_ds, neuN_ds)
        p_ds = ttest_ds.pvalue
        sig_ds = "***" if p_ds <= 0.001 else (
            "**" if p_ds <= 0.01 else (
                "*" if p_ds <= 0.05 else "n.s."
            )
        )
    else:
        p_ds = np.nan
        sig_ds = "n/a"

    per_dataset_stats.append({
        "Dataset_ID": dsid,
        "Neuropil_Mean": neu_ds_mean,
        "Neuropil_Std": neu_ds_std,
        "Neurons_Mean": neuN_ds.mean(),
        "Neurons_Std": neuN_ds.std(),
        "P_Value": p_ds,
        "Significance": sig_ds
    })

    # Plot
    fig, ax = plt.subplots(figsize=(6, 5))
    sns.boxplot(x="Group", y="Slope_Max", data=df_ds, showfliers=False, ax=ax)
    sns.stripplot(x="Group", y="Slope_Max", data=df_ds, color="black", alpha=0.5, size=5, ax=ax)

    # Dotted lines: neuropil mean ± std for this dataset
    ax.axhline(neu_ds_mean + neu_ds_std, linestyle=":", linewidth=1.5,
               label=f"Neuropil +1 SD = {(neu_ds_mean + neu_ds_std):.3f}")

    ax.set_title(f"Max Slope: {dsid}")
    ax.set_xlabel("Group")
    ax.set_ylabel("Slope")
    ax.grid(True)
    ax.legend(frameon=False, fontsize=9)

    plt.tight_layout()

    out_tiff = os.path.join(per_dataset_plot_dir, f"{dsid}_max_slope_Normalised.tiff")
    out_pdf = os.path.join(per_dataset_plot_dir, f"{dsid}_max_slope_Normalised.pdf")

    plt.savefig(out_tiff, format="tiff", dpi=300)
    plt.savefig(out_pdf, format="pdf")
    plt.close()

# Write Excel, now including per dataset stats too
with pd.ExcelWriter(excel_output_path) as writer:
    features.to_excel(writer, sheet_name="Metrics", index=False)
    pd.DataFrame(stat_summary).to_excel(writer, sheet_name="Statistics", index=False)
    neuropil_summary.to_excel(writer, sheet_name="Neuropil_Summary", index=False)
    input_files_df.to_excel(writer, sheet_name="Input_CSV_Files", index=False)
    pd.DataFrame(per_dataset_stats).to_excel(writer, sheet_name="Per_Dataset_Stats", index=False)

print(f"Saved global plot (TIFF): {output_tiff_path}")
print(f"Saved global plot (PDF):  {output_pdf_path}")
print(f"Saved per dataset plots in: {per_dataset_plot_dir}")
print(f"Saved Excel: {excel_output_path}")
```
