## Supplemental file 3 for "Protocol for calcium imaging of acute brain slices from *Octopus vulgaris* hatchlings during application of neurotransmitters": Octo_CaImaging_Part1_InitialProcessing.ipynb

```
# %% [code]
import json
import os
import sys

import numpy as np
import napari
from PIL import Image
from pathlib import Path  # filesystem paths

import matplotlib.pyplot as plt
from matplotlib.colors import ListedColormap, BoundaryNorm
from matplotlib.backends.backend_pdf import PdfPages
from matplotlib.path import Path as MplPath  # matplotlib geometry Path (renamed)

import ipywidgets as widgets
from ipyfilechooser import FileChooser
from IPython.display import display

import seaborn as sns
import plotly.graph_objects as go

import yaml
import pandas as pd
import openpyxl

from scipy.ndimage import gaussian_filter1d
from scipy.signal import savgol_filter

# %% [code]
DEFAULT_MAIN = r"C:/Users\Amy Courtney\Documents/Temp/Calcium_Imaging_Paper/Biorxiv_Submission1_Mar26/Supp_Files/Courtney_et_al_2026_Supp_File_3_Sample_Dataset/Calcium_Imaging_Analysis"
DEFAULT_BASE = r"C:/Users\Amy Courtney\Documents/Temp/Calcium_Imaging_Paper/Biorxiv_Submission1_Mar26/Supp_Files/Courtney_et_al_2026_Supp_File_3_Sample_Dataset/Calcium_Imaging_Analysis"

main_fc = FileChooser(DEFAULT_MAIN, select_desc="Main Folder")
main_fc.show_only_dirs = True

base_fc = FileChooser(DEFAULT_BASE, select_desc="Single Video Folder")
base_fc.show_only_dirs = True

display(main_fc)
display(base_fc)


# %% [code]
if "matplotlib.path" in str(Path):
    raise ImportError(
        "Path is imported from matplotlib.path. "
        "Remove 'from matplotlib.path import Path' from your notebook."
    )

# ---- Validate folder chooser objects ----
if main_fc.selected_path is None:
    raise ValueError("No main folder selected.")

if base_fc.selected_path is None:
    raise ValueError("No base folder selected.")

# ---- Convert to pathlib paths safely ----
main_folder = Path(main_fc.selected_path).expanduser().resolve()
base_folder = Path(base_fc.selected_path).expanduser().resolve()

# ---- Validate existence ----
if not main_folder.exists():
    raise FileNotFoundError(f"Main folder does not exist: {main_folder}")

if not base_folder.exists():
    raise FileNotFoundError(f"Base folder does not exist: {base_folder}")

if not main_folder.is_dir():
    raise NotADirectoryError(f"Main path is not a directory: {main_folder}")

if not base_folder.is_dir():
    raise NotADirectoryError(f"Base path is not a directory: {base_folder}")

# ---- Print results ----
print("Main folder:", main_folder)
print("Base folder:", base_folder)

# %% [code]
suite2p_output = base_folder / 'suite2p/plane0'
suite2p_output
iscell = np.load(suite2p_output / 'iscell.npy')
ops = np.load(suite2p_output / 'ops.npy', allow_pickle= True).item()
fluos = np.load(suite2p_output / 'F.npy')
neurops = np.load(suite2p_output / 'Fneu.npy')
stats = np.load(suite2p_output / 'stat.npy', allow_pickle= True)
print(fluos.shape,iscell.shape)

# %% [code]
fluos_df = pd.DataFrame(fluos).T
fluos_df.columns = [f'ROI{i}' for i in range(fluos_df.shape[1])]
fluos_df.index = range(1, fluos_df.shape[0] + 1)  # Set index to 1, 2, ..., n
# Keep only rows 1 to 1600
#fluos_df = fluos_df.iloc[1:1600]

# Assuming `iscell` is a NumPy array and the first column is what you want to check
fluos_df = fluos_df.loc[:, iscell[:, 0] == 1.0]


# %% [code]
folder_name = os.path.basename(os.path.normpath(base_folder))

# Plot all traces on one plot
plt.figure(figsize=(10, 6))
plt.style.use('default')

# Plot all traces
for column in fluos_df.columns:
    plt.plot(fluos_df[column], alpha=0.8, linewidth=0.6)

# Add axis labels and title
plt.xlabel('Frame', fontsize=10, labelpad=8)
plt.ylabel('Fluorescence Intensity', fontsize=10, labelpad=8)
plt.title('Fluorescence Traces', fontsize=12, weight='bold')

# Improve axis and visibility
plt.grid(False)
plt.gca().spines['top'].set_visible(False)
plt.gca().spines['right'].set_visible(False)

# Layout
plt.tight_layout()

# Save using your folder convention
save_path = os.path.join(base_folder, f"{folder_name}_Fluorescence_Traces.pdf")
plt.savefig(save_path, dpi=300, bbox_inches='tight')

# Show the plot
plt.show()


# %% [code]
# Plot specific trace by column name
trace_name = 'ROI10'  # Replace this with the desired column name
plt.figure(figsize=(10, 6))  # Larger figure size
plt.style.use('default')  # Use default style for white background

# Plot the specified trace
plt.plot(fluos_df[trace_name], alpha=0.8, linewidth=0.6, label=trace_name)

# Add axis labels and title
plt.xlabel('Frame', fontsize=10, labelpad=8)  # Smaller font size and padding
plt.ylabel('Fluorescence Intensity', fontsize=10, labelpad=8)
plt.title(f'Fluorescence Trace - {trace_name}', fontsize=12, weight='bold')


# Improve axis and visibility
plt.grid(False)  # Turn off grid lines
plt.gca().spines['top'].set_visible(False)  # Remove top border
plt.gca().spines['right'].set_visible(False)  # Remove right border

# Show the plot
plt.tight_layout()
plt.show()

# %% [code]
stats = np.load(suite2p_output / 'stat.npy', allow_pickle=True)
print(len(stats))
print(stats[0].keys())

# %% [code]
# Load stats data
stats = np.load(suite2p_output / 'stat.npy', allow_pickle=True)

# Create a DataFrame for specific keys
df_list = []
for idx, entry in enumerate(stats):
    # Extract the 'med' key and keep only its values
    flat_entry = {'med': entry['med'], 'Index': idx}  # Keep index for reference
    df_list.append(flat_entry)

# Convert the list of dictionaries into a DataFrame
df_stats = pd.DataFrame(df_list)

# Extract 'med' column into a new DataFrame
ROI_Location = pd.DataFrame(df_stats['med'].tolist(), columns=['x', 'y'])  # Split 'med' into 'x' and 'y'

image = ops['max_proj']

# Extract x and y coordinates
x_coords = ROI_Location['x']
y_coords = ROI_Location['y']

# Plot the image and overlay ROI locations
plt.figure(figsize=(10, 10))
plt.imshow(image)
plt.scatter(y_coords, x_coords, color='red', s=15, label='ROI')
plt.grid(True, color='blue', linestyle='--', linewidth=0.5)  # Add grid for debugging
plt.legend()
plt.show()


# %% [code]
#ONLY RUN IF WANT TO DEFINE ANATOMICAL REGIONS

# Load the image and convert to NumPy array
nuclei = ops['max_proj']

# Define variables
resolution_level = 1  # Replace with the actual resolution level if needed

# Get the directory of the image and set region file path

regionname = os.path.join(base_folder, "regions.json")  # Save JSON in the same directory as the image

# Open the image in Napari
v = napari.view_image(nuclei, colormap='gray')  # Open the image in grayscale

# If region data exists, load and display it
try:
    with open(regionname, "r") as outfile:
        poly = json.load(outfile)
    c = pow(2, poly['resolution_level'] - resolution_level)
    sdata = [c * np.array(p) for p in poly['regions']]
    v.add_shapes(sdata, shape_type='polygon', edge_color='yellow')  # Display saved regions
except FileNotFoundError:
    print(f"No saved regions found in {regionname}.")

# Launch Napari viewer
napari.run()


# %% [code]
#ONLY RUN IF WANT TO DEFINE ANATOMICAL REGIONS

#save regions with napari still open
poly = {
    'shape': nuclei.shape,
    'resolution_level': resolution_level,
    'regions': [ p.tolist() for p in v.layers[-1].data ] 
}

with open(regionname, "w") as outfile:
    json.dump(poly, outfile)

# %% [code]
# Create regions_path by appending '/regions.json'
regions_path = os.path.join(base_folder, 'regions.json')

# Load regions data from JSON
with open(regions_path, 'r') as file:
    regions_data = json.load(file)

# Plot the image and polygons
plt.figure(figsize=(10, 10))
plt.imshow(image)

# Iterate through all regions in the JSON file
for idx, region in enumerate(regions_data['regions']):
    # Extract polygon vertices
    polygon_vertices = region  # Updated to treat region as a list of vertices

    # Close the polygon by adding the first point at the end
    polygon_vertices.append(polygon_vertices[0])  # Append the first point to close the loop

    # Separate into x and y coordinates
    polygon_x, polygon_y = zip(*polygon_vertices)

    # Plot the polygon
    plt.plot(polygon_y, polygon_x, linewidth=2, label=f'Polygon {idx + 1}')
    

# %% [code]
# ------------------------------------------------------------
# 1) Save ROI locations to Excel
# ------------------------------------------------------------
ROI_Location = ROI_Location.copy()
ROI_Location["name"] = [f"ROI{i}" for i in range(len(ROI_Location))]

lowest_folder = os.path.basename(os.path.normpath(str(base_folder)))
file_name = f"{lowest_folder}_ROIxy.xlsx"
file_path = os.path.join(str(base_folder), file_name)
ROI_Location.to_excel(file_path, index=False)

# ------------------------------------------------------------
# 2) Prepare outputs
# ------------------------------------------------------------
polygon_ROIs = {}

pdf_file_name = os.path.join(str(base_folder), f"{lowest_folder}_ROIs_within_Polygons.pdf")
pdf_pages = PdfPages(pdf_file_name)

num_regions = len(regions_data["regions"])
nrows = (num_regions + 1) // 2
fig, axes = plt.subplots(nrows=nrows, ncols=2, figsize=(15, max(1, num_regions) * 3))
axes = np.array(axes).flatten()

# Make sure coordinates are numpy arrays (handles pandas Series too)
x_arr = np.asarray(x_coords)
y_arr = np.asarray(y_coords)
roi_names = ROI_Location["name"].to_numpy()

# Points for contains_points must match the polygon coordinate convention.
# Your code swapped polygon vertices to (y, x) and tested points as (y, x),
# so we keep everything consistently (y, x) here.
roi_points_yx = np.column_stack((y_arr, x_arr))

# ------------------------------------------------------------
# 3) Iterate polygons, classify ROIs, plot
# ------------------------------------------------------------
for idx, region in enumerate(regions_data["regions"]):
    # Copy vertices so we do not modify the JSON data in-place
    verts = [tuple(p) for p in region]  # expects [(x, y), (x, y), ...]

    # Close polygon safely (only if not already closed)
    if len(verts) > 0 and verts[0] != verts[-1]:
        verts = verts + [verts[0]]

    # Swap to (y, x) to match your ROI coordinate system
    polygon_vertices_yx = [(y, x) for (x, y) in verts]

    # Matplotlib polygon path (geometry)
    polygon_path = MplPath(polygon_vertices_yx)

    # Vectorized point-in-polygon test
    inside_mask = polygon_path.contains_points(roi_points_yx)
    inside_indices = np.where(inside_mask)[0]
    inside_names = roi_names[inside_indices].tolist()

    polygon_ROIs[f"Polygon {idx + 1}"] = inside_names

    # Plot
    ax = axes[idx]
    ax.imshow(image)
    ax.scatter(
        y_arr[inside_indices],
        x_arr[inside_indices],
        color="magenta",
        s=15,
        label="Inside ROI",
    )
    ax.plot(*zip(*polygon_vertices_yx), color="white", linewidth=2, label=f"Polygon {idx + 1}")
    ax.grid(True, color="blue", linestyle="--", linewidth=0.5)
    ax.set_title(f"Polygon {idx + 1} ROIs")

# Hide unused subplots
for ax in axes[num_regions:]:
    ax.axis("off")

# ------------------------------------------------------------
# 4) Save and show
# ------------------------------------------------------------
plt.tight_layout()
pdf_pages.savefig(fig)
pdf_pages.close()

tiff_file_name = os.path.join(str(base_folder), f"{lowest_folder}_ROIs_within_Polygons.tiff")
fig.savefig(tiff_file_name, format="tiff")

plt.show()

# ------------------------------------------------------------
# 5) Print ROI names per polygon
# ------------------------------------------------------------
for polygon, rois in polygon_ROIs.items():
    print(f"{polygon}: {', '.join(rois)}")

# %% [code]
# Create a new DataFrame for ROI names and regions
roi_region_df = []
for polygon, rois in polygon_ROIs.items():
    for roi in rois:
        roi_region_df.append({'ID': roi, 'Region': polygon.replace('POLYGON ', 'Polygon').replace('Polygon ', 'Polygon')})

roi_region_df = pd.DataFrame(roi_region_df)

# Identify unique polygons in the Region column
unique_polygons = roi_region_df['Region'].unique()

# Create a dictionary to hold DataFrames for each polygon
polygon_dataframes = {}

# Iterate through each unique polygon and split the DataFrame
for polygon in unique_polygons:
    # Filter ROIs to include only those present in the fluorescence data
    polygon_rois = roi_region_df.loc[roi_region_df['Region'] == polygon, 'ID'].tolist()
    existing_rois = [roi for roi in polygon_rois if roi in fluos_df.columns]  # Keep only existing ROIs

    polygon_df = fluos_df[existing_rois] if existing_rois else pd.DataFrame()  # Create DataFrame if ROIs exist
    polygon_dataframes[polygon] = polygon_df

    # Debugging output
    print(f"{polygon} DataFrame shape: {polygon_df.shape}")

# Save each polygon DataFrame as a variable for later use
for polygon, df in polygon_dataframes.items():
    if not df.empty:
        globals()[f"{polygon}_df"] = df
        print(f"Saved DataFrame for {polygon} as a variable.")

# Plot fluorescence data for each polygon
for polygon, df in polygon_dataframes.items():
    if df.empty:
        print(f"No valid data for {polygon}.")
        continue

    plt.figure(figsize=(12, 8))
    plt.title(f'Fluorescence Traces - {polygon}')

    for roi in df.columns:
        plt.plot(df[roi], label=roi, alpha=0.6, linewidth=0.8)

    plt.xlabel('Frame')
    plt.ylabel('Fluorescence Intensity')
    plt.grid(False)
    plt.gca().spines['top'].set_visible(False)
    plt.gca().spines['right'].set_visible(False)
    plt.tight_layout()
    plt.show()


# %% [code]
# ------------------------------------------------------------
# 1) Build ROI → Region DataFrame
# ------------------------------------------------------------
roi_region_df = []

for polygon, rois in polygon_ROIs.items():
    for roi in rois:
        roi_region_df.append({
            "ID": roi,
            "Region": polygon.replace("POLYGON ", "Polygon").replace("Polygon ", "Polygon ")
        })

roi_region_df = pd.DataFrame(roi_region_df)

# ------------------------------------------------------------
# 2) Split fluorescence data per polygon
# ------------------------------------------------------------
unique_polygons = roi_region_df["Region"].unique()
polygon_dataframes = {}

for polygon in unique_polygons:
    polygon_rois = roi_region_df.loc[
        roi_region_df["Region"] == polygon, "ID"
    ].tolist()

    existing_rois = [roi for roi in polygon_rois if roi in fluos_df.columns]

    polygon_df = fluos_df[existing_rois] if existing_rois else pd.DataFrame()
    polygon_dataframes[polygon] = polygon_df

    print(f"{polygon} DataFrame shape: {polygon_df.shape}")

# ------------------------------------------------------------
# 3) Save plots to PDF
# ------------------------------------------------------------
lowest_folder = os.path.basename(os.path.normpath(str(base_folder)))
pdf_output_path = os.path.join(
    str(base_folder),
    f"{lowest_folder}_Fluorescence_Traces_by_Polygon.pdf"
)

with PdfPages(pdf_output_path) as pdf:

    for polygon, df in polygon_dataframes.items():

        if df.empty:
            print(f"No valid data for {polygon}.")
            continue

        fig = plt.figure(figsize=(12, 8))
        plt.title(f"Fluorescence Traces - {polygon}")

        for roi in df.columns:
            plt.plot(df[roi], alpha=0.6, linewidth=0.8)

        plt.xlabel("Frame")
        plt.ylabel("Fluorescence Intensity")
        plt.grid(False)
        plt.gca().spines["top"].set_visible(False)
        plt.gca().spines["right"].set_visible(False)

        plt.tight_layout()
        pdf.savefig(fig)   # Save current figure to PDF
        plt.close(fig)     # Important to prevent memory buildup

print(f"Fluorescence plots saved to:\n{pdf_output_path}")

# %% [code]
# Define base_folder explicitly
lowest_folder_name = os.path.basename(os.path.normpath(base_folder))

# Load the file into a DataFrame
full_file_path = os.path.join(main_folder, "CaImaging_PolygonAnnotation_V1.xlsx" )
try:
    df = pd.read_excel(full_file_path)
except FileNotFoundError:
    print(f"File not found: {full_file_path}")
    exit()

# Check if the 'Folder' column exists
if 'Folder' not in df.columns:
    print("The 'Folder' column is not in the file.")
    exit()

# Filter rows where the 'Folder' column matches the lowest folder name
matching_rows = df[df['Folder'] == lowest_folder_name]

if matching_rows.empty:
    print(f"No rows in the 'Folder' column match the name: {lowest_folder_name}")
else:
    print(f"Rows with matching 'Folder' name ({lowest_folder_name}):")
    print(matching_rows)

    # Check all strings in the matching rows
    unique_strings = matching_rows.drop(columns=['Folder']).applymap(str).stack().unique()
    matching_columns = {string: [] for string in unique_strings}

    # Find all columns that contain each string WITHIN matching rows
    for string in unique_strings:
        for col in matching_rows.columns:
            if matching_rows[col].astype(str).str.contains(string, na=False).any():
                matching_columns[string].append(col)

    print("Columns containing strings from matching rows:")
    for string, columns in matching_columns.items():
        print(f"String '{string}' is found in columns: {', '.join(columns)}")

    # Track DataFrame names
    created_df_names = []

    # Combine DataFrames for strings found in two columns
    for string, columns in matching_columns.items():
        if len(columns) == 2:  # Only process cases with exactly two matching columns
            col1, col2 = columns

            # Retrieve corresponding DataFrames
            try:
                df1 = globals()[f"{col1}_df"]
                df2 = globals()[f"{col2}_df"]

                # Combine DataFrames by adding columns (not rows)
                combined_df = pd.concat([df1, df2], axis=1)

                # Assign the combined DataFrame a new name
                combined_df_name = f"{string}_df"
                globals()[combined_df_name] = combined_df
                created_df_names.append(combined_df_name)

                print(f"Created combined DataFrame: {combined_df_name}")

            except KeyError as e:
                print(f"Error: DataFrame for column {e} not found. Skipping combination.")

        elif len(columns) == 1:  # Handle unique strings
            col = columns[0]

            # Retrieve the corresponding DataFrame
            try:
                unique_df = globals()[f"{col}_df"]

                # Rename the DataFrame with the unique string
                unique_df_name = f"{string}_df"
                globals()[unique_df_name] = unique_df
                created_df_names.append(unique_df_name)

                print(f"Renamed DataFrame: {unique_df_name}")

            except KeyError:
                print(f"Error: DataFrame for column {col} not found. Skipping rename.")

        else:
            print(f"Skipping string '{string}' as it is found in {len(columns)} columns, not 2 or 1.")

    # Save the names of all created DataFrames
    all_created_dfs = created_df_names
    print("All created DataFrame names:", all_created_dfs)


# %% [code]
FR=5
Met_win=10
Mean_win=50
Fil_Win=50

# -------------------------------------------------------------------
# THRESHOLD LOADER: read Column D from Statistics sheet
# -------------------------------------------------------------------
def load_neuropil_max_slope_threshold_from_col_d(main_folder: Path) -> float:
    """
    Reads the max slope threshold from:
    main_folder/Neuropil_Filter/Neuropil_Filter_max_slope_metrics_Normalised.xlsx
    Sheet: 'Statistics'
    Column: D (first numeric value found in that column, skipping the header row)
    """
    excel_path = (
        main_folder
        / "Neuropil_Filter"
        / "Neuropil_Filter_max_slope_metrics_Normalised.xlsx"
    )

    if not excel_path.exists():
        raise FileNotFoundError(f"Could not find neuropil threshold file: {excel_path}")

    wb = openpyxl.load_workbook(excel_path, data_only=True)

    if "Statistics" not in wb.sheetnames:
        raise KeyError(f"Sheet 'Statistics' not found in {excel_path}")

    ws = wb["Statistics"]

    for row in range(2, ws.max_row + 1):
        val = ws[f"D{row}"].value
        if val is None:
            continue
        try:
            return float(val)
        except (TypeError, ValueError):
            continue

    raise ValueError(f"No numeric threshold found in column D of 'Statistics' in {excel_path}")


NEUROPIL_MAX_SLOPE_THRESHOLD = load_neuropil_max_slope_threshold_from_col_d(main_folder)
print(f"Using Neuropil max slope threshold (from column D): {NEUROPIL_MAX_SLOPE_THRESHOLD}")


# -------------------------------------------------------------------
# PREPROCESSING
# -------------------------------------------------------------------
def apply_savitzky_golay(df, window_length=Fil_Win, polyorder=3):
    """Apply Savitzky-Golay filter to smooth the data."""
    return df.apply(
        lambda x: savgol_filter(x, window_length=window_length, polyorder=polyorder),
        axis=0,
    )


def normalize_data(df):
    """Normalize data to a range of 0 to 1."""
    return (df - df.min()) / (df.max() - df.min())


# -------------------------------------------------------------------
# METRICS + CLASSIFICATION
# -------------------------------------------------------------------
def calculate_metrics(
    polygon_df,
    window_size=Met_win,
    max_slope_threshold=NEUROPIL_MAX_SLOPE_THRESHOLD,
):
    """Calculate metrics for each ROI in a polygon DataFrame, including classification."""
    metrics_list = []
    accepted_rois = []

    for roi in polygon_df.columns:
        data = polygon_df[roi]
        max_slope = -float("inf")
        max_slope_start_frame = None

        for start in range(0, len(data) - window_size):
            subset = data.iloc[start : start + window_size]
            slope = (subset.iloc[-1] - subset.iloc[0]) / (len(subset) - 1) * FR

            if slope > max_slope:
                max_slope = slope
                max_slope_start_frame = start

        mean_first_50 = data.iloc[:Mean_win].mean()
        mean_last_50 = data.iloc[-Mean_win:].mean()

        classification = (
            "Reject"
            if (max_slope <= max_slope_threshold) or (mean_first_50 > mean_last_50 + 0.5)
            else "Accept"
        )

        if classification == "Accept":
            accepted_rois.append(roi)

        metrics_list.append(
            pd.DataFrame(
                {
                    "ROI": [roi],
                    "Slope_Max": [max_slope],
                    "Max_Slope_Start_Frame": [max_slope_start_frame],
                    "Mean_First_50": [mean_first_50],
                    "Mean_Last_50": [mean_last_50],
                    "Classification": [classification],
                    "Max_Slope_Threshold_Used": [max_slope_threshold],
                }
            )
        )

    return pd.concat(metrics_list).reset_index(drop=True), accepted_rois


# -------------------------------------------------------------------
# PROCESS ONE POLYGON DF
# -------------------------------------------------------------------
def process_polygon_data(polygon_df, polygon_name, base_folder, window_size=Met_win):
    if polygon_df.empty:
        print(f"No valid data for {polygon_name}.")
        return pd.DataFrame(), pd.DataFrame()

    print(f"Processing {polygon_name}")

    data_smoothed = apply_savitzky_golay(polygon_df)
    data_normalized = normalize_data(data_smoothed)

    metrics_df, accepted_rois = calculate_metrics(data_normalized, window_size=window_size)

    folder_name = os.path.basename(os.path.normpath(base_folder))
    output_path = os.path.join(
        base_folder, f"{folder_name}_{polygon_name}_Accepted_Rejected_Plots.pdf"
    )

    with PdfPages(output_path) as pdf:
        for _, row in metrics_df.iterrows():
            roi = row["ROI"]
            classification = row["Classification"]
            line_color = "green" if classification == "Accept" else "red"

            fig, axes = plt.subplots(nrows=1, ncols=2, figsize=(12, 4), tight_layout=True)

            # Left: raw trace
            sns.lineplot(data=polygon_df[roi], ax=axes[0], label=roi, color=line_color)
            axes[0].set_title(f"{roi} - Raw Data ({classification})", fontsize=12)
            axes[0].set_xlabel("Frame")
            axes[0].set_ylabel("Intensity")
            axes[0].grid(False)

            # Right: smoothed & normalized trace
            sns.lineplot(data=data_normalized[roi], ax=axes[1], label=roi, color=line_color)
            axes[1].set_title(
                f'{roi} - Smoothed & Normalized ({classification})\n'
                f'Max Slope: {row["Slope_Max"]:.4f}  Threshold: {row["Max_Slope_Threshold_Used"]:.4f}',
                fontsize=12,
            )
            axes[1].set_xlabel("Frame")
            axes[1].set_ylabel("Normalized Intensity")
            axes[1].grid(False)

            if row["Max_Slope_Start_Frame"] is not None:
                start = int(row["Max_Slope_Start_Frame"])
                subset = data_normalized[roi].iloc[start : start + window_size]
                axes[1].plot(
                    range(start, start + window_size),
                    subset,
                    color="orange",
                    label="Max Slope Window",
                )
                axes[1].legend()

            pdf.savefig(fig)
            plt.close(fig)

    accepted_df = polygon_df[accepted_rois].copy()
    accepted_df.insert(0, "Frames", range(1, len(accepted_df) + 1))

    return metrics_df, accepted_df


# -------------------------------------------------------------------
# RUN ALL POLYGONS
# -------------------------------------------------------------------
metrics_results = {}
accepted_dfs = {}

for df_name in all_created_dfs:
    polygon_df = globals()[df_name]
    polygon_name = df_name.replace("_df", "")
    metrics_df, accepted_df = process_polygon_data(polygon_df, polygon_name, base_folder)

    metrics_results[polygon_name] = metrics_df
    accepted_dfs[polygon_name] = accepted_df


# -------------------------------------------------------------------
# SUMMARY BAR PLOT
# -------------------------------------------------------------------
def plot_trace_classification_summary(metrics_results, base_folder):
    """Generate a bar plot showing the number of accepted and rejected traces per region."""
    summary_data = []
    folder_name = os.path.basename(os.path.normpath(base_folder))

    for region, metrics_df in metrics_results.items():
        accepted_count = (metrics_df["Classification"] == "Accept").sum()
        rejected_count = (metrics_df["Classification"] == "Reject").sum()
        summary_data.append(
            {"Region": region, "Accepted": accepted_count, "Rejected": rejected_count}
        )

    summary_df = pd.DataFrame(summary_data)
    summary_df.set_index("Region", inplace=True)

    summary_df.plot(
        kind="bar",
        stacked=True,
        figsize=(10, 6),
        color=["green", "red"],
    )
    plt.title("Accepted vs Rejected Traces per Region")
    plt.xlabel("Region")
    plt.ylabel("Number of Traces")
    plt.legend(title="Classification")
    plt.xticks(rotation=45)
    plt.tight_layout()
    plt.savefig(os.path.join(base_folder, f"{folder_name}_Accepted_Rejected_Traces_BarPlot.pdf"))
    plt.close()


plot_trace_classification_summary(metrics_results, base_folder)


# -------------------------------------------------------------------
# SAVE ACCEPTED TRACES PER POLYGON
# -------------------------------------------------------------------
for polygon_name, accepted_df in accepted_dfs.items():
    folder_name = os.path.basename(os.path.normpath(base_folder))
    accepted_output_path = os.path.join(
        base_folder, f"{folder_name}_{polygon_name}_Accepted_Traces.xlsx"
    )
    accepted_df.to_excel(accepted_output_path, index=False)
    print(f"Accepted traces for {polygon_name} saved to {accepted_output_path}")
```
