## Supplemental file 3 for "Protocol for calcium imaging of acute brain slices from *Octopus vulgaris* hatchlings during application of neurotransmitters": Octo_CaImaging_Part2_Categorisation.ipynb

```
# %% [code]
import os
import sys
import json
from pathlib import Path

import copy

from PIL import Image

import matplotlib.pyplot as plt
from matplotlib.colors import ListedColormap, BoundaryNorm
from matplotlib.backends.backend_pdf import PdfPages
from matplotlib.patches import Rectangle

import numpy as np
import pandas as pd

from sklearn.linear_model import LinearRegression
from sklearn.preprocessing import MinMaxScaler

from scipy.ndimage import gaussian_filter1d, uniform_filter1d

import seaborn as sns
import plotly.graph_objects as go

from ipyfilechooser import FileChooser
from IPython.display import display
import ipywidgets as widgets
DEFAULT_MAIN = r"C:/Users\Amy Courtney\Documents/Temp/Calcium_Imaging_Paper/Biorxiv_Submission1_Mar26/Supp_Files/Courtney_et_al_2026_Supp_File_3_Sample_Dataset/Calcium_Imaging_Analysis"
DEFAULT_BASE = r"C:/Users\Amy Courtney\Documents/Temp/Calcium_Imaging_Paper/Biorxiv_Submission1_Mar26/Supp_Files/Courtney_et_al_2026_Supp_File_3_Sample_Dataset/Calcium_Imaging_Analysis"

main_fc = FileChooser(DEFAULT_MAIN, select_desc="Main Folder")
main_fc.show_only_dirs = True

base_fc = FileChooser(DEFAULT_BASE, select_desc="Single Video Folder")
base_fc.show_only_dirs = True

display(main_fc)
display(base_fc)

# %% [code]
from pathlib import Path

main_folder = Path(main_fc.selected_path).resolve()
base_folder = Path(base_fc.selected_path).resolve()

print("Main folder:", main_folder)
print("Base folder:", base_folder)


# %% [code]
desired_order = ['OGL', 'IGL', 'Med', 'CB']

# %% [code]
#categorisation step 1

import os
import pandas as pd

# Define helper functions
def apply_rolling_average(df, window_size=20):
    return df.rolling(window=window_size, min_periods=1).mean()

def normalize_data(df):
    return (df - df.min()) / (df.max() - df.min())

def categorize_cell(cell_data):
    baseline = cell_data[cell_data['Timepoint'] == 'Baseline']['Mean'].values
    ligand = cell_data[cell_data['Timepoint'] == 'Ligand']['Mean'].values
    highk = cell_data[cell_data['Timepoint'] == 'HighK+']['Mean'].values

    if len(baseline) > 0 and len(ligand) > 0:
        baseline_val = baseline[0]
        ligand_val = ligand[0]

        if ligand_val <= baseline_val - 0.2:
            return 'Inhibition'
        elif ligand_val > baseline_val + 0.2:
            return 'Excitation'
    return 'Non-Responding'

# ---------------------------------------------------------
# Fixed time windows for ALL videos (no Excel lookup)
Base_Start = 10
Base_End   = 100
Lig_Start  = 500
Lig_End    = 600
K_Start    = 1500
K_End      = 1600

print(
    f"Using fixed time windows for all videos: "
    f"Base_Start={Base_Start}, Base_End={Base_End}, "
    f"Lig_Start={Lig_Start}, Lig_End={Lig_End}, "
    f"K_Start={K_Start}, K_End={K_End}."
)
# ---------------------------------------------------------

# Process all accepted traces
files_to_process = [
    os.path.join(base_folder, file_name)
    for file_name in os.listdir(base_folder)
    if file_name.endswith("_Accepted_Traces.xlsx")
]

if not files_to_process:
    print(f"No files ending with '_Accepted_Traces.xlsx' found in {base_folder}.")
    exit()

categorization_data_per_file = {}
normalized_data_dict = {}

for file_path in files_to_process:
    file_name = os.path.basename(file_path).replace('_Accepted_Traces.xlsx', '')

    try:
        data = pd.read_excel(file_path)
        print(f"Successfully loaded data from {file_path}.")
    except Exception as e:
        print(f"Error loading file: {e}")
        continue

    if data.empty:
        print(f"The file {file_path} is empty. Skipping.")
        continue

    if 'Frames' in data.columns:
        data = data.drop(columns=['Frames'])

    # Apply rolling average and normalization
    filtered_data = apply_rolling_average(data)
    normalized_data = normalize_data(filtered_data)

    # Store the normalized data
    normalized_data_dict[file_name] = normalized_data

    # Ensure mean_values_df is initialized correctly
    all_mean_values = []

    for file_name, normalized_data in normalized_data_dict.items():
        mean_values = []
        
        for roi in normalized_data.columns:
            baseline_mean = normalized_data[roi].iloc[Base_Start:Base_End].mean() if not normalized_data[roi].iloc[Base_Start:Base_End].isna().all() else None
            ligand_mean = normalized_data[roi].iloc[Lig_Start:Lig_End].mean() if not normalized_data[roi].iloc[Lig_Start:Lig_End].isna().all() else None
            highk_mean = normalized_data[roi].iloc[K_Start:K_End].mean() if not normalized_data[roi].iloc[K_Start:K_End].isna().all() else None

            mean_values.append({
                'ROI': roi,
                'Baseline_Mean': baseline_mean,
                'Ligand_Mean': ligand_mean,
                'HighK_Mean': highk_mean
            })

        # Append all ROIs to the global list
        all_mean_values.extend(mean_values)

    # Convert to DataFrame
    mean_values_df = pd.DataFrame(all_mean_values)

    # Ensure missing values are handled
    mean_values_df.fillna(0, inplace=True)

    categorization_data = []
    for roi in normalized_data.columns:
        means = {
            'Baseline': mean_values_df.loc[mean_values_df['ROI'] == roi, 'Baseline_Mean'].values[0],
            'Ligand': mean_values_df.loc[mean_values_df['ROI'] == roi, 'Ligand_Mean'].values[0],
            'HighK+': mean_values_df.loc[mean_values_df['ROI'] == roi, 'HighK_Mean'].values[0]
        }
        category = categorize_cell(pd.DataFrame({
            'Timepoint': list(means.keys()),
            'Mean': list(means.values())
        }))
        categorization_data.append({'ID': roi, 'Category': category})

    categorization_data_per_file[file_name] = pd.DataFrame(categorization_data)
    print(f"Categorization data stored for {file_name}.")

# Save categorized data and mean values
%store categorization_data_per_file
%store normalized_data_dict
%store mean_values_df


# %% [code]
#cat step 2 - refine inhib - in frames

lowest_folder_name = os.path.basename(os.path.normpath(base_folder))
FR = 5
Win_St = 200
Win_End = 200
SMOOTH_WINDOW = 50
WINDOW_SIZE = 150
MAX_FRAME = 300
INHIBITION_THRESHOLD = -0.0199 # Assuming threshold should also be reduced (adjust if necessary)
MEAN_WINDOW = 100
MEAN_THRESHOLD = 0.2 # Assuming this threshold should also be reduced (adjust if necessary)

# Function to smooth the trace
def smooth_trace(trace, window=SMOOTH_WINDOW):
    return uniform_filter1d(trace, size=window, mode='nearest')

def detect_steepest_slope(trace):
    smoothed_trace = smooth_trace(trace)
    max_search_index = min(MAX_FRAME, len(smoothed_trace))
    steepest_slope = None
    steepest_window = (None, None)

    for i in range(max_search_index - WINDOW_SIZE):
        window_x = np.arange(i, i + WINDOW_SIZE).reshape(-1, 1)
        window_y = smoothed_trace[i:i + WINDOW_SIZE]
        
        model = LinearRegression().fit(window_x, window_y)
        slope = model.coef_[0]  # Original slope (fluorescence per frame)

        # Convert to fluorescence change per second
        slope_per_second = slope * FR  

        if steepest_slope is None or slope_per_second < steepest_slope:
            steepest_slope = slope_per_second  # Store converted slope
            steepest_window = (i, i + WINDOW_SIZE)

    return steepest_slope, steepest_window


# Function to compute mean fluorescence after the steepest slope
def compute_mean_after_slope(trace, slope_end):
    smoothed_trace = smooth_trace(trace)

    if slope_end is None or slope_end + MEAN_WINDOW > len(smoothed_trace):
        return None  # Avoid out-of-range errors

    mean_value = np.mean(smoothed_trace[slope_end:slope_end + MEAN_WINDOW])
    return mean_value

# Function to classify traces based on the steepest slope and mean value
def classify_traces(matching_data):
    trace_data = []
    for file_key, df in matching_data.items():
        for col in df.columns:
            trace = smooth_trace(df[col])  
            steepest_slope, (slope_start, slope_end) = detect_steepest_slope(trace)

            # Compute mean fluorescence after steepest slope
            mean_value = compute_mean_after_slope(trace, slope_end)

            # Ensure valid mean values
            if mean_value is None:
                classification = "Non-Responding"
            else:
                # If slope is negative enough AND mean is below threshold, classify as inhibition
                if steepest_slope < INHIBITION_THRESHOLD and mean_value < MEAN_THRESHOLD:
                    classification = "Inhibition"
                else:
                    classification = "Non-Responding"

            trace_data.append({
                'File': file_key, 
                'ID': col, 
                'Steepest_Slope': steepest_slope,
                'Slope_Start': slope_start,
                'Slope_End': slope_end,
                'Mean_After_Slope': mean_value,
                'Classification': classification
            })
    
    df_traces = pd.DataFrame(trace_data)
    df_traces.sort_values(by='Steepest_Slope', inplace=True)
    return df_traces

# Function to plot traces sorted by steepest slope
def plot_traces(matching_data, classified_traces_df):
    sorted_ids = classified_traces_df['ID'].tolist()
    
    num_traces = len(sorted_ids)
    num_rows = (num_traces // 5) + (1 if num_traces % 5 else 0)
    fig, axes = plt.subplots(num_rows, 5, figsize=(15, 3 * num_rows))
    axes = axes.flatten()
    
    plotted_ids = set()
    for i, col in enumerate(sorted_ids):
        if col in plotted_ids:
            continue
        for file_key, df in matching_data.items():
            if col in df.columns:
                smoothed_trace = smooth_trace(df[col])
                trend_row = classified_traces_df[classified_traces_df['ID'] == col].iloc[0]
                slope_start, slope_end = trend_row['Slope_Start'], trend_row['Slope_End']
                steepest_slope = trend_row['Steepest_Slope']
                classification = trend_row['Classification']
                mean_after_slope = trend_row['Mean_After_Slope']
                
                # Set color based on classification
                color = "red" if classification == "Inhibition" else "black"
                
                axes[i].plot(df.index, smoothed_trace, label='Smoothed', color=color)
                
                # Highlight the steepest slope window
                if slope_start is not None and slope_end is not None:
                    axes[i].axvline(df.index[slope_start], color='red', linestyle='--', label='Slope Start')
                    axes[i].axvline(df.index[slope_end], color='orange', linestyle='--', label='Slope End')
                
                    # Highlight the 200-frame window after slope_end in green
                    if slope_end + MEAN_WINDOW < len(df.index):  # Ensure within range
                        axes[i].axvspan(df.index[slope_end], df.index[slope_end + MEAN_WINDOW], 
                                        color='green', alpha=0.3, label='Mean Window')

                        # Display the mean value in a green box
                        axes[i].text(df.index[slope_end], np.mean(smoothed_trace), 
                                     f"Mean: {mean_after_slope:.3f}", 
                                     bbox=dict(facecolor='green', alpha=0.5), fontsize=10, color='black')

                axes[i].set_title(f"{col} (Sl: {steepest_slope:.5f})")
                axes[i].set_xlabel('Time (frames)')
                axes[i].set_ylabel('Normalized Intensity')
                
                plotted_ids.add(col)
                break  # Ensure the ID is plotted only once
    
    plt.suptitle("Fluorescence Traces Sorted by Steepest Slope")
    plt.tight_layout(rect=[0, 0.03, 1, 0.95])
    plt.show()
def expand_columns_to_dfs(categorization_data_per_file):
    dfs = {}
    for col_name, df in categorization_data_per_file.items():
        dfs[col_name] = df.copy()
    return dfs

def extract_inhibition_ids(dfs):
    inhibition_ids = []
    for name, df in dfs.items():
        if 'Category' in df.columns and 'ID' in df.columns:
            filtered_ids = df[df['Category'].str.contains('Inhibition', na=False)]['ID'].tolist()
            inhibition_ids.extend(filtered_ids)
    return pd.DataFrame({'ID': inhibition_ids})

def extract_matching_data(normalized_data_dict, inhibition_df):
    extracted_data = {}
    for file_key, df in normalized_data_dict.items():
        matching_columns = [col for col in df.columns if col in inhibition_df['ID'].values]
        if matching_columns:
            df_extracted = df.loc[Lig_Start - Win_St:Lig_End + Win_End, matching_columns]
            df_normalized = df_extracted.apply(lambda x: (x - x.min()) / (x.max() - x.min()), axis=0)
            extracted_data[file_key] = df_normalized
        else:
            print(f"Warning: No matching IDs found in {file_key}")
    return extracted_data
# Run the function
expanded_dfs = expand_columns_to_dfs(categorization_data_per_file)
Inhibition_IDs = extract_inhibition_ids(expanded_dfs)


# Check if there are any inhibition IDs before proceeding
if Inhibition_IDs.empty:
    print("No inhibition IDs found. Skipping classification and plotting.")
else:
    matching_data = extract_matching_data(normalized_data_dict, Inhibition_IDs)
    
    if not matching_data:  # Ensure that matching_data is not empty
        print("No matching inhibition data found. Skipping classification and plotting.")
    else:
        classified_traces_df = classify_traces(matching_data)
        plot_traces(matching_data, classified_traces_df)
        
        def update_categories(categorization_data_per_file, min_slope_df):
            """
            Updates the Category values in a copy of categorization_data_per_file based on matching ID values in min_slope_df.
            
            Parameters:
            categorization_data_per_file (dict): Dictionary containing DataFrames per file.
            min_slope_df (pd.DataFrame): DataFrame with columns 'ID' and 'New_Cat'.
            
            Returns:
            dict: Updated copy of categorization_data_per_file with updated categories.
            """
            # Convert min_slope_df to a dictionary for faster lookup
            id_to_new_cat = dict(zip(min_slope_df['ID'], min_slope_df['Classification']))
            
            # Debugging: Print the mapping dictionary
            print("ID to New Category Mapping:", id_to_new_cat)
            
            # Create a deep copy to avoid modifying the original dictionary
            updated_data = copy.deepcopy(categorization_data_per_file)
            
            # Iterate through the dictionary and update Category column in DataFrame
            for file_name, df in updated_data.items():
                print(f"\nProcessing file: {file_name}")
                
                if isinstance(df, pd.DataFrame):
                    print("Before Update:")
                    print(df.head())

                    # Update the 'Category' column based on matching IDs
                    df['Category'] = df['ID'].map(id_to_new_cat).fillna(df['Category'])

                    print("After Update:")
                    print(df.head())
                else:
                    print(f"Warning: {file_name} does not contain a DataFrame!")

            return updated_data


        # Example input
        updated_data = update_categories(categorization_data_per_file, classified_traces_df)

        print("\nUpdated Data:")
        for file_name, df in updated_data.items():
            print(f"\n{file_name}:")
            print(df)

        %store -r updated_data


        def expand_columns_to_dfs(categorization_data_per_file):
            dfs = {}
            for col_name, df in categorization_data_per_file.items():
                dfs[col_name] = df.copy()
            return dfs

        def extract_inhibition_ids(dfs):
            inhibition_ids = []
            for name, df in dfs.items():
                if 'Category' in df.columns and 'ID' in df.columns:
                    filtered_ids = df[df['Category'].str.contains('Inhibition', na=False)]['ID'].tolist()
                    inhibition_ids.extend(filtered_ids)
            return pd.DataFrame({'ID': inhibition_ids})

        def extract_matching_data(normalized_data_dict, inhibition_df):
            extracted_data = {}
            for file_key, df in normalized_data_dict.items():
                matching_columns = [col for col in df.columns if col in inhibition_df['ID'].values]
                if matching_columns:
                    df_extracted = df.loc[Lig_Start - 200:Lig_End + 300, matching_columns]
                    df_normalized = df_extracted.apply(lambda x: (x - x.min()) / (x.max() - x.min()), axis=0)
                    extracted_data[file_key] = df_normalized
                else:
                    print(f"Warning: No matching IDs found in {file_key}")
            return extracted_data


        def plot_traces_only(matching_data):
            for file_key, df in matching_data.items():
                num_traces = df.shape[1]
                num_rows = (num_traces // 5) + (1 if num_traces % 5 else 0)
                fig, axes = plt.subplots(num_rows, 5, figsize=(15, 3 * num_rows))
                axes = axes.flatten()
                
                for i, col in enumerate(df.columns):
                    axes[i].plot(df.index, df[col], label=col)
                    axes[i].set_title(f"{col}")
                    axes[i].set_xlabel('Time')
                    axes[i].set_ylabel('Normalized Intensity')
                    axes[i].legend()
                
                for j in range(i + 1, len(axes)):
                    fig.delaxes(axes[j])
                
                plt.suptitle(f"Fluorescence Traces for {file_key}")
                plt.tight_layout(rect=[0, 0.03, 1, 0.95])
                plt.show()

        # Run the function
        # Run the function
        expanded_dfs = expand_columns_to_dfs(updated_data)
        Inhibition_IDs = extract_inhibition_ids(expanded_dfs)
        matching_data = extract_matching_data(normalized_data_dict, Inhibition_IDs)
        plot_traces_only(matching_data)


# %% [code]
def plot_traces_overlay(matching_data, base_folder):
    lowest_folder_name = os.path.basename(os.path.normpath(base_folder))
    
    for file_key, df in matching_data.items():
        plt.figure(figsize=(6, 6))

        # Plot all traces in grey
        for col in df.columns:
            plt.plot(df.index, df[col], color='grey', linewidth=0.1)

        # Compute and plot the mean trace in #AF8DC3
        mean_trace = df.mean(axis=1)
        plt.plot(df.index, mean_trace, color='#AF8DC3', linewidth=1, label='Mean Trace')

        plt.title(f"Fluorescence Traces for {file_key}")
        plt.xlabel('Time')
        plt.ylabel('Normalized Intensity')
        
        # Generate unique filenames per file_key
        plot_file_name_pdf = f"{file_key}_Example_Inhib_Traces.pdf"
        plot_file_name_tiff = f"{file_key}_Example_Inhib_Traces.tiff"
        plot_file_path_pdf = os.path.join(base_folder, plot_file_name_pdf)
        plot_file_path_tiff = os.path.join(base_folder, plot_file_name_tiff)

        # Save the plot as PDF and TIFF
        plt.savefig(plot_file_path_pdf, format='pdf', bbox_inches='tight')
        plt.savefig(plot_file_path_tiff, format='tiff', bbox_inches='tight')
        
        plt.close()  # Close the figure to free memory

# Call the function with matching_data and base_folder
plot_traces_overlay(matching_data, base_folder)


# %% [code]
def plot_traces_overlay(matching_data, base_folder):
    lowest_folder_name = os.path.basename(os.path.normpath(base_folder))
    
    plt.figure(figsize=(8, 6))  # Create a single figure
    
    all_traces = []
    
    # Iterate through all files and overlay traces on one plot
    for file_key, df in matching_data.items():
        for col in df.columns:
            plt.plot(df.index, df[col], color='grey', linewidth=0.1)
        
        all_traces.append(df)
    
    # Compute and plot the overall mean trace in #AF8DC3
    combined_df = np.concatenate([df.values for df in all_traces], axis=1)
    overall_mean_trace = np.mean(combined_df, axis=1)
    plt.plot(df.index, overall_mean_trace, color='#AF8DC3', linewidth=2, label='Overall Mean Trace')
    
    plt.xlabel("Time")
    plt.ylabel("Normalized Intensity")
    
    # Generate filenames for the combined plot
    plot_file_name_pdf = "Combined_Example_Inhib_Traces.pdf"
    plot_file_name_tiff = "Combined_Example_Inhib_Traces.tiff"
    plot_file_path_pdf = os.path.join(base_folder, plot_file_name_pdf)
    plot_file_path_tiff = os.path.join(base_folder, plot_file_name_tiff)
    
    # Save the plot
    plt.savefig(plot_file_path_pdf, format='pdf', bbox_inches='tight')
    plt.savefig(plot_file_path_tiff, format='tiff', bbox_inches='tight')
    
    plt.show()  # Show the final combined plot


plot_traces_overlay(matching_data, base_folder)


# %% [code]


# Ensure the DataFrame contains 'Steepest_Slope' column
if 'Steepest_Slope' not in classified_traces_df.columns:
    raise ValueError("The DataFrame must contain a 'Steepest_Slope' column.")

# Create the figure
fig, ax = plt.subplots(figsize=(8, 6))

# Define aesthetics
palette = "muted"
alpha_violin = 0.5  # Transparency for violin plot
alpha_strip = 0.7  # Transparency for individual data points

# Violin plot (distribution) - Background layer
sns.violinplot(
    y=classified_traces_df['Steepest_Slope'],
    ax=ax,
    inner=None,
    palette=palette,
    alpha=alpha_violin  # Transparency for visibility
)

# Strip plot (individual data points) - Middle layer
sns.stripplot(
    y=classified_traces_df['Steepest_Slope'],
    ax=ax,
    jitter=True,
    size=8,
    alpha=alpha_strip,
    color="black"
)

# Point plot (mean values) - Top layer
sns.pointplot(
    y=classified_traces_df['Steepest_Slope'],
    ax=ax,
    color="grey",  # Mean values in grey
    markers="D",  # Diamond marker for mean
    scale=1.2,  # Slightly larger marker
    errwidth=1.5,  # Error bar width
    capsize=0.2,  # Small caps on error bars
    join=False,
    estimator=np.mean,
    zorder=3  # Ensure point plot is on top
)

# Add a dotted black line at slope -0.0039
ax.axhline(y=-0.0199, color='black', linestyle='dotted', linewidth=1.5)

# Labels and title
ax.set_title('Enhanced Violin Plot of Steepest Slope')
ax.set_ylabel('Steepest Slope')

# Set y-axis limits
#ax.set_ylim(-0.009, -0.00001)

# Show the plot
plt.show()


# %% [code]
# Retrieve stored variables
%store -r updated_data
%store -r normalized_data_dict
%store -r mean_values_df
%store -r Base_Start
%store -r Base_End
%store -r Lig_Start
%store -r Lig_End
%store -r K_Start
%store -r K_End

for file_name, normalized_data in normalized_data_dict.items():
    output_pdf_path = os.path.join(base_folder, f"{file_name}_TimepointPlots.pdf")

    
    # Check if there are any inhibition IDs before proceeding
    if Inhibition_IDs.empty:
        print("No inhibition IDs found. Keeping categorization_data_per_file")
        categorization_df = categorization_data_per_file.get(file_name, None)
    else:
        print("Inhibition IDs found. Using updated_data")
        # Retrieve categorization data for this file
        categorization_df = updated_data.get(file_name, None)

    if categorization_df is None:
        print(f"Warning: No categorization data found for {file_name}. Skipping.")
        continue

    with PdfPages(output_pdf_path) as pdf:
        num_rois = len(normalized_data.columns)
        rois_per_page = 4
        num_pages = (num_rois + rois_per_page - 1) // rois_per_page

        for page in range(num_pages):
            start_idx = page * rois_per_page
            end_idx = min((page + 1) * rois_per_page, num_rois)
            rois_to_plot = normalized_data.columns[start_idx:end_idx]

            fig, axes = plt.subplots(len(rois_to_plot), 2, figsize=(12, 4 * len(rois_to_plot)), constrained_layout=True)
            if len(rois_to_plot) == 1:
                axes = [axes]

            for idx, roi in enumerate(rois_to_plot):
                axes[idx][0].plot(normalized_data[roi], color='blue')
                axes[idx][0].set_title(f'{roi} - Normalized Intensity Trace')
                axes[idx][0].set_xlabel('Frame')
                axes[idx][0].set_ylabel('Normalized Intensity')
                axes[idx][0].grid(True)

                # Add time window markers as dotted black lines
                time_windows = [Base_Start, Base_End, Lig_Start, Lig_End, K_Start, K_End]
                for tw in time_windows:
                    axes[idx][0].axvline(tw, color='blue', linestyle='dotted', linewidth=1.5)

                # Timepoint mean plot (right)
                timepoints = ['Baseline', 'Ligand', 'HighK+']
                roi_data = mean_values_df[mean_values_df['ROI'] == roi]
                
                if roi_data.empty:
                    print(f"Warning: ROI {roi} not found in mean_values_df. Skipping.")
                    continue

                means = [
                    roi_data['Baseline_Mean'].values[0],
                    roi_data['Ligand_Mean'].values[0],
                    roi_data['HighK_Mean'].values[0]
                ]

                # Get category from categorization data
                category_row = categorization_df[categorization_df['ID'] == roi]
                category = category_row['Category'].values[0] if not category_row.empty else 'Unknown'

                # Set color based on category
                line_color = {'Non-Responding': 'black', 'Excitation': 'green', 'Inhibition': 'red'}.get(category, 'gray')

                axes[idx][1].plot(timepoints, means, marker='o', color=line_color, linestyle='-', label='Mean Values')
                axes[idx][1].set_ylim(0, 1)
                axes[idx][1].set_title(f'{roi} - Timepoint Means ({category})')
                axes[idx][1].grid(True)
                axes[idx][1].legend()

            pdf.savefig(fig)
            plt.close(fig)

    print(f"Timepoint plots saved in {output_pdf_path}.")


# %% [code]
# Check if there are any inhibition IDs before proceeding
if Inhibition_IDs.empty:
    print("No inhibition IDs found. Keeping categorization_data_per_file")
    # Save each categorization DataFrame as an Excel file
    for file_name, categorization_df in categorization_data_per_file.items():
        excel_path = os.path.join(base_folder, f"{file_name}_Categorization.xlsx")
        
        try:
            categorization_df.to_excel(excel_path, index=False)
            print(f"Categorization data saved: {excel_path}")
        except Exception as e:
            print(f"Error saving {file_name} categorization data: {e}")

else:
    print("Inhibition IDs found. Using updated_data")
    # Save each categorization DataFrame as an Excel file
    for file_name, categorization_df in updated_data.items():
        excel_path = os.path.join(base_folder, f"{file_name}_Categorization.xlsx")
        
        try:
            categorization_df.to_excel(excel_path, index=False)
            print(f"Categorization data saved: {excel_path}")
        except Exception as e:
            print(f"Error saving {file_name} categorization data: {e}")


# %% [code]
# Locate the folder name from base_folder
folder_name = os.path.basename(os.path.normpath(base_folder))
end = "_Categorization.xlsx"
start_index = len(folder_name) + 1  # Start right after base_folder and the underscore
end_index = -len(end)  # Go up to the start of the end string


# Automatically find all files ending with '_Categorization.xlsx'
categorization_files = [
    os.path.join(base_folder, file_name)
    for file_name in os.listdir(base_folder)
    if file_name.endswith('_Categorization.xlsx')
]

# Categories to calculate percentages for
categories = ['Excitation', 'Inhibition', 'Non-Responding']

# Initialize a list to store results
results = []

# Loop through each file
for file_path in categorization_files:
    try:
        # Load categorization data
        categorization_data = pd.read_excel(file_path)

        # Extract the polygon name from the file name
        polygon_name = os.path.basename(file_path)[start_index:end_index]
        
        # Calculate total ROIs
        total_rois = len(categorization_data)

        # Calculate percentages
        percentages = {'Polygon': polygon_name, 'n': total_rois}  # Add total ROIs to the results
        for category in categories:
            category_count = categorization_data[categorization_data['Category'] == category].shape[0]
            percentages[category] = (category_count / total_rois) * 100

        results.append(percentages)

    except Exception as e:
        print(f"Error processing file {file_path}: {e}")

# Convert results to a DataFrame
results_df = pd.DataFrame(results)

# Save results to an Excel file in the base folder
output_file = os.path.join(base_folder, f"{os.path.basename(base_folder)}_Category_Percentages.xlsx")
results_df.to_excel(output_file, index=False)
print(f"Category percentages saved to {output_file}")


# %% [code]
def find_file_with_suffix(base_folder, suffix="Category_Percentages.xlsx"):
    """
    Searches for a file ending with the specified suffix in the given base folder.

    Args:
        base_folder (str): The directory to search in.
        suffix (str): The suffix to look for. Defaults to "Category_Percentages.xlsx".

    Returns:
        str: The full path to the file if found, else None.
    """
    for root, dirs, files in os.walk(base_folder):
        for file in files:
            if file.endswith(suffix):
                return os.path.join(root, file)
    return None

file_path = find_file_with_suffix(base_folder)

if file_path:
    print(f"File found: {file_path}")
else:
    print("No file ending with 'Category_Percentages.xlsx' found.")

# Load the data
category_data = pd.read_excel(file_path)

# Filter and reorder the DataFrame based on the desired order
category_data['Polygon'] = pd.Categorical(category_data['Polygon'], categories=desired_order, ordered=True)
category_data = category_data.sort_values(by='Polygon').reset_index(drop=True)

# Extract the reordered regions, category percentages, and total cell counts
regions = category_data['Polygon']
non_responding = category_data['Non-Responding']
excitation = category_data['Excitation']
inhibition = category_data['Inhibition']
total_cells = category_data['n']

# Plotting
x = np.arange(len(regions))  # the label locations
width = 0.25  # the width of the bars

fig, ax = plt.subplots(figsize=(10, 6))
# Adjust the order of bars: Excitation, Inhibition, Non-Responding
bars1 = ax.bar(x - width, excitation, width, edgecolor='black', label='Excitation', color='#7fbf7b')
bars2 = ax.bar(x, inhibition, width, edgecolor='black', label='Inhibition', color='#af8dc3')
bars3 = ax.bar(x + width, non_responding, width, edgecolor='black', label='Non-Responding', color='#7F7F7F')

# Add labels, title, and custom x-axis tick labels
ax.set_ylabel('Percent Cells', fontsize=12)
ax.set_title('Percentage of Cells by Category in Each Polygon', fontsize=14)
ax.set_xticks(x)
ax.set_xticklabels(regions, fontsize=10)

# Move the legend outside the plot
ax.legend(loc='upper left', bbox_to_anchor=(1, 1), fontsize=10)

# Add n= label under each region
for i in range(len(regions)):
    ax.text(x[i], -10, f'n={int(total_cells[i])}', ha='center', fontsize=10)  # Adjusted to -10 for lower positioning

# Save the plot as a PDF and TIFF
lowest_folder_name = os.path.basename(os.path.normpath(base_folder))
plot_file_name_pdf = f"{lowest_folder_name}_Category_Barplot.pdf"
plot_file_name_tiff = f"{lowest_folder_name}_Category_Barplot.tiff"
plot_file_path_pdf = os.path.join(base_folder, plot_file_name_pdf)
plot_file_path_tiff = os.path.join(base_folder, plot_file_name_tiff)

# Save the plot as PDF
plt.savefig(plot_file_path_pdf, format='pdf', bbox_inches='tight')  # Ensure the legend is included in the saved plot
# Save the plot as TIFF
plt.savefig(plot_file_path_tiff, format='tiff', bbox_inches='tight')

# Display the plot
plt.show()


# %% [code]
#generate heatmaps per region - min/max normalisation

# Load fluorescence data
def load_fluorescence_data(file_path):
    try:
        data = pd.read_excel(file_path)
        print(f"Fluorescence data loaded successfully from {file_path}.")
        return data
    except Exception as e:
        print(f"Error loading fluorescence file: {e}")
        return None

# Load categorization data
def load_categorization_data(file_path):
    try:
        data = pd.read_excel(file_path)
        print(f"Categorization data loaded successfully from {file_path}.")
        return data
    except Exception as e:
        print(f"Error loading categorization file: {e}")
        return None

# Normalize data to [0, 1] range
def normalize_data(df):
    scaler = MinMaxScaler()
    return pd.DataFrame(scaler.fit_transform(df), columns=df.columns, index=df.index)

# Smooth data using a rolling average
def apply_rolling_average(df, window_size= SMOOTH_WINDOW):
    return df.rolling(window=window_size, min_periods=1).mean()

# Find all polygons dynamically
def get_polygons(base_folder):
    files = [f for f in os.listdir(base_folder) if f.endswith("_Accepted_Traces.xlsx")]
    polygons = [os.path.splitext(f)[0].split("_Accepted_Traces")[0] for f in files]
    print("Detected polygons:", polygons)
    return polygons

# Generate heatmaps for multiple polygons
def generate_heatmaps(base_folder):
    polygons = get_polygons(base_folder)

    # Extract base folder name to prepend to polygon identifiers
    base_folder_name = os.path.basename(base_folder)

    # Sort polygons by the desired order
    polygons = sorted(
        polygons,
        key=lambda x: desired_order.index(x.split(f"{base_folder_name}_")[-1])
        if x.split(f"{base_folder_name}_")[-1] in desired_order else len(desired_order)
    )
    print("Polygons after sorting:", polygons)

    heights = []

    for polygon in polygons:
        fluorescence_file_path = os.path.join(base_folder, f"{polygon}_Accepted_Traces.xlsx")
        categorization_file_path = os.path.join(base_folder, f"{polygon}_Categorization.xlsx")

        print(f"Processing polygon: {polygon}")

        fluorescence_data = load_fluorescence_data(fluorescence_file_path)
        categorization_data = load_categorization_data(categorization_file_path)

        if fluorescence_data is None or fluorescence_data.empty:
            print(f"No fluorescence data available for {polygon}. Skipping.")
            continue

        if categorization_data is None or categorization_data.empty:
            print(f"No categorization data available for {polygon}. Skipping.")
            continue

        # Keep only common ROIs between fluorescence data and categorization data
        common_rois = set(fluorescence_data.columns[1:]).intersection(categorization_data["ID"])
        if not common_rois:
            print(f"No matching ROIs between fluorescence and categorization data for {polygon}. Skipping.")
            continue

        fluorescence_data = fluorescence_data[["Frames"] + list(common_rois)]
        categorization_data = categorization_data[categorization_data["ID"].isin(common_rois)]

        # Determine the number of ROIs for dynamic sizing
        roi_count = fluorescence_data.shape[1] - 1  # Exclude the Frames column
        heights.append(roi_count * 0.05)  # Scale height per ROI

    total_height = sum(heights) + len(polygons)  # Add space between heatmaps

    fig, axes = plt.subplots(len(polygons), 1, figsize=(12, total_height), gridspec_kw={'height_ratios': heights})

    for idx, polygon in enumerate(polygons):
        fluorescence_file_path = os.path.join(base_folder, f"{polygon}_Accepted_Traces.xlsx")
        categorization_file_path = os.path.join(base_folder, f"{polygon}_Categorization.xlsx")

        print(f"Plotting heatmap for polygon: {polygon}")

        fluorescence_data = load_fluorescence_data(fluorescence_file_path)
        categorization_data = load_categorization_data(categorization_file_path)

        if fluorescence_data is None or fluorescence_data.empty:
            print(f"No fluorescence data available for {polygon}. Skipping.")
            continue

        if categorization_data is None or categorization_data.empty:
            print(f"No categorization data available for {polygon}. Skipping.")
            continue

        # Keep only common ROIs between fluorescence data and categorization data
        common_rois = set(fluorescence_data.columns[1:]).intersection(categorization_data["ID"])
        if not common_rois:
            print(f"No matching ROIs between fluorescence and categorization data for {polygon}. Skipping.")
            continue

        fluorescence_data = fluorescence_data[["Frames"] + list(common_rois)]
        categorization_data = categorization_data[categorization_data["ID"].isin(common_rois)]

        # Separate Frames and ROI data
        frames = fluorescence_data.iloc[:, 0]
        roi_data = fluorescence_data.iloc[:, 1:]

        # Set Frames as the index
        roi_data.index = frames

        # Apply smoothing and normalization
        smoothed_data = apply_rolling_average(roi_data)
        normalized_data = normalize_data(smoothed_data)

        # Reorder ROIs based on categorization
        category_order = ["Excitation", "Inhibition", "Non-Responding"]
        categorization_data["Category"] = pd.Categorical(
            categorization_data["Category"], categories=category_order, ordered=True
        )
        categorization_data = categorization_data.sort_values(by=["Category", "ID"])

        ordered_rois = categorization_data["ID"].values
        ordered_data = normalized_data[ordered_rois]

        # Identify ROIs by category
        non_responding_rois = categorization_data[categorization_data["Category"] == "Non-Responding"]["ID"].values
        excitation_rois = categorization_data[categorization_data["Category"] == "Excitation"]["ID"].values
        inhibition_rois = categorization_data[categorization_data["Category"] == "Inhibition"]["ID"].values

        # Transpose for heatmap (ROIs on y-axis, Frames on x-axis)
        transposed_data = ordered_data.T

        # Plot the heatmap
        ax = axes[idx]
        sns.heatmap(
            transposed_data, cmap="magma", cbar=True,
            yticklabels=1, cbar_kws={'label': 'Fluorescence Intensity', 'shrink': 0.5}, ax=ax
        )
        ax.set_title(f"{polygon}", fontsize=16)
        ax.set_xlabel("Frames", fontsize=14)
        ax.set_ylabel("ROIs", fontsize=14, labelpad=60)
        ax.tick_params(axis='y', labelsize=4)

        # Add rectangles next to ROI labels based on category
        for ytick, roi in enumerate(transposed_data.index):
            if roi in non_responding_rois:
                ax.add_patch(Rectangle((-150, ytick - 0.5), 40, 1, color="#7F7F7F", transform=ax.transData, clip_on=False))
            elif roi in excitation_rois:
                ax.add_patch(Rectangle((-150, ytick - 0.5), 40, 1, color="#7fbf7b", transform=ax.transData, clip_on=False))
            elif roi in inhibition_rois:
                ax.add_patch(Rectangle((-150, ytick - 0.5), 40, 1, color="#af8dc3", transform=ax.transData, clip_on=False))


        # Save the heatmap for the current polygon as a TIFF file with uniform ROI height, without labels or legends
        roi_height = 0.15  # Set uniform height for each ROI
        individual_tiff_output_file = os.path.join(base_folder, f"{polygon}_Category_Heatmap_min_max_Norm.tiff")
        plt.figure(figsize=(20, roi_height * transposed_data.shape[0]))
        sns.heatmap(
            transposed_data, cmap="magma", cbar=False, xticklabels=False, yticklabels=False
        )
        plt.axis('off')
        plt.savefig(individual_tiff_output_file, format='tiff', dpi=100, bbox_inches='tight', pad_inches=0)
        print(f"Individual heatmap saved to {individual_tiff_output_file}")
        plt.close()

    plt.tight_layout()

    # Save the combined heatmaps as a low-resolution PDF
    pdf_output_file = os.path.join(base_folder, f"{os.path.basename(base_folder)}_Category_Heatmaps_min_max_Norm.pdf")
    plt.savefig(pdf_output_file, format='pdf')
    print(f"Heatmaps saved to {pdf_output_file}")

    # Save the combined heatmaps as a TIFF
    tiff_output_file = os.path.join(base_folder, f"{os.path.basename(base_folder)}_Category_Heatmaps_min_max_Norm.tiff")
    plt.savefig(tiff_output_file, format='tiff', dpi=300)
    print(f"Heatmaps saved to {tiff_output_file}")

    plt.show()

# Main execution
generate_heatmaps(base_folder)


# %% [code]
#generate trace plots (overlayed) - min/max normalisation


region_colors = {
    'OGL': '#8DA0CB',
    'IGL': '#FC8D62',
    'Med': '#66C2A5'
}

# -----------------------------
# IO
# -----------------------------
def load_fluorescence_data(file_path):
    try:
        return pd.read_excel(file_path)
    except Exception as e:
        print(f"Error loading fluorescence file: {e}")
        return None


def load_categorization_data(file_path):
    try:
        return pd.read_excel(file_path)
    except Exception as e:
        print(f"Error loading categorization file: {e}")
        return None


# -----------------------------
# PROCESSING
# -----------------------------
def normalize_data(df):
    scaler = MinMaxScaler()
    return pd.DataFrame(
        scaler.fit_transform(df),
        columns=df.columns,
        index=df.index
    )


def apply_rolling_average(df, window_size=SMOOTH_WINDOW):
    return df.rolling(window=window_size, min_periods=1).mean()


def get_polygons(base_folder):
    files = [f for f in os.listdir(base_folder) if f.endswith("_Accepted_Traces.xlsx")]
    return [os.path.splitext(f)[0].split("_Accepted_Traces")[0] for f in files]


def get_region_from_polygon(polygon_name):
    for region in region_colors.keys():
        if region in polygon_name:
            return region
    return None


def plot_overlay_traces(ax, data, polygon_name):
    for col in data.columns:
        ax.plot(data.index, data[col], color="grey", alpha=0.2, linewidth=0.5)

    region = get_region_from_polygon(polygon_name)
    mean_color = region_colors.get(region, "black")

    mean_trace = data.mean(axis=1)
    ax.plot(data.index, mean_trace, color=mean_color, linewidth=2.5)

    ax.set_title(polygon_name)
    ax.set_xlabel("Frames")
    ax.set_ylabel("Normalized Intensity")


# -----------------------------
# MAIN
# -----------------------------
def generate_overlay_plots(base_folder):

    polygons = get_polygons(base_folder)
    base_folder_name = os.path.basename(base_folder)

    polygons = sorted(
        polygons,
        key=lambda x: desired_order.index(x.split(f"{base_folder_name}_")[-1])
        if x.split(f"{base_folder_name}_")[-1] in desired_order else len(desired_order)
    )

    heights = []
    valid_polygons = []

    # Store data per region for global means
    region_data = {region: [] for region in region_colors.keys()}

    # -------- Pass 1: sizing + collect region data --------
    for polygon in polygons:
        fluorescence_path = os.path.join(base_folder, f"{polygon}_Accepted_Traces.xlsx")
        categorization_path = os.path.join(base_folder, f"{polygon}_Categorization.xlsx")

        fluorescence_data = load_fluorescence_data(fluorescence_path)
        categorization_data = load_categorization_data(categorization_path)

        if fluorescence_data is None or fluorescence_data.empty:
            continue
        if categorization_data is None or categorization_data.empty:
            continue

        common_rois = set(fluorescence_data.columns[1:]).intersection(categorization_data["ID"])
        if not common_rois:
            continue

        fluorescence_data = fluorescence_data[["Frames"] + list(common_rois)]

        frames = fluorescence_data.iloc[:, 0]
        roi_data = fluorescence_data.iloc[:, 1:]
        roi_data.index = frames

        smoothed = apply_rolling_average(roi_data)
        normalized = normalize_data(smoothed)

        region = get_region_from_polygon(polygon)
        if region in region_data:
            region_data[region].append(normalized)

        heights.append(max(2.5, len(common_rois) * 0.02))
        valid_polygons.append(polygon)

    if not valid_polygons:
        print("No valid polygons found.")
        return

    # -------- Pass 2: overlay plots --------
    fig, axes = plt.subplots(
        len(valid_polygons),
        1,
        figsize=(12, sum(heights) + len(valid_polygons)),
        gridspec_kw={"height_ratios": heights}
    )

    if len(valid_polygons) == 1:
        axes = [axes]

    for idx, polygon in enumerate(valid_polygons):
        fluorescence_path = os.path.join(base_folder, f"{polygon}_Accepted_Traces.xlsx")
        categorization_path = os.path.join(base_folder, f"{polygon}_Categorization.xlsx")

        fluorescence_data = load_fluorescence_data(fluorescence_path)
        categorization_data = load_categorization_data(categorization_path)

        common_rois = set(fluorescence_data.columns[1:]).intersection(categorization_data["ID"])

        fluorescence_data = fluorescence_data[["Frames"] + list(common_rois)]
        frames = fluorescence_data.iloc[:, 0]
        roi_data = fluorescence_data.iloc[:, 1:]
        roi_data.index = frames

        smoothed = apply_rolling_average(roi_data)
        normalized = normalize_data(smoothed)

        ax = axes[idx]
        plot_overlay_traces(ax, normalized, polygon)

    plt.tight_layout()

    pdf_output = os.path.join(base_folder, f"{base_folder_name}_Overlay_Traces_min_max_norm.pdf")
    plt.savefig(pdf_output)
    print(f"Saved: {pdf_output}")

    tiff_output = os.path.join(base_folder, f"{base_folder_name}_Overlay_Traces_min_max_Norm.tiff")
    plt.savefig(tiff_output, dpi=300)
    print(f"Saved: {tiff_output}")

    plt.show()

    # -------- New: REGION MEAN PLOT --------
    plt.figure(figsize=(10, 6))

    for region, datasets in region_data.items():
        if not datasets:
            continue

        # Concatenate all ROIs across polygons
        combined = pd.concat(datasets, axis=1)

        # Mean across all ROIs
        region_mean = combined.mean(axis=1)

        plt.plot(
            region_mean.index,
            region_mean,
            color=region_colors[region],
            linewidth=3,
            label=region
        )

    plt.xlabel("Frames")
    plt.ylabel("Normalized Intensity")
    plt.title("Region Mean Responses")
    plt.legend()

    region_pdf = os.path.join(base_folder, f"{base_folder_name}_Region_MeanTraces_min_max_Norm.pdf")
    plt.savefig(region_pdf)
    print(f"Saved: {region_pdf}")

    region_tiff = os.path.join(base_folder, f"{base_folder_name}_Region_MeanTraces_min_max_Norm.tiff")
    plt.savefig(region_tiff, dpi=300)
    print(f"Saved: {region_tiff}")

    plt.show()


# -----------------------------
# RUN
# -----------------------------
generate_overlay_plots(base_folder)
```
