## Supplemental file 3 for "Protocol for calcium imaging of acute brain slices from *Octopus vulgaris* hatchlings during application of neurotransmitters": Octo_CaImaging_Part3_ExcitatorySlope.ipynb

```
# %% [code]
import os
import sys
from pathlib import Path

import numpy as np
import pandas as pd

import matplotlib as mpl
import matplotlib.pyplot as plt
from matplotlib.gridspec import GridSpec

import seaborn as sns
import cmasher as cmr

from scipy.ndimage import gaussian_filter1d
import scipy.stats as stats

import statsmodels.api as sm
import statsmodels.formula.api as smf
import statsmodels.stats.multicomp as mc

import pingouin as pg

from IPython.display import display
from ipyfilechooser import FileChooser
import ipywidgets as widgets
import copy

from PIL import Image
from ipyfilechooser import FileChooser
from IPython.display import display
import ipywidgets as widgets

DEFAULT_MAIN = r"C:/Users\Amy Courtney\Documents/Temp/Calcium_Imaging_Paper/Biorxiv_Submission1_Mar26/Supp_Files/Courtney_et_al_2026_Supp_File_3_Sample_Dataset/Calcium_Imaging_Analysis"
DEFAULT_BASE = r"C:/Users\Amy Courtney\Documents/Temp/Calcium_Imaging_Paper/Biorxiv_Submission1_Mar26/Supp_Files/Courtney_et_al_2026_Supp_File_3_Sample_Dataset/Calcium_Imaging_Analysis"

main_fc = FileChooser(DEFAULT_MAIN, select_desc="Main Folder")
main_fc.show_only_dirs = True

base_fc = FileChooser(DEFAULT_BASE, select_desc="Single Video Folder")
base_fc.show_only_dirs = True

display(main_fc)
display(base_fc)

print(display)
print(widgets)
print(FileChooser)


# %% [code]
main_folder = Path(main_fc.selected_path).resolve()
base_folder = Path(base_fc.selected_path).resolve()

print("Main folder:", main_folder)
print("Base folder:", base_folder)

# %% [code]
lowest_folder = os.path.basename(os.path.normpath(base_folder))
###order
desired_order = ['OGL', 'IGL', 'Med', 'CB']
###colours
region_colors = {'OGL': '#8da0cb', 'IGL': '#66c2a5', 'Med': '#fc8d62', 'CB': '#e78ac3'}

# %% [code]
FR=5
Slope_win= 10
End_win=400
baseline_frames = 30 # First 50 frames in stimulus window are baseline
Smooth_win=20

# %% [code]
# List to store paths of matching files
matching_files = []

# Walk through the directory to find matching files
for root, _, files in os.walk(base_folder):
    for file in files:
        if file.endswith('_Accepted_Traces.xlsx'):
            matching_files.append(os.path.join(root, file))

# Load data from all matching files
if matching_files:
    print("Matching files found:")
    for file_path in matching_files:
        print(file_path)

    # Initialize an empty DataFrame to concatenate columns
    combined_data = pd.DataFrame()
    
    for file_path in matching_files:
        # Extract file name without the base folder and '_Accepted_Traces.xlsx' suffix
        relative_path = os.path.relpath(file_path, base_folder)
        filename_core = relative_path.replace('_Accepted_Traces.xlsx', '')
        
        # Load the data
        data = pd.read_excel(file_path)
        
        # Update column names to include filename core
        data.columns = [f"{col}_{filename_core}" for col in data.columns]
        
        # Concatenate columns from the new file to the combined DataFrame
        combined_data = pd.concat([combined_data, data], axis=1)
        print(f"Data added from: {file_path}")
    
    print("All data combined successfully.")
else:
    print("No matching files found.")


# %% [code]
# Initialize an empty list to store DataFrames
dataframes = []

# Loop through all files in the base folder
for file_name in os.listdir(base_folder):
    # Check if the file ends with '_Categorization.xlsx'
    if file_name.endswith('_Categorization.xlsx'):
        # Construct the full file path
        file_path = os.path.join(base_folder, file_name)
        
        # Read the Excel file into a DataFrame
        try:
            df = pd.read_excel(file_path)
            dataframes.append(df)
        except Exception as e:
            print(f"Error reading {file_name}: {e}")

# Combine all DataFrames into one
Categories = pd.concat(dataframes, ignore_index=True)

# Display the resulting DataFrame
print(Categories)


# %% [code]
# Filter the DataFrame to keep only rows where the 'Category' column is 'Non-Responding' or 'Inhibition'
filtered_categories = Categories[Categories['Category'].isin(['Non-Responding', 'Inhibition'])]

# Display the filtered DataFrame
print(filtered_categories)

# %% [code]
# Iterate over each ID in the 'ID' column of 'filtered_categories'
for id_string in filtered_categories['ID']:
    # Check if any column in 'combined_data' contains the ID string before the first '_'
    columns_to_remove = [
        col for col in combined_data.columns 
        if col.split('_')[0] == id_string
    ]
    
    # Drop the identified columns from 'combined_data'
    combined_data.drop(columns=columns_to_remove, inplace=True)

# %% [code]
#define Dopamine stimulus start time in frames
stimulus_win_min = 250
print(f"stimulus_win_min: {stimulus_win_min}")

# %% [code]
stimulus_win_max=stimulus_win_min+End_win
combined_data = combined_data.loc[:, ~combined_data.columns.str.contains('Frames', case=False)]
combined_data_col_names = combined_data.columns
print(combined_data_col_names)
fcorr= combined_data
fcorr = fcorr.T
fcorr = fcorr.to_numpy()

# %% [code]
# Create the IDs DataFrame from column names
IDs = pd.DataFrame({
    'Regions': combined_data_col_names
})

# Split the 'Regions' column to extract ROI and last part
IDs['ROI'] = IDs['Regions'].str.split('_').str[0]
IDs['Regions'] = IDs['Regions'].str.split('_').str[-1]

# Additional processing for 'left' or 'right' in the 'Regions' string
for index, row in IDs.iterrows():
    if 'left' in row['Regions'] or 'right' in row['Regions']:
        # Get the original string
        original_string = combined_data_col_names[index]
        
        # Split the original string into parts
        parts = original_string.split('_')
        
        if len(parts) > 3:
            # Extract the string between the second-to-last and third-to-last '_'
            in_between = parts[-2]
            
            # Extract whether it's 'left' or 'right'
            left_or_right = row['Regions']
            
            # Update the Regions value to the new format
            updated_region = f"{in_between}_{left_or_right}"
            IDs.at[index, 'Regions'] = updated_region

# Print the resulting DataFrame
print(IDs)


# %% [code]
#plot a subset of ROI
Num_Plots=3
# File names
pdf_file_path = os.path.join(base_folder, f"{lowest_folder}_LigSlope_Ordered.pdf")
tiff_file_path = os.path.join(base_folder, f"{lowest_folder}_LigSlope_Ordered.tiff")

def normalize_trace(traces, stim_start, stim_end, baseline_frames=50):
    """
    Normalize entire traces between 0 and 1 after baseline correction.
    - `traces`: 2D NumPy array (num_traces, num_timepoints)
    - `stim_start`: Start frame for normalization
    - `stim_end`: End frame for normalization
    - `baseline_frames`: Number of frames to use for F0 calculation
    """
    # Compute F0 (5th percentile of the first 50 frames in the stimulus window)
    F0 = np.percentile(traces[:, stim_start:stim_start + baseline_frames], q=5, axis=1, keepdims=True)

    # Subtract F0 from the entire trace (baseline correction)
    traces_corrected = traces - F0

    # Find min and max values across the entire trace for each ROI
    min_val = np.min(traces_corrected, axis=1, keepdims=True)
    max_val = np.max(traces_corrected, axis=1, keepdims=True)

    # Normalize entire trace between 0 and 1
    normalized_traces = (traces_corrected - min_val) / (max_val - min_val + 1e-6)  # Avoid division by zero

    return normalized_traces


# Function to smooth traces using a moving average filter
def smooth_trace(traces, window_size=Smooth_win):
    kernel = np.ones(window_size) / window_size
    return np.apply_along_axis(lambda m: np.convolve(m, kernel, mode='same'), axis=1, arr=traces)

# Vectorized function to calculate max slopes
def calculate_max_slope(traces, stim_start=stimulus_win_min, stim_end=stimulus_win_max, window_size=Slope_win):
    num_traces, trace_length = traces.shape
    max_slopes = np.full(num_traces, -np.inf)
    max_slope_starts = np.full(num_traces, stim_start)

    for start in range(stim_start, stim_end - window_size + 1):
        end = start + window_size
        y_windows = traces[:, start:end]
        slopes = (y_windows[:, -1] - y_windows[:, 0]) / (window_size - 1) * FR
        
        improved_mask = slopes > max_slopes
        max_slopes[improved_mask] = slopes[improved_mask]
        max_slope_starts[improved_mask] = start

    return max_slopes, max_slope_starts


# Normalize and smooth traces with full normalization between 0 and 1
normalized_traces = normalize_trace(fcorr, stimulus_win_min, stimulus_win_max)
smoothed_traces = smooth_trace(normalized_traces)

# Calculate max slopes
slopes, max_slope_starts = calculate_max_slope(smoothed_traces)

# Create DataFrame
df_slopes = pd.DataFrame({
    "ROI": IDs['ROI'],
    "Regions": IDs['Regions'],
    "Slope": slopes,
    "Trace": list(smoothed_traces),
    "MaxSlopeStart": max_slope_starts
})

# Sort DataFrame by slope in descending order
df_slopes = df_slopes.sort_values(by="Slope", ascending=False).reset_index(drop=True)

# Extract every 5th sorted slope, trace, and max slope start
#sorted_slopes = df_slopes["Slope"].values[::Num_Plots]  # Take every 5th element
#sorted_traces = np.array(df_slopes["Trace"].tolist())[::Num_Plots]  
#sorted_rois = df_slopes["ROI"].values[::Num_Plots]
#sorted_max_slope_starts = df_slopes["MaxSlopeStart"].values[::Num_Plots]

# Extract every 5th sorted slope, trace, and max slope start
sorted_slopes = df_slopes["Slope"].values  # Take every 5th element
sorted_traces = np.array(df_slopes["Trace"].tolist())  
sorted_rois = df_slopes["ROI"].values
sorted_max_slope_starts = df_slopes["MaxSlopeStart"].values

# Adjust the number of plots
num_selected_rois = len(sorted_slopes)
plots_per_row = 10
num_rows = int(np.ceil(num_selected_rois / plots_per_row))

fig, axes = plt.subplots(num_rows, plots_per_row, figsize=(15, num_rows * 2), sharex=True, sharey=True)
axes = axes.flatten()

for i in range(num_selected_rois):
    trace = sorted_traces[i]
    slope = sorted_slopes[i] 
    max_slope_start = sorted_max_slope_starts[i]

    # Plot the smoothed and normalized trace
    axes[i].plot(trace)
    axes[i].set_title(f"{sorted_rois[i]}\nSlope: {slope:.5f}", fontsize=8)
    
    # Mark the max slope region
    axes[i].axvline(max_slope_start, color='red', linestyle='-', linewidth=1, label="Max Slope Start")
    axes[i].axvline(max_slope_start + 10, color='red', linestyle='-', linewidth=1)

    # Set x-axis limits
    axes[i].set_xlim([stimulus_win_min, stimulus_win_max])

    axes[i].tick_params(axis='both', which='both', labelsize=6)
    axes[i].legend(fontsize=6)

# Hide empty subplots
for j in range(num_selected_rois, len(axes)):
    axes[j].axis('off')

plt.tight_layout()
# Save the plot
plt.savefig(pdf_file_path, format='pdf')
plt.savefig(tiff_file_path, format='tiff', dpi=300)


# %% [code]
# Modify the Regions column to keep only the string before '_'
df_slopes['Regions'] = df_slopes['Regions'].str.split('_').str[0]

# %% [code]
#plot every second shallow trace

def plot_selected_traces(df_slopes, base_folder, lowest_folder):
    if 'Trace' not in df_slopes.columns or 'Slope' not in df_slopes.columns:
        raise ValueError("The dataframe must contain both 'Trace' and 'Slope' columns.")
    
    selected_rows = df_slopes[(df_slopes['Slope'] >= 0.025) & (df_slopes['Slope'] <= 0.075)]
    
    if selected_rows.empty:
        raise ValueError("No traces with 'Slope' in the specified range were found.")
    
    plt.figure(figsize=(10, 5))
    
    traces = []
    selected_rows = selected_rows.iloc[::6]  # Select every second trace
    
    for _, row in selected_rows.iterrows():
        trace = row['Trace']
        
        if isinstance(trace, str):
            try:
                trace = np.array(eval(trace))  # Convert from string representation if needed
            except Exception as e:
                raise ValueError("Error converting Trace column to array:", e)
        
        if not isinstance(trace, (list, np.ndarray, pd.Series)):
            raise TypeError("The 'Trace' column should contain list-like time-series data.")
        
        trace_segment = trace[stimulus_win_min:(stimulus_win_min+End_win)]
        traces.append(trace_segment)
        plt.plot(trace_segment, color='black', linewidth=0.1)
    
    mean_trace = np.mean(traces, axis=0)
    plt.plot(mean_trace, color='#7FBF7B', linewidth=1)
    
    plt.xlabel("Time (frames or seconds)")
    plt.ylabel("Fluorescence Intensity")
    plt.title("")
    
    # Save the plot
    pdf_file_path = os.path.join(base_folder, f"{lowest_folder}_LigSlope_Traces_Shallow_Subset.pdf")
    tiff_file_path = os.path.join(base_folder, f"{lowest_folder}_LigSlope_Traces_Shallow_Subset.tiff")
    
    plt.savefig(pdf_file_path, format='pdf')
    plt.savefig(tiff_file_path, format='tiff', dpi=300)
    
    plt.show()

# Example usage
plot_selected_traces(df_slopes, base_folder, lowest_folder)


# %% [code]
# File names
pdf_file_path = os.path.join(base_folder, f"{lowest_folder}_LigSlope_Traces_Steep.pdf")
tiff_file_path = os.path.join(base_folder, f"{lowest_folder}_LigSlope_Traces_Steep.tiff")

# Assuming df_slopes is already loaded as a DataFrame
def plot_selected_traces(df_slopes):
    if 'Trace' not in df_slopes.columns or 'Slope' not in df_slopes.columns:
        raise ValueError("The dataframe must contain both 'Trace' and 'Slope' columns.")
    
    selected_rows = df_slopes[df_slopes['Slope'] >= 0.13]
    
    if selected_rows.empty:
        raise ValueError("No traces with 'Slope' above 0.1 were found.")
    
    plt.figure(figsize=(10, 5))
    
    traces = []
    for _, row in selected_rows.iterrows():
        trace = row['Trace']
        
        if isinstance(trace, str):
            try:
                trace = np.array(eval(trace))  # Convert from string representation if needed
            except Exception as e:
                raise ValueError("Error converting Trace column to array:", e)
        
        if not isinstance(trace, (list, np.ndarray, pd.Series)):
            raise TypeError("The 'Trace' column should contain list-like time-series data.")
        
        trace_segment = trace[stimulus_win_min:(stimulus_win_min+End_win)]
        traces.append(trace_segment)
        plt.plot(trace_segment, color='black', linewidth=0.1)
    
    mean_trace = np.mean(traces, axis=0)
    plt.plot(mean_trace, color='#7FBF7B', linewidth=1)
    
    plt.xlabel("Time (frames or seconds)")
    plt.ylabel("Fluorescence Intensity")
    plt.title("")

    # Save the plot
    plt.savefig(pdf_file_path, format='pdf')
    plt.savefig(tiff_file_path, format='tiff', dpi=300)
    plt.show()

# Example usage
plot_selected_traces(df_slopes)


# %% [code]
# Extract the lowest folder name from base_folder
lowest_folder = os.path.basename(os.path.normpath(base_folder))

# File names
pdf_file_path = os.path.join(base_folder, f"{lowest_folder}_LigSlope_perRegion.pdf")
tiff_file_path = os.path.join(base_folder, f"{lowest_folder}_LigSlope_perRegion.tiff")

# Create the figure
plt.figure(figsize=(8, 6))

# Violin plot (distribution) - Background layer
sns.violinplot(
    data=df_slopes,
    x="Regions",
    y="Slope",
    order=desired_order,
    palette=region_colors,
    inner=None,  # Hide inner lines to prevent overlap with strip plot
    alpha=0.5  # Transparency for visibility
)

# Strip plot (individual data points) - Middle layer
sns.stripplot(
    data=df_slopes,
    x="Regions",
    y="Slope",
    order=desired_order,
    palette=region_colors,
    jitter=True,
    size=8,
    alpha=0.7
)

# Point plot (mean values) - Top layer
sns.pointplot(
    data=df_slopes,
    x="Regions",
    y="Slope",
    order=desired_order,
    color="grey",  # Mean values in grey
    markers="D",  # Diamond marker for mean
    scale=1.2,  # Slightly larger marker
    errwidth=1.5,  # Error bar width
    capsize=0.2,  # Small caps on error bars
    join=False,
    estimator=np.mean,
    zorder=3  # Ensure point plot is on top
)

# Add a horizontal line at slope = 0.004
plt.axhline(y=0.075, color='black', linestyle='--', linewidth=1.5, label="Threshold (0.10)")

# Customizing the plot
plt.title("Slopes with Mean Values", fontsize=14)
plt.xlabel("", fontsize=12)
plt.ylabel("Slope", fontsize=12)
plt.xticks(fontsize=10)
plt.yticks(fontsize=10)
plt.grid(axis='y', linestyle='--', alpha=0.6)

# Adjust layout
plt.tight_layout()

# Save the plot
plt.savefig(pdf_file_path, format='pdf')
plt.savefig(tiff_file_path, format='tiff', dpi=300)

# Show the plot
# plt.show()


# %% [code]
# Check Normality (Shapiro-Wilk test for each region)
normality_results = df_slopes.groupby("Regions")["Slope"].apply(lambda x: stats.shapiro(x)[1])  # p-values
print("Normality Test (Shapiro-Wilk):\n", normality_results)

# Check Homogeneity of Variance (Levene's test)
levene_p = stats.levene(*[group["Slope"].values for name, group in df_slopes.groupby("Regions")])[1]
print(f"Levene’s test for equality of variances: p = {levene_p}")

# One-way ANOVA if normality holds and variances are equal
if all(p > 0.05 for p in normality_results) and levene_p > 0.05:
    anova_result = stats.f_oneway(*[group["Slope"].values for name, group in df_slopes.groupby("Regions")])
    print("One-way ANOVA:\n", anova_result)

    # Post-hoc test: Tukey's HSD
    tukey = mc.pairwise_tukeyhsd(df_slopes["Slope"], df_slopes["Regions"])
    print("Tukey's HSD Results:\n", tukey)

# Welch’s ANOVA if variances are unequal
elif levene_p < 0.05:
    welch_anova = pg.welch_anova(dv="Slope", between="Regions", data=df_slopes)
    print("Welch’s ANOVA:\n", welch_anova)

    # Games-Howell Post-hoc test
    games_howell = pg.pairwise_gameshowell(dv="Slope", between="Regions", data=df_slopes)
    print("Games-Howell Post-hoc Test:\n", games_howell)

# Kruskal-Wallis test if normality is violated
else:
    kw_result = stats.kruskal(*[group["Slope"].values for name, group in df_slopes.groupby("Regions")])
    print("Kruskal-Wallis Test:\n", kw_result)

    # Dunn’s Post-hoc test
    dunn_test = pg.pairwise_tests(dv="Slope", between="Regions", data=df_slopes, padjust="bonferroni")
    print("Dunn's Test Results:\n", dunn_test)


# %% [code]

# Extract the lowest folder name from base_folder
lowest_folder = os.path.basename(os.path.normpath(main_folder))
region_colors = {'OGL': '#8da0cb', 'IGL': '#66c2a5', 'Med': '#fc8d62'}
desired_order = ['OGL', 'IGL', 'Med']

# File names
pdf_file_path = os.path.join(base_folder, f"{lowest_folder}_LigSlope_perRegion.pdf")
tiff_file_path = os.path.join(base_folder, f"{lowest_folder}_LigSlope_perRegion.tiff")

# Create the figure
plt.figure(figsize=(8, 6))

# Plot KDE for each region
for region in desired_order:
    subset = df_slopes[df_slopes["Regions"] == region]
    sns.kdeplot(
        data=subset, 
        x="Slope", 
        label=region, 
        color=region_colors[region], 
        fill=True,  # Fill under the KDE curve
        alpha=0.4,  # Transparency for better visualization
        linewidth=2
    )

# Customizing the plot
plt.title("Kernel Density Estimate of Slopes", fontsize=14)
plt.xlabel("Slope", fontsize=12)
plt.ylabel("Density", fontsize=12)
plt.xticks(fontsize=10)
plt.yticks(fontsize=10)
plt.grid(axis='both', linestyle='--', alpha=0.6)

# Add legend
plt.legend(title="Regions", fontsize=10, loc="upper right")

# Adjust layout
plt.tight_layout()

# Save the plot
plt.savefig(pdf_file_path, format='pdf')
plt.savefig(tiff_file_path, format='tiff', dpi=300)

# Show the plot
plt.show()


# %% [code]
# Categorizing slopes
df_slopes['Slope_Category'] = df_slopes['Slope'].apply(lambda x: 'Steep Slope' if x > 0.075 else 'Shallow Slope')


# %% [code]
# File names
pdf_file_path = os.path.join(base_folder, f"{lowest_folder}_LigSlope_Category_perRegion.pdf")
tiff_file_path = os.path.join(base_folder, f"{lowest_folder}_LigSlope_Category_perRegion.tiff")

# Define response order
slope_cat_order = ['Steep Slope', 'Shallow Slope']

# Define colors for categories
category_colors = {
    'Shallow Slope': '#1f78b4', 
    'Steep Slope': '#33a02c' 
}

# Count occurrences of each Slope_Category per Region
category_counts = df_slopes.groupby(['Regions', 'Slope_Category']).size().unstack(fill_value=0)

# Convert counts to proportions
category_proportions = category_counts.div(category_counts.sum(axis=1), axis=0)

# Reorder index based on desired order
category_proportions = category_proportions.reindex(desired_order)

# Reorder columns based on slope category order
category_proportions = category_proportions.reindex(columns=slope_cat_order, fill_value=0)

# Create a grouped bar plot to show proportions side by side with custom colors
ax = category_proportions.plot(kind='bar', stacked=False, figsize=(8, 6), color=[category_colors[col] for col in category_proportions.columns])

plt.ylabel("Proportion")
plt.xlabel("")
plt.title("Proportion of Slope Categories per Region")
plt.legend(title="Slope Category")
plt.grid(axis='y', linestyle='--', alpha=0.7)

# Save the plot
plt.savefig(pdf_file_path, format='pdf')
plt.savefig(tiff_file_path, format='tiff', dpi=300)

plt.show()


# %% [code]
# List to store paths of matching files
matching_files = []

# Walk through the directory to find matching files
for root, _, files in os.walk(base_folder):
    for file in files:
        if file.endswith('_ROIxy.xlsx'):
            matching_files.append(os.path.join(root, file))

# Load the first matching file if it exists
if matching_files:
    print("Matching files found:")
    for file_path in matching_files:
        print(file_path)
    
    # Load the first matching file
    ROI_Location = pd.read_excel(matching_files[0])
    print("Data loaded from:", matching_files[0])
else:
    print("No matching files found.")


# %% [code]
# Merge the DataFrames
props_df = pd.merge(
    df_slopes,
    ROI_Location[['name', 'x', 'y']],
    left_on='ROI',
    right_on='name',
    how='inner'  # Use 'inner' to ensure only matching rows are included
)

# Drop the redundant 'name' column from the resulting DataFrame
props_df.drop(columns=['name'], inplace=True)


# %% [code]
def make_linear_colormap(array, cmap='inferno', desat=1, reference_numbers=None, percentile=100):
    '''
    Create color map - linear version. 
    Uses seaborn color_palette() function to create color map. 
    
    Paramters
    ---------
        - array: 1-D Numpy array to loop over and create colors for
        - cmap: Color palette name 
        - desat: seaborn specific desaturation of color palette. 1 = no desaturation.
        - reference_numbers: reference array (to use instead of input array)
        - percentile: To get rid of outlier artefacts, use percentile < 100 as maximum    
    Returns
    -------
        - colors: array of length len(array) of specified palette
    '''
    
    array = np.nan_to_num(array.copy())
    
    if reference_numbers is not None:
        reference_array = np.linspace(reference_numbers.min(),\
                                      reference_numbers.max(),\
                                      len(array))
    else:
        reference_array = None
        
    # Correct for negative values in array
    if (array < 0).any():
        minimum = array.min()
        array += -1 * minimum
        if reference_array is not None: # Also correct reference number range
            reference_array += -1 * minimum
            
    color_palette = sns.color_palette(cmap, len(array), desat)
    colors = []
    for el in array:
        if reference_array is None:
            index = int((el/np.percentile(array, percentile))*len(color_palette))
        else:
            index = int((el/np.percentile(reference_array, percentile))*len(color_palette))
            
        if index > len(color_palette)-1: index = -1
        color = color_palette[index]
        colors.append(color)
        
    return np.array(colors)


def make_colorbar(array, no_steps=5, font_scale=10, cmap='inferno', **kwargs):
    ''' Draw a colorbar that can be copy+pasted next to another graph '''
    
    return_figure = kwargs.get('return_figure', None)
    show_labels = kwargs.get('show_labels', True)
    
    start_value = np.min(array)
    end_value = np.max(array)

    labels = [''] * no_steps
    labels[0] = '{:.1f}'.format(start_value)
    labels[-1]= '{:.1f}'.format(end_value)
    figure = plt.figure();
    with sns.axes_style("dark", {"font.size": font_scale}):
        g = sns.palplot(sns.color_palette(cmap, no_steps))
    plt.gca().set_xticklabels(labels,  rotation = -90, ha="center", va='top')
    if not show_labels:
        plt.gca().set_xticklabels([])
    #sns.despine(left=True,bottom=True)

        
    if return_figure:
        return g


# %% [code]
# Choose a property
property_cmap = props_df['Slope'] # centroid-0 is the cell's y position
property_cmap.min(), property_cmap.max()
#colormap = 'inferno'
#colormap = 'cmr.prinsenvlag'
colormap = 'cmr.guppy'
colors_prop = make_linear_colormap(property_cmap, cmap=colormap)
colors_prop.shape # We created 274 colors in RGB format

suite2p_output = base_folder / 'suite2p/plane0'
ops = np.load(suite2p_output / 'ops.npy', allow_pickle= True).item()
image = ops['max_proj']
data=image
import cmasher as cmr

figure = plt.figure(figsize=(15,8))
ax = figure.add_subplot(121)
ax.imshow(data, vmin=0, vmax=np.nanpercentile(data,99.9), cmap='cmr.sunburst')
ax.set_xticks([]); ax.set_yticks([]);
# Plot average and rois 
ax = figure.add_subplot(122)
ax.imshow(data, vmin=0, vmax=np.nanpercentile(data,99.99), cmap='cmr.neutral')
# Create outlines 
for no, roi in enumerate(props_df.iterrows()):
    # Unpack the row data
    _, roi_data = roi
    
    # Get centroid coordinates
    y_centroid = roi_data['x']  # Row coordinate
    x_centroid = roi_data['y']  # Column coordinate
    
    # Plot the centroid as a single dot
    ax.scatter(x_centroid, 
               y_centroid, 
               s=5,        # Adjust size for better visibility
               alpha=0.7,   # Adjust transparency
               color=colors_prop[no])  # Use a unique color per ROI

    
    #ax.text(np.min(outline[1]), np.min(outline[0]), no)

ax.set_xticks([]); ax.set_yticks([]);
plt.tight_layout()

pdf_file_path = os.path.join(base_folder, f"{lowest_folder}_LigSlope_HeatMap_guppy.pdf")
tiff_file_path = os.path.join(base_folder, f"{lowest_folder}_LigSlope_HeatMap_guppy.tiff")

plt.savefig(pdf_file_path, format='pdf', dpi=300)
plt.savefig(tiff_file_path, format='tiff', dpi=300)

# Display the plot
plt.show()

# %% [code]
def make_colorbar(array, no_steps=10, font_scale=1, cmap='hls', **kwargs):
    ''' Draw a colorbar that can be copy+pasted next to another graph '''
    
    return_figure = kwargs.get('return_figure', None)
    show_labels = kwargs.get('show_labels', True)
    
    start_value = np.min(array)
    end_value = np.max(array)

    labels = [''] * no_steps
    labels[0] = '{:.4f}'.format(start_value)
    labels[-1]= '{:.4f}'.format(end_value)
    figure = plt.figure();
    with sns.axes_style("dark", {"font.size": font_scale}):
        g = sns.palplot(sns.color_palette(cmap, no_steps))
    plt.gca().set_xticklabels(labels,  rotation = -90, ha="center", va='top')
    if not show_labels:
        plt.gca().set_xticklabels([])
    #sns.despine(left=True,bottom=True)

        
    if return_figure:
        return g

#  We can also create a separate colorbar that we can copy and paste into the final figure if we want to ... 
make_colorbar(property_cmap, cmap=colormap)


scale_file_path = os.path.join(base_folder, f"{lowest_folder}_LigSlope_HeatMap_Scale_guppy.png")
# Save the plot as a PNG file
plt.savefig(scale_file_path, dpi=300, bbox_inches='tight')  # Adjust dpi and bbox_inches for quality and layout

# Show the plot
plt.show()

# %% [code]
filename = f"{lowest_folder}_Lig_Slope.xlsx"

# Define the full path to save the file
file_path = os.path.join(base_folder, filename)

# Save the DataFrame as an Excel file
props_df.to_excel(file_path, index=False)

print(f"File saved to: {file_path}")
```
