## Supplemental file 3 for "Protocol for calcium imaging of acute brain slices from *Octopus vulgaris* hatchlings during application of neurotransmitters": Octo_CaImaging_Part4_CombineData.ipynb

```
# %% [code]
import os

from pathlib import Path
import numpy as np
import pandas as pd

import matplotlib.pyplot as plt
import seaborn as sns

import scipy.stats as stats

import statsmodels.api as sm
import statsmodels.formula.api as smf
import statsmodels.stats.multicomp as mc

import pingouin as pg
from PIL import Image
from ipyfilechooser import FileChooser
from IPython.display import display
import ipywidgets as widgets

DEFAULT_MAIN = r"C:/Users\Amy Courtney\Documents/Temp/Calcium_Imaging_Paper/Biorxiv_Submission1_Mar26/Supp_Files/Courtney_et_al_2026_Supp_File_3_Sample_Dataset/Calcium_Imaging_Analysis"
DEFAULT_BASE = r"C:/Users\Amy Courtney\Documents/Temp/Calcium_Imaging_Paper/Biorxiv_Submission1_Mar26/Supp_Files/Courtney_et_al_2026_Supp_File_3_Sample_Dataset/Calcium_Imaging_Analysis"

main_fc = FileChooser(DEFAULT_MAIN, select_desc="Main Folder")
main_fc.show_only_dirs = True

base_fc = FileChooser(DEFAULT_BASE, select_desc="where to save")
base_fc.show_only_dirs = True

display(main_fc)
display(base_fc)

print(display)
print(widgets)
print(FileChooser)

# %% [code]
main_folder = Path(main_fc.selected_path).resolve()
base_folder = Path(base_fc.selected_path).resolve()

print("Main folder:", main_folder)
print("Base folder:", base_folder)

# %% [code]
# Initialize an empty list to hold dataframes
dataframes = []

#load dopamine categorisation data

# Walk through the subfolders in the main folder
for root, dirs, files in os.walk(main_folder):
    #change 'Dop' to whatever defines your stimulus group
    if 'Dop' in root:
        for file in files:
            # Check if the file ends with _Categorization.xlsx
            if file.endswith('_Categorization.xlsx'):
                file_path = os.path.join(root, file)
                print(f"Loading file: {file_path}")  # Print the file being loaded
                
                # Extract the region name from the file name
                file_name = os.path.basename(file)
                parts = file_name.split('_')
                if len(parts) >= 3:  # Ensure there are enough parts to extract
                    region_name = parts[-2]
                    # Special case for 'left' or 'right'
                    if region_name.lower() in ['left', 'right'] and len(parts) >= 3:
                        region_name = parts[3]  # Take the string between the third and fourth underscores
                else:
                    region_name = 'Unknown'

                # Load the Excel file into a dataframe
                df = pd.read_excel(file_path)
                
                # Add the Regions column
                df['Regions'] = region_name
                
                # Append the dataframe to the list
                dataframes.append(df)

# Combine all the dataframes into one by rows
combined_df = pd.concat(dataframes, ignore_index=True)

# Display the combined dataframe
print(combined_df)


# %% [code]
# File names
pdf_file_path = os.path.join(Saving_folder, "All_Dop_Slices_Category_Bar.pdf")
tiff_file_path = os.path.join(Saving_folder, "All_Dop_Slices_Category_Bar.tiff")

# Ensure columns are strings
combined_df['Category'] = combined_df['Category'].astype(str)
combined_df['Regions'] = combined_df['Regions'].astype(str)

# Set the order of Regions
region_order = ['OGL', 'IGL', 'Med', 'CB']
#region_order = ['OGL', 'IGL', 'Med']
combined_df['Regions'] = pd.Categorical(combined_df['Regions'], categories=region_order, ordered=True)

# Count occurrences of each Category within each Region
data_counts = combined_df.groupby(['Regions', 'Category']).size().reset_index(name='Count')

# Calculate proportions within each Region
data_counts['Percent'] = data_counts.groupby('Regions')['Count'].transform(lambda x: (x / x.sum()) * 100)

# Calculate total counts for each Region
region_totals = data_counts.groupby('Regions')['Count'].sum()

# Pivot the data to prepare for plotting
plot_data = data_counts.pivot(index='Regions', columns='Category', values='Percent').fillna(0)

# Define custom colors for the categories
category_colors = {
    'Excitation': '#7fbf7b',
    'Inhibition': '#af8dc3',
    'Non-Responding': '#7F7F7F'
}

# Map the colors to the categories in the correct order
colors = [category_colors[cat] for cat in plot_data.columns]

# Plotting the data
fig, ax = plt.subplots(figsize=(10, 6))

# Adjust bar width and edge aesthetics
bars = plot_data.plot(
    kind='bar', 
    figsize=(10, 6), 
    width=0.8, 
    edgecolor='black', 
    color=colors,
    ax=ax
)

# Add grey grid lines
ax.grid(axis='y', color='grey', linestyle='--', linewidth=0.5, alpha=0.7, zorder=0)

# Ensure bars are above grid lines
for bar_group in bars.containers:
    for bar in bar_group:
        bar.set_zorder(3)

# Customize the plot
ax.set_title('Dopamine', fontsize=14)
ax.set_xlabel('', fontsize=12)
ax.set_ylabel('Percent of Cells', fontsize=12)
ax.set_xticks(range(len(region_order)))
ax.set_xticklabels(region_order, fontsize=10, rotation=0)
ax.tick_params(axis='y', labelsize=10)

# Add legend outside the plot
ax.legend(title='Category', fontsize=10, title_fontsize=12, loc='upper left', bbox_to_anchor=(1, 1))

# Add total counts below each region
for idx, region in enumerate(region_order):
    total_count = region_totals.get(region, 0)  # Get total count or 0 if region is missing
    ax.text(idx, -5, f'n={total_count}', ha='center', va='top', fontsize=10, transform=ax.transData)

# Adjust layout to make space for annotations
plt.tight_layout(rect=[0, 0.05, 1, 1])

# Save the plot as PDF and TIFF
plt.savefig(pdf_file_path, format='pdf', bbox_inches='tight')
plt.savefig(tiff_file_path, format='tiff', dpi=300, bbox_inches='tight')

# Display the plot
plt.show()


# %% [code]
# File names
pdf_file_path = os.path.join(Saving_folder, "All_Dop_Slices_Category_Bar_WithoutCB.pdf")
tiff_file_path = os.path.join(Saving_folder, "All_Dop_Slices_Category_Bar_WithoutCB.tiff")

# Ensure columns are strings
combined_df['Category'] = combined_df['Category'].astype(str)
combined_df['Regions'] = combined_df['Regions'].astype(str)

# Set the order of Regions
region_order = ['OGL', 'IGL', 'Med']
combined_df['Regions'] = pd.Categorical(combined_df['Regions'], categories=region_order, ordered=True)

# Count occurrences of each Category within each Region
data_counts = combined_df.groupby(['Regions', 'Category']).size().reset_index(name='Count')

# Calculate proportions within each Region
data_counts['Percent'] = data_counts.groupby('Regions')['Count'].transform(lambda x: (x / x.sum()) * 100)

# Calculate total counts for each Region
region_totals = data_counts.groupby('Regions')['Count'].sum()

# Pivot the data to prepare for plotting
plot_data = data_counts.pivot(index='Regions', columns='Category', values='Percent').fillna(0)

# Define custom colors for the categories
category_colors = {
    'Excitation': '#7fbf7b',
    'Inhibition': '#af8dc3',
    'Non-Responding': '#7F7F7F'
}

# Map the colors to the categories in the correct order
colors = [category_colors[cat] for cat in plot_data.columns]

# Plotting the data
fig, ax = plt.subplots(figsize=(10, 6))

# Adjust bar width and edge aesthetics
bars = plot_data.plot(
    kind='bar', 
    figsize=(10, 6), 
    width=0.8, 
    edgecolor='black', 
    color=colors,
    ax=ax
)

# Add grey grid lines
ax.grid(axis='y', color='grey', linestyle='--', linewidth=0.5, alpha=0.7, zorder=0)

# Ensure bars are above grid lines
for bar_group in bars.containers:
    for bar in bar_group:
        bar.set_zorder(3)

# Customize the plot
ax.set_title('Dopamine', fontsize=14)
ax.set_xlabel('', fontsize=12)
ax.set_ylabel('Percent of Cells', fontsize=12)
ax.set_xticks(range(len(region_order)))
ax.set_xticklabels(region_order, fontsize=10, rotation=0)
ax.tick_params(axis='y', labelsize=10)

# Add legend outside the plot
ax.legend(title='Category', fontsize=10, title_fontsize=12, loc='upper left', bbox_to_anchor=(1, 1))

# Add total counts below each region
for idx, region in enumerate(region_order):
    total_count = region_totals.get(region, 0)  # Get total count or 0 if region is missing
    ax.text(idx, -5, f'n={total_count}', ha='center', va='top', fontsize=10, transform=ax.transData)

# Adjust layout to make space for annotations
plt.tight_layout(rect=[0, 0.05, 1, 1])

# Save the plot as PDF and TIFF
plt.savefig(pdf_file_path, format='pdf', bbox_inches='tight')
plt.savefig(tiff_file_path, format='tiff', dpi=300, bbox_inches='tight')

# Display the plot
plt.show()


# %% [code]
# File names
pdf_file_path = os.path.join(Saving_folder, "All_Dop_Slices_Category_PieCharts_WithoutCB.pdf")
tiff_file_path = os.path.join(Saving_folder, "All_Dop_Slices_Category_PieCharts_WithoutCB.tiff")

# Ensure columns are strings
combined_df['Category'] = combined_df['Category'].astype(str)
combined_df['Regions'] = combined_df['Regions'].astype(str)

# Set the order of Regions
region_order = ['OGL', 'IGL', 'Med']
combined_df['Regions'] = pd.Categorical(combined_df['Regions'], categories=region_order, ordered=True)

# Count occurrences of each Category within each Region
data_counts = combined_df.groupby(['Regions', 'Category']).size().reset_index(name='Count')

# Calculate total counts for each Region
region_totals = data_counts.groupby('Regions')['Count'].sum()

# Calculate proportions within each Region
data_counts['Percent'] = data_counts.groupby('Regions')['Count'].transform(lambda x: (x / x.sum()) * 100)

# Define custom colors for the categories
category_colors = {
    'Excitation': '#7fbf7b',
    'Inhibition': '#af8dc3',
    'Non-Responding': '#7F7F7F'
}

# Start plotting
fig, axs = plt.subplots(1, len(region_order), figsize=(14, 5))

for i, region in enumerate(region_order):
    ax = axs[i]
    region_data = data_counts[data_counts['Regions'] == region]
    
    # Get labels, sizes, and colors for pie
    labels = region_data['Category']
    sizes = region_data['Count']
    colors = [category_colors[cat] for cat in labels]

    total_count = region_totals.get(region, 0)

    # Plot pie chart
    wedges, texts, autotexts = ax.pie(
        sizes,
        labels=labels,
        colors=colors,
        autopct='%1.1f%%',
        startangle=90,
        textprops={'fontsize': 10}
    )
    
    ax.set_title(f"{region} (n={total_count})", fontsize=12)

# Add central title
fig.suptitle('Dopamine: Cell Response Categories per Region', fontsize=14)

# Adjust layout
plt.tight_layout(rect=[0, 0, 1, 0.95])

# Save the figure
plt.savefig(pdf_file_path, format='pdf', bbox_inches='tight')
plt.savefig(tiff_file_path, format='tiff', dpi=300, bbox_inches='tight')

# Display
plt.show()


# %% [code]
# Initialize an empty list to hold dataframes
dataframes = []

#load all acetylcholine categorisation data

# Walk through the subfolders in the main folder
for root, dirs, files in os.walk(main_folder):
    # Check if the folder name contains the desired string
    if 'ACh' in root:
        for file in files:
            # Check if the file ends with _Categorization.xlsx
            if file.endswith('_Categorization.xlsx'):
                file_path = os.path.join(root, file)
                print(f"Loading file: {file_path}")  # Print the file being loaded
                
                # Extract the region name from the file name
                file_name = os.path.basename(file)
                parts = file_name.split('_')
                if len(parts) >= 3:  # Ensure there are enough parts to extract
                    region_name = parts[-2]
                    # Special case for 'left' or 'right'
                    if region_name.lower() in ['left', 'right'] and len(parts) >= 3:
                        region_name = parts[3]  # Take the string between the third and fourth underscores
                else:
                    region_name = 'Unknown'

                # Load the Excel file into a dataframe
                df = pd.read_excel(file_path)
                
                # Add the Regions column
                df['Regions'] = region_name
                
                # Append the dataframe to the list
                dataframes.append(df)

# Combine all the dataframes into one by rows
combined_df = pd.concat(dataframes, ignore_index=True)

# Display the combined dataframe
print(combined_df)


# %% [code]
# File names
pdf_file_path = os.path.join(Saving_folder, "All_ACh_Slices_Category_Bar.pdf")
tiff_file_path = os.path.join(Saving_folder, "All_ACh_Slices_Category_Bar.tiff")

# Ensure columns are strings
combined_df['Category'] = combined_df['Category'].astype(str)
combined_df['Regions'] = combined_df['Regions'].astype(str)

# Set the order of Regions
region_order = ['OGL', 'IGL', 'Med', 'CB']
#region_order = ['OGL', 'IGL', 'Med']
combined_df['Regions'] = pd.Categorical(combined_df['Regions'], categories=region_order, ordered=True)

# Count occurrences of each Category within each Region
data_counts = combined_df.groupby(['Regions', 'Category']).size().reset_index(name='Count')

# Calculate proportions within each Region
data_counts['Percent'] = data_counts.groupby('Regions')['Count'].transform(lambda x: (x / x.sum()) * 100)

# Calculate total counts for each Region
region_totals = data_counts.groupby('Regions')['Count'].sum()

# Pivot the data to prepare for plotting
plot_data = data_counts.pivot(index='Regions', columns='Category', values='Percent').fillna(0)

# Define custom colors for the categories
category_colors = {
    'Excitation': '#7fbf7b',
    'Inhibition': '#af8dc3',
    'Non-Responding': '#7F7F7F'
}

# Map the colors to the categories in the correct order
colors = [category_colors[cat] for cat in plot_data.columns]

# Plotting the data
fig, ax = plt.subplots(figsize=(10, 6))

# Adjust bar width and edge aesthetics
bars = plot_data.plot(
    kind='bar', 
    figsize=(10, 6), 
    width=0.8, 
    edgecolor='black', 
    color=colors,
    ax=ax
)

# Add grey grid lines
ax.grid(axis='y', color='grey', linestyle='--', linewidth=0.5, alpha=0.7, zorder=0)

# Ensure bars are above grid lines
for bar_group in bars.containers:
    for bar in bar_group:
        bar.set_zorder(3)

# Customize the plot
ax.set_title('Acetylcholine', fontsize=14)
ax.set_xlabel('', fontsize=12)
ax.set_ylabel('Percent of Cells', fontsize=12)
ax.set_xticks(range(len(region_order)))
ax.set_xticklabels(region_order, fontsize=10, rotation=0)
ax.tick_params(axis='y', labelsize=10)

# Add legend outside the plot
ax.legend(title='Category', fontsize=10, title_fontsize=12, loc='upper left', bbox_to_anchor=(1, 1))

# Add total counts below each region
for idx, region in enumerate(region_order):
    total_count = region_totals.get(region, 0)  # Get total count or 0 if region is missing
    ax.text(idx, -5, f'n={total_count}', ha='center', va='top', fontsize=10, transform=ax.transData)

# Adjust layout to make space for annotations
plt.tight_layout(rect=[0, 0.05, 1, 1])

# Save the plot as PDF and TIFF
plt.savefig(pdf_file_path, format='pdf', bbox_inches='tight')
plt.savefig(tiff_file_path, format='tiff', dpi=300, bbox_inches='tight')

# Display the plot
plt.show()


# %% [code]
# File names
pdf_file_path = os.path.join(Saving_folder, "All_ACh_Slices_Category_Bar_WithoutCB.pdf")
tiff_file_path = os.path.join(Saving_folder, "All_ACh_Slices_Category_Bar_WithoutCB.tiff")

# Ensure columns are strings
combined_df['Category'] = combined_df['Category'].astype(str)
combined_df['Regions'] = combined_df['Regions'].astype(str)

# Set the order of Regions
region_order = ['OGL', 'IGL', 'Med']
combined_df['Regions'] = pd.Categorical(combined_df['Regions'], categories=region_order, ordered=True)

# Count occurrences of each Category within each Region
data_counts = combined_df.groupby(['Regions', 'Category']).size().reset_index(name='Count')

# Calculate proportions within each Region
data_counts['Percent'] = data_counts.groupby('Regions')['Count'].transform(lambda x: (x / x.sum()) * 100)

# Calculate total counts for each Region
region_totals = data_counts.groupby('Regions')['Count'].sum()

# Pivot the data to prepare for plotting
plot_data = data_counts.pivot(index='Regions', columns='Category', values='Percent').fillna(0)

# Define custom colors for the categories
category_colors = {
    'Excitation': '#7fbf7b',
    'Inhibition': '#af8dc3',
    'Non-Responding': '#7F7F7F'
}

# Map the colors to the categories in the correct order
colors = [category_colors[cat] for cat in plot_data.columns]

# Plotting the data
fig, ax = plt.subplots(figsize=(10, 6))

# Adjust bar width and edge aesthetics
bars = plot_data.plot(
    kind='bar', 
    figsize=(10, 6), 
    width=0.8, 
    edgecolor='black', 
    color=colors,
    ax=ax
)

# Add grey grid lines
ax.grid(axis='y', color='grey', linestyle='--', linewidth=0.5, alpha=0.7, zorder=0)

# Ensure bars are above grid lines
for bar_group in bars.containers:
    for bar in bar_group:
        bar.set_zorder(3)

# Customize the plot
ax.set_title('Acetylcholine', fontsize=14)
ax.set_xlabel('', fontsize=12)
ax.set_ylabel('Percent of Cells', fontsize=12)
ax.set_xticks(range(len(region_order)))
ax.set_xticklabels(region_order, fontsize=10, rotation=0)
ax.tick_params(axis='y', labelsize=10)

# Add legend outside the plot
ax.legend(title='Category', fontsize=10, title_fontsize=12, loc='upper left', bbox_to_anchor=(1, 1))

# Add total counts below each region
for idx, region in enumerate(region_order):
    total_count = region_totals.get(region, 0)  # Get total count or 0 if region is missing
    ax.text(idx, -5, f'n={total_count}', ha='center', va='top', fontsize=10, transform=ax.transData)

# Adjust layout to make space for annotations
plt.tight_layout(rect=[0, 0.05, 1, 1])

# Save the plot as PDF and TIFF
plt.savefig(pdf_file_path, format='pdf', bbox_inches='tight')
plt.savefig(tiff_file_path, format='tiff', dpi=300, bbox_inches='tight')

# Display the plot
plt.show()


# %% [code]
# File names
pdf_file_path = os.path.join(Saving_folder, "All_ACh_Slices_Category_Pie_WithoutCB.pdf")
tiff_file_path = os.path.join(Saving_folder, "All_ACh_Slices_Category_Pie_WithoutCB.tiff")

# Ensure columns are strings
combined_df['Category'] = combined_df['Category'].astype(str)
combined_df['Regions'] = combined_df['Regions'].astype(str)

# Set the order of Regions
region_order = ['OGL', 'IGL', 'Med']
combined_df['Regions'] = pd.Categorical(combined_df['Regions'], categories=region_order, ordered=True)

# Count occurrences of each Category within each Region
data_counts = combined_df.groupby(['Regions', 'Category']).size().reset_index(name='Count')

# Calculate proportions within each Region
data_counts['Percent'] = data_counts.groupby('Regions')['Count'].transform(lambda x: (x / x.sum()) * 100)

# Define custom colors for the categories
category_colors = {
    'Excitation': '#7fbf7b',
    'Inhibition': '#af8dc3',
    'Non-Responding': '#7F7F7F'
}

# Prepare figure
fig, axes = plt.subplots(1, len(region_order), figsize=(15, 5))

# If only one axis (e.g. one region), wrap in list
if len(region_order) == 1:
    axes = [axes]

# Loop through each region and create pie chart
for i, region in enumerate(region_order):
    region_data = data_counts[data_counts['Regions'] == region]

    # Get labels and values
    labels = region_data['Category'].tolist()
    sizes = region_data['Percent'].tolist()
    counts = region_data['Count'].tolist()
    colors = [category_colors[label] for label in labels]

    # Plot pie chart
    wedges, texts, autotexts = axes[i].pie(
        sizes,
        labels=labels,
        autopct='%1.1f%%',
        colors=colors,
        startangle=90,
        counterclock=False,
        textprops={'fontsize': 10}
    )

    # Title and count annotation
    total_count = sum(counts)
    axes[i].set_title(f'{region}\nn={total_count}', fontsize=12)

# Overall title
fig.suptitle('Acetylcholine Response by Region', fontsize=16)

# Tight layout to avoid overlap
plt.tight_layout(rect=[0, 0, 1, 0.95])

# Save the plot
plt.savefig(pdf_file_path, format='pdf', bbox_inches='tight')
plt.savefig(tiff_file_path, format='tiff', dpi=300, bbox_inches='tight')

# Show plot
plt.show()


# %% [code]
# Ensure the folder exists
if not os.path.exists(main_folder):
    print(f"Error: The folder '{main_folder}' does not exist.")
else:
    # List all folders inside main_folder that contain ' Dop' in their name
    matching_folders = [os.path.join(main_folder, f) for f in os.listdir(main_folder) 
                        if os.path.isdir(os.path.join(main_folder, f)) and 'Dop' in f]

    # Display the matching folders
    if matching_folders:
        print("Folders containing 'Dop':")
        for folder in matching_folders:
            print(folder)
    else:
        print("No folders containing 'Dop' found.")


# %% [markdown]
dop slope vs Cell Pose radius - all data

# %% [code]
# Initialize DataFrames to store combined data
lig_Slope_combined = pd.DataFrame()

# Iterate through folders
for folder in matching_folders:
    for file in os.listdir(folder):
        file_path = os.path.join(folder, file)
        if file.endswith('_Lig_Slope.xlsx'):
            df = pd.read_excel(file_path)
            df['Folder'] = file.replace('_Lig_Slope.xlsx', '')  # Add Folder column without suffix
            lig_Slope_combined = pd.concat([lig_Slope_combined, df], ignore_index=True)

# Output results
print(f"Combined {len(lig_Slope_combined)} rows of Lig Slope data.")


# %% [code]
# File names
pdf_file_path = os.path.join(Saving_folder, f"All_Dop_Slope_Histogram.pdf")
tiff_file_path = os.path.join(Saving_folder, f"All_Dop_Slope_Histogram.tiff")

# Plot the histogram
plt.figure(figsize=(8, 6))
plt.hist(lig_Slope_combined['Slope'], bins=30, edgecolor='black', alpha=0.7)

# Add labels and title
plt.xlabel('Ligand Slope', fontsize=14)
plt.ylabel('Frequency', fontsize=14)
plt.title('Histogram of Ligand Slope', fontsize=16)

# Show the plot
plt.grid(axis='y', linestyle='--', alpha=0.7)

# Save the plot
plt.savefig(pdf_file_path, format='pdf')
plt.savefig(tiff_file_path, format='tiff', dpi=300)

plt.show()


# %% [code]
region_colors = {'OGL': '#8da0cb', 'IGL': '#66c2a5', 'Med': '#fc8d62', 'CB': '#e78ac3'}
desired_order = ['OGL', 'IGL', 'Med','CB']

# File names
pdf_file_path = os.path.join(Saving_folder, f"All_Dop_Slope_Violin_perRegion.pdf")
tiff_file_path = os.path.join(Saving_folder, f"All_Dop_Slope_Violin_perRegion.tiff")

# Create the figure
plt.figure(figsize=(8, 6))

# Violin plot (distribution) - Background layer
sns.violinplot(
    data=lig_Slope_combined,
    x="Regions",
    y="Slope",
    order=desired_order,
    palette=region_colors,
    inner=None,  # Hide inner lines to prevent overlap with strip plot
    alpha=0.5  # Transparency for visibility
)

# Strip plot (individual data points) - Middle layer
sns.stripplot(
    data=lig_Slope_combined,
    x="Regions",
    y="Slope",
    order=desired_order,
    palette=region_colors,
    jitter=True,
    size=8,
    alpha=0.7
)

# Point plot (mean values) - Top layer
sns.pointplot(
    data=lig_Slope_combined,
    x="Regions",
    y="Slope",
    order=desired_order,
    color="grey",  # Mean values in grey
    markers="D",  # Diamond marker for mean
    scale=1.2,  # Slightly larger marker
    errwidth=1.5,  # Error bar width
    capsize=0.2,  # Small caps on error bars
    join=False,
    estimator=np.mean,
    zorder=3  # Ensure point plot is on top
)

# Set y-axis limits from 0.001 to 0.3
plt.ylim(-0.03, 0.4)

# Customizing the plot
plt.title("Slopes with Mean Values", fontsize=14)
plt.xlabel("", fontsize=12)
plt.ylabel("Slope", fontsize=12)
plt.xticks(fontsize=10)
plt.yticks(fontsize=10)
plt.grid(axis='y', linestyle='--', alpha=0.6)

# Add legend
plt.legend(loc="upper right", fontsize=10)

# Adjust layout
plt.tight_layout()

# Save the plot
plt.savefig(pdf_file_path, format='pdf')
plt.savefig(tiff_file_path, format='tiff', dpi=300)

# Show the plot
# plt.show()


# %% [code]
region_colors = {'OGL': '#8da0cb', 'IGL': '#66c2a5', 'Med': '#fc8d62', 'CB': '#e78ac3'}
desired_order = ['OGL', 'IGL', 'Med']

# File names
pdf_file_path = os.path.join(Saving_folder, f"All_Dop_Slope_Violin_perRegion_WithoutCB.pdf")
tiff_file_path = os.path.join(Saving_folder, f"All_Dop_Slope_Violin_perRegion_WithoutCB.tiff")

# Create the figure
plt.figure(figsize=(8, 6))

# Violin plot (distribution) - Background layer
sns.violinplot(
    data=lig_Slope_combined,
    x="Regions",
    y="Slope",
    order=desired_order,
    palette=region_colors,
    inner=None,  # Hide inner lines to prevent overlap with strip plot
    alpha=0.5  # Transparency for visibility
)

# Strip plot (individual data points) - Middle layer
sns.stripplot(
    data=lig_Slope_combined,
    x="Regions",
    y="Slope",
    order=desired_order,
    palette=region_colors,
    jitter=True,
    size=8,
    alpha=0.7
)

# Point plot (mean values) - Top layer
sns.pointplot(
    data=lig_Slope_combined,
    x="Regions",
    y="Slope",
    order=desired_order,
    color="grey",  # Mean values in grey
    markers="D",  # Diamond marker for mean
    scale=1.2,  # Slightly larger marker
    errwidth=1.5,  # Error bar width
    capsize=0.2,  # Small caps on error bars
    join=False,
    estimator=np.mean,
    zorder=3  # Ensure point plot is on top
)

# Set y-axis limits from 0.001 to 0.3
plt.ylim(-0.03, 0.4)

# Customizing the plot
plt.title("Slopes with Mean Values", fontsize=14)
plt.xlabel("", fontsize=12)
plt.ylabel("Slope", fontsize=12)
plt.xticks(fontsize=10)
plt.yticks(fontsize=10)
plt.grid(axis='y', linestyle='--', alpha=0.6)

# Add legend
plt.legend(loc="upper right", fontsize=10)

# Add a black dotted line at slope = 0.08
plt.axhline(y=0.08, color='black', linestyle='dotted', linewidth=1.5)

# Adjust layout
plt.tight_layout()

# Save the plot
plt.savefig(pdf_file_path, format='pdf')
plt.savefig(tiff_file_path, format='tiff', dpi=300)

# Show the plot
# plt.show()


# %% [code]
# Extract the lowest folder name from base_folder
lowest_folder = os.path.basename(os.path.normpath(main_folder))
region_colors = {'OGL': '#8da0cb', 'IGL': '#66c2a5', 'Med': '#fc8d62'}
desired_order = ['OGL', 'IGL', 'Med']

# File names
pdf_file_path = os.path.join(Saving_folder, f"All_Dop_Slope_ECDF_perRegion.pdf")
tiff_file_path = os.path.join(Saving_folder, f"All_Dop_Slope_ECDF_perRegion.tiff")

# Create the figure
plt.figure(figsize=(8, 6))

# Plot ECDFs for each region
for region in desired_order:
    subset = lig_Slope_combined[lig_Slope_combined["Regions"] == region]
    sns.ecdfplot(
        data=subset, 
        x="Slope", 
        label=region, 
        color=region_colors[region], 
        linewidth=2
    )

# Customizing the plot
plt.title("Cumulative Distribution of Slopes", fontsize=14)
plt.xlabel("Slope", fontsize=12)
plt.ylabel("Cumulative Probability", fontsize=12)
plt.xticks(fontsize=10)
plt.yticks(fontsize=10)
plt.grid(axis='both', linestyle='--', alpha=0.6)

# Add legend
plt.legend(title="Regions", fontsize=10, loc="lower right")

# Adjust layout
plt.tight_layout()

# Save the plot
plt.savefig(pdf_file_path, format='pdf')
plt.savefig(tiff_file_path, format='tiff', dpi=300)

# Show the plot
plt.show()


# %% [code]
# Extract the lowest folder name from base_folder
lowest_folder = os.path.basename(os.path.normpath(main_folder))
region_colors = {'OGL': '#8da0cb', 'IGL': '#66c2a5', 'Med': '#fc8d62'}
desired_order = ['OGL', 'IGL', 'Med']

# File names
pdf_file_path = os.path.join(Saving_folder, f"All_Dop_Slope_KDE_perRegion.pdf")
tiff_file_path = os.path.join(Saving_folder, f"All_Dop_Slope_KDE_perRegion.tiff")

# Create the figure
plt.figure(figsize=(8, 6))

# Plot KDE for each region
for region in desired_order:
    subset = lig_Slope_combined[lig_Slope_combined["Regions"] == region]
    sns.kdeplot(
        data=subset, 
        x="Slope", 
        label=region, 
        color=region_colors[region], 
        fill=True,  # Fill under the KDE curve
        alpha=0.4,  # Transparency for better visualization
        linewidth=2
    )

# Customizing the plot
plt.title("Kernel Density Estimate of Slopes", fontsize=14)
plt.xlabel("Slope", fontsize=12)
plt.ylabel("Density", fontsize=12)
plt.xticks(fontsize=10)
plt.yticks(fontsize=10)
plt.grid(axis='both', linestyle='--', alpha=0.6)

# Add legend
plt.legend(title="Regions", fontsize=10, loc="upper right")

# Adjust layout
plt.tight_layout()

# Save the plot
plt.savefig(pdf_file_path, format='pdf')
plt.savefig(tiff_file_path, format='tiff', dpi=300)

# Show the plot
plt.show()


# %% [code]
#stats with CB
# Check Normality (Shapiro-Wilk test for each region)
normality_results = lig_Slope_combined.groupby("Regions")["Slope"].apply(lambda x: stats.shapiro(x)[1])  # p-values
print("Normality Test (Shapiro-Wilk):\n", normality_results)

# Check Homogeneity of Variance (Levene's test)
levene_p = stats.levene(*[group["Slope"].values for name, group in lig_Slope_combined.groupby("Regions")])[1]
print(f"Levene’s test for equality of variances: p = {levene_p}")

# One-way ANOVA if normality holds and variances are equal
if all(p > 0.05 for p in normality_results) and levene_p > 0.05:
    anova_result = stats.f_oneway(*[group["Slope"].values for name, group in lig_Slope_combined.groupby("Regions")])
    print("One-way ANOVA:\n", anova_result)

    # Post-hoc test: Tukey's HSD
    tukey = mc.pairwise_tukeyhsd(lig_Slope_combined["Slope"], lig_Slope_combined["Regions"])
    print("Tukey's HSD Results:\n", tukey)

# Welch’s ANOVA if variances are unequal
elif levene_p < 0.05:
    welch_anova = pg.welch_anova(dv="Slope", between="Regions", data=lig_Slope_combined)
    print("Welch’s ANOVA:\n", welch_anova)

    # Games-Howell Post-hoc test
    games_howell = pg.pairwise_gameshowell(dv="Slope", between="Regions", data=lig_Slope_combined)
    print("Games-Howell Post-hoc Test:\n", games_howell)

# Kruskal-Wallis test if normality is violated
else:
    kw_result = stats.kruskal(*[group["Slope"].values for name, group in lig_Slope_combined.groupby("Regions")])
    print("Kruskal-Wallis Test:\n", kw_result)

    # Dunn’s Post-hoc test
    dunn_test = pg.pairwise_tests(dv="Slope", between="Regions", data=lig_Slope_combined, padjust="bonferroni")
    print("Dunn's Test Results:\n", dunn_test)


# %% [code]
# Categorizing slopes
lig_Slope_combined['Slope_Category'] = lig_Slope_combined['Slope'].apply(lambda x: 'Steep Slope' if x > 0.08 else 'Shallow Slope')

# %% [code]
desired_order = ['OGL', 'IGL', 'Med']
# File names
pdf_file_path = os.path.join(Saving_folder, f"All_Dop_Slope_Category_perRegion_withoutCB.pdf")
tiff_file_path = os.path.join(Saving_folder, f"All_Dop_Slope_Category_perRegion_withoutCB.tiff")

# Define response order
slope_cat_order = ['Steep Slope', 'Shallow Slope']

# Define colors for categories
category_colors = {
    'Shallow Slope': '#1f78b4', 
    'Steep Slope': '#33a02c' 
}

# Count occurrences of each Slope_Category per Region
category_counts = lig_Slope_combined.groupby(['Regions', 'Slope_Category']).size().unstack(fill_value=0)

# Convert counts to proportions
category_proportions = category_counts.div(category_counts.sum(axis=1), axis=0)

# Reorder index based on desired order
category_proportions = category_proportions.reindex(desired_order)

# Reorder columns based on slope category order
category_proportions = category_proportions.reindex(columns=slope_cat_order, fill_value=0)

# Create a grouped bar plot to show proportions side by side with custom colors
ax = category_proportions.plot(kind='bar', stacked=False, figsize=(8, 6), color=[category_colors[col] for col in category_proportions.columns])

plt.ylabel("Proportion")
plt.xlabel("")
plt.title("Proportion of Slope Categories per Region")
plt.legend(title="Slope Category")
plt.grid(axis='y', linestyle='--', alpha=0.7)

# Save the plot
plt.savefig(pdf_file_path, format='pdf')
plt.savefig(tiff_file_path, format='tiff', dpi=300)

plt.show()


# %% [code]
desired_order = ['OGL', 'IGL', 'Med','CB']
# File names
pdf_file_path = os.path.join(Saving_folder, f"All_Dop_Slope_Category_perRegion.pdf")
tiff_file_path = os.path.join(Saving_folder, f"All_Dop_Slope_Category_perRegion.tiff")

# Define response order
slope_cat_order = ['Steep Slope', 'Shallow Slope']

# Define colors for categories
category_colors = {
    'Shallow Slope': '#1f78b4', 
    'Steep Slope': '#33a02c' 
}

# Count occurrences of each Slope_Category per Region
category_counts = lig_Slope_combined.groupby(['Regions', 'Slope_Category']).size().unstack(fill_value=0)

# Convert counts to proportions
category_proportions = category_counts.div(category_counts.sum(axis=1), axis=0)

# Reorder index based on desired order
category_proportions = category_proportions.reindex(desired_order)

# Reorder columns based on slope category order
category_proportions = category_proportions.reindex(columns=slope_cat_order, fill_value=0)

# Create a grouped bar plot to show proportions side by side with custom colors
ax = category_proportions.plot(kind='bar', stacked=False, figsize=(8, 6), color=[category_colors[col] for col in category_proportions.columns])

plt.ylabel("Proportion")
plt.xlabel("")
plt.title("Proportion of Slope Categories per Region")
plt.legend(title="Slope Category")
plt.grid(axis='y', linestyle='--', alpha=0.7)

# Save the plot
plt.savefig(pdf_file_path, format='pdf')
plt.savefig(tiff_file_path, format='tiff', dpi=300)

plt.show()
```
